# Single-molecule spatial genomics reveals the multi-scale organization and plasticity of extrachromosomal DNA in glioblastoma

**DOI:** 10.64898/2026.03.05.709911

**Authors:** Brett Taylor, Weixiu Dong, Tula Keal, Zhaoning Wang, Bharath Saravanan, Zane A. Gibbs, Yohei Miyake, Daisuke Kawauchi, Raghavendra Vadla, Abhinaba Banerjee, Omar Elkassih, Sahana Kashyap, Brandon M. Jones, Kseniya Malukhina, Mahsa Nafisi, Timothy H. Loe, Joseph Bendik, Tom McCallister, Yang Xie, Lei Chang, Clark Chen, Bing Ren, Frank Furnari, Bogdan Bintu

## Abstract

Extrachromosomal DNA (ecDNA) is a major driver of intratumoral heterogeneity and is associated with poor clinical outcomes across cancers, yet how individual ecDNA molecules are organized and regulated within intact tumors remains unknown. Here, we leveraged single-molecule, multi-modal spatial genomics to resolve the three-dimensional chromatin organization and transcriptional activity of individual *EGFR*-containing ecDNA molecules in glioblastoma (GBM) cells *in vitro*, in orthotopic xenografts, and in patient-derived GBM tissue. At the larger scale, we find that distinct GBM molecular and functional states emerge depending on the local cellular environment. *EGFR* expression was markedly different between GBM subpopulations, and perturbations of *EGFR* dosage shifted GBM cellular states. ecDNA expression was modulated by multiple mechanisms, including variation in copy number, chromatin organization, DNA sequence, and chromosomal reintegration, which were simultaneously measured within the same cells. At the single-molecule scale, ecDNA adopts a physically expanded chromatin configuration with larger ecDNA molecules having higher transcriptional activity and interaction with active transcriptional machinery. ecDNA regulation was coordinated within cells and across GBM states, and ecDNA copy number, structure, and transcription were spatially organized across the tumor architecture. Co-culturing GBM cells with neurons recapitulated key features of infiltrative regions, including lower *EGFR* expression, reduced ecDNA copy number, and increased chromosomal reintegration, suggesting a causal role for the microenvironment in shaping ecDNA regulation. Collectively, these findings support a model in which GBM states and ecDNA are linked, plastic, and influenced by microenvironmental contexts, revealing a previously inaccessible layer of genome organization underlying tumor heterogeneity and malignant cell behavior.

## Introduction

Every tumor is a complex cellular ecosystem whose organization and evolutionary dynamics influence disease progression and therapeutic response:^1–5^ “Perhaps cancer cells, like organs and organisms, should also be imagined as a community…a cooperative assembly…an ecology gone wrong,” writes physician-scientist Siddhartha Mukherjee.^6^ A central feature of this complexity is intratumoral heterogeneity, which enables cancer cells to adapt to environmental pressures and evade targeted therapies.^3–5,7^ In recent years, extrachromosomal DNA (ecDNA), a class of focally amplified circular DNA elements that harbor oncogenes and enhancers, has emerged as a key driver of genetic heterogeneity, underscored by its recent designation as a “Cancer Grand Challenge.”^8–10^ Lacking centromeres, ecDNA molecules are randomly segregated into daughter cells, promoting cell-to-cell variation in copy number and oncogene dosage.^8,11^ Consistent with this behavior, the presence of ecDNA is strongly associated with elevated oncogene expression, cellular adaptation, and poor clinical outcomes.^9,11–14^

Beyond driving copy number variation, ecDNA exhibits regulatory properties that distinguish it from chromosomal amplifications. Oncogenes encoded on ecDNA are among the most highly transcribed genes in cancer^15^ owing to both copy number-dependent and copy number-independent mechanisms.^16^ While ecDNA copy number, genetic composition, and re-integration into chromosomes have previously been shown to vary across different tumor regions and metastatic lesions,^17,18^ these properties have not been directly measured within transcriptionally defined malignant cell states, microenvironmental contexts, or the broader tumor organization. Existing sequencing approaches average ecDNA signals across or within cells, while traditional imaging approaches lack the multimodal throughput to resolve ecDNA structure, its transcriptional activity, and cell type/state identity of the containing cells.^13,15,19–21^ Multi-modal Multiplexed Error-Robust Fluorescence *in situ* Hybridization (mm-MERFISH) simultaneously captures DNA, RNA, and protein, enabling the investigation of chromatin organization and regulation at single-molecule resolution in its native spatial and functional contexts;^22^ however, this technology has not been used to study copy number aberrations and complex structural variants like ecDNA.^22–29^

Glioblastoma (GBM), the most common and lethal primary brain tumor in adults, provides a compelling system in which to address these questions.^30^ Historically called glioblastoma multiforme (“many forms”) based on the histological variation observed in primary specimens,^31^ GBM tumors consist of cells spanning distinct, interconverting cellular states,^32–37^ such as the four defined in a landmark GBM study^33^: astrocyte-like (AC-like), neural progenitor cell-like (NPC-like), oligodendrocyte precursor cell-like (OPC-like), and mesenchymal-like (MES-like). GBM harbors ecDNA more frequently than any other tumor type (57-86%), with EGFR-amplifying ecDNA (ecEGFR) as the most common species (33-63%).^9,17^ We previously showed that ecEGFR copy number varied across states and that treatment with an EGFR inhibitor shifted GBM states *in vitro*, thereby linking ecDNA to GBM transcriptional heterogeneity.^19^ However, how individual ecDNA molecules are regulated across scales and how they interact with cellular states and the microenvironment remain largely unexplored.

Here, we extended mm-MERFISH to amplified circular genomes to directly visualize more than a hundred thousand individual EGFR-containing ecDNA molecules in glioblastoma cells *in vitro*, in orthotopic xenograft models, and in patient-derived GBM tissue. We tiled ecEGFR amplicons (>1-Mb; chr7: 54.9-56.2-Mb) at 5-Kb resolution and developed ecDNATracer to reconstruct individual ecEGFR molecules *in situ*. In addition to chromatin tracing single ecEGFR molecules, we imaged nascent transcription of the coding genes on ecEGFR and proteins associated with transcriptional regulation within the same cells. After characterizing ecEGFR molecules *in vitro*, GBM cells orthotopically engrafted into mouse brains were imaged to determine how ecEGFR molecules are shaped by the microenvironment *in vivo*. Finally, we applied mm-MERFISH to patient-derived GBM tissue and, together with reanalysis of spatial transcriptomic data from 26 GBM samples, evaluated how ecEGFR phenotypes spatially map onto GBM cellular states and non-malignant cell populations within tumors. With functional experiments, we establish that modulating EGFR expression shifted GBM cellular states, and that changing the microenvironmental context by co-culturing GBM cells with neurons reduced ecDNA copy number and increased its chromosomal reintegration. Collectively, these findings establish the heterogeneity and plasticity of ecDNA, which integrates intrinsic chromatin regulation with extrinsic ecological pressures, revealing how the structural and functional diversity of ecDNA underlies cell state heterogeneity, malignant cell behavior, and the broader tumor architecture.

## Results

### Chromatin tracing resolves the three-dimensional organization of individual ecDNA molecules

To directly investigate the single-molecule 3D structure and activity of ecEGFR in GBM, we applied mm-MERFISH to the well-characterized 1.2-Mb ecEGFR amplicon (chr7:54.9 - 56.2 Mb) in the cell line GBM39.^15,38^ To reconstruct the entire structure in 3D, probes were designed to sequentially target 5-Kb regions of the ecEGFR amplicon. Primary probes were first hybridized to fixed GBM39 cells in interphase, and 252 consecutive 5-Kb regions were sequentially read out across 84 hybridization cycles using 3-color imaging (**Fig.1A-B**). To distinguish ecEGFR from its chromosomal counterpart, three additional loci were imaged: two targeting 50-Kb segments immediately downstream of the ecEGFR end (chr7: 56.20 - 56.25 Mb, chr7: 56.25 - 56.3-Mb) and another targeting the 27-Kb EGFRvIII deletion––exons 2-7––on ecEGFR (chr7: 55.287 - 55.315-Mb) (**Fig.1A-B**). These marked loci distinguished traces corresponding to ecEGFR, which do not colocalize with these signals, from traces corresponding to the endogenous chromosomal EGFR loci (**Fig.1A-B**).

**Figure 1.**
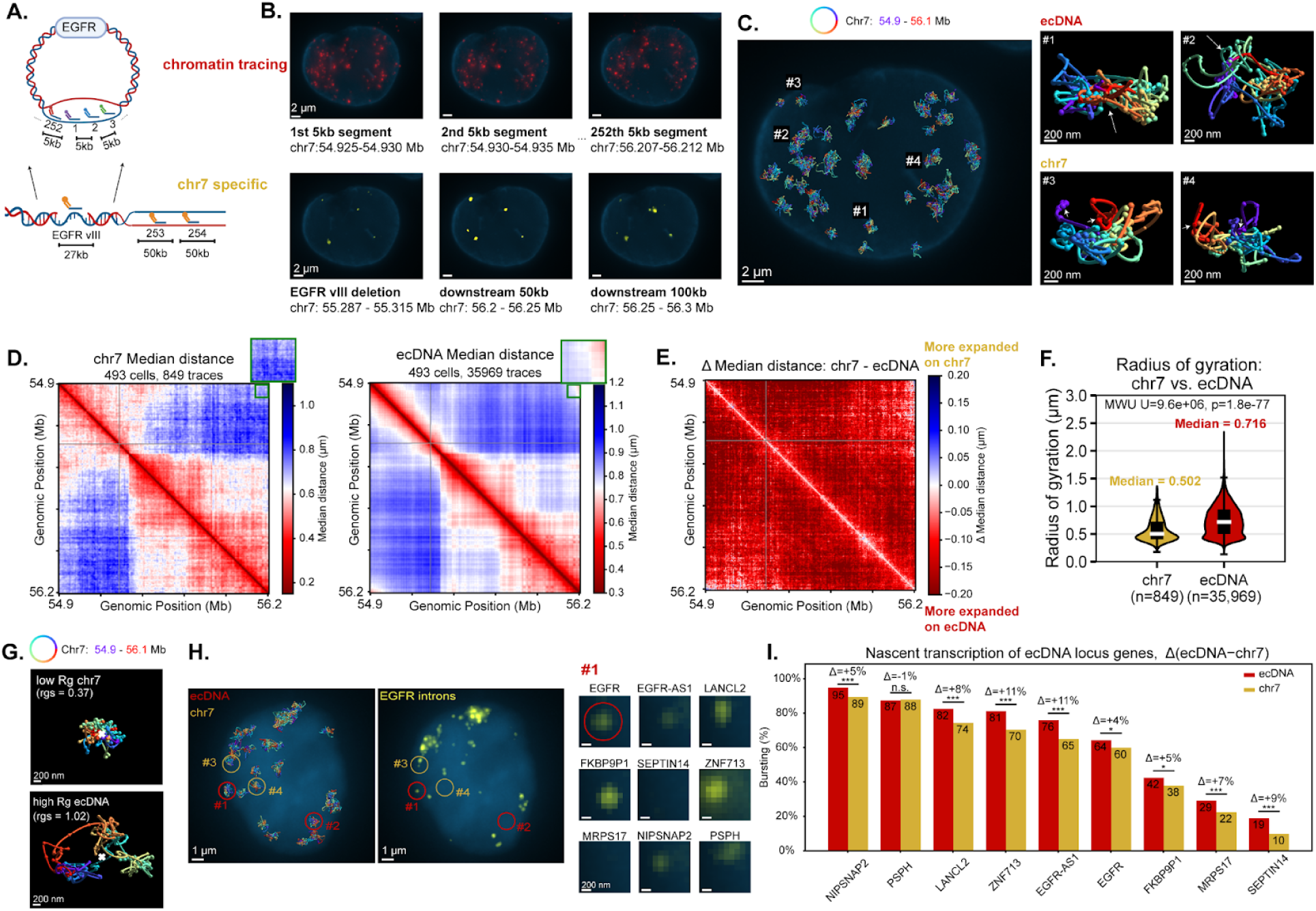
mm-MERFISH resolves the single-molecule chromatin structure and transcriptional activity of ecDNA and chromosomes. **(A.)** Schematic of labeling strategy highlighting: 1) chromatin tracing probes (red) targeting both ecDNA and EGFR locus in chr 7 and 2) chr 7 specific probes (yellow). **(B.)** Top row: Representative images of 5-Kb segments of the EGFR ecDNA and chr 7 loci. 84 3-color cycles are used to image 252 5-Kb segments spanning the entire ∼1.2-Mb EGFR ecDNA. Bottom row: Representative images of chr 7 specific probes targeting two downstream loci (50- and 100-Kb beyond the ecDNA) and the EGFRvIII region that is deleted on ecEGFR. Blue - lamina stain. **(C.)** 3D traces of ecDNA and chr7 EGFR in a representative cell. White arrows mark the first and last 5-Kb segments. **(D.)** Median distance matrices of the chr7 EGFR locus (left) and ecDNA (right). **(E.)** Difference in median physical distance between chr7 EGFR locus and ecDNA. **(F.)** Violin plots showing the radius of gyration of chr7 EGFR locus (orange) and ecDNA (red). A two-sided Mann-Whitney U test was used to test for statistical significance. **(G.)** Representative 3D traces of chr7 EGFR locus (top) and ecDNA (bottom) with different radii of gyration. White cross marks the center of mass for each trace. **(H.)** Traces of ecDNA and chr 7 EGFR locus (left) and images of EGFR nascent transcription (right) for the same representative cell. Red circles mark representative ecDNA traces, while orange circles mark chr7 traces. Each inset represents nascent RNA for each of the 9 genes on the single ecDNA molecule labeled by #1. **(I.)** Paired bar graph of nascent transcription rate for the genes on ecEGFR (red) versus chr7 (orange). Genes are ordered by their transcription rate. ***: p < 1e-4, **: p < 1e-3, *: p < 1e-2, n.s.: not significant by two-sided Fisher’s exact test. *N* = 849 chr7 traces and 35,969 ecDNA traces.

With the fitted 3D positions of these 252 5-Kb genomic loci, we adapted our previous 3D tracing tools^39^ with a new spatial genome aligner called ecDNATracer, facilitating the 3D reconstruction of the ∼1.2-Mb EGFR locus from within ecEGFR and chromosome 7 (**Extended Data Fig.1A**). Briefly, for each ecDNA molecule we considered multiple ways to connect the candidate 5-Kb fluorescent loci detected across imaging cycles and selected the configuration best constrained by polymer physics. As in prior work, spatial consistency was evaluated using a Gaussian chain model, in which the expected three-dimensional separation between loci scales with their genomic distance.^39,40^ However, rather than relying solely on spatial proximity, our framework also incorporates fluorescence brightness as an independent measure of signal reliability, jointly weighting physical plausibility and signal intensity when reconstructing each trace. In ecDNATracer, we further extended this approach to explicitly accommodate circular DNA topology, enabling reconstruction of ecDNA molecules (Methods). In sum, we imaged a total of 655 cells across 2 experiments capturing 48,460 ecDNA traces and 1,063 endogenous chromosomal EGFR loci at 5-Kb resolution and reconstructed their configuration in 3D (**Fig.1C, Extended Fig.2A,B**).

**Figure 2.**
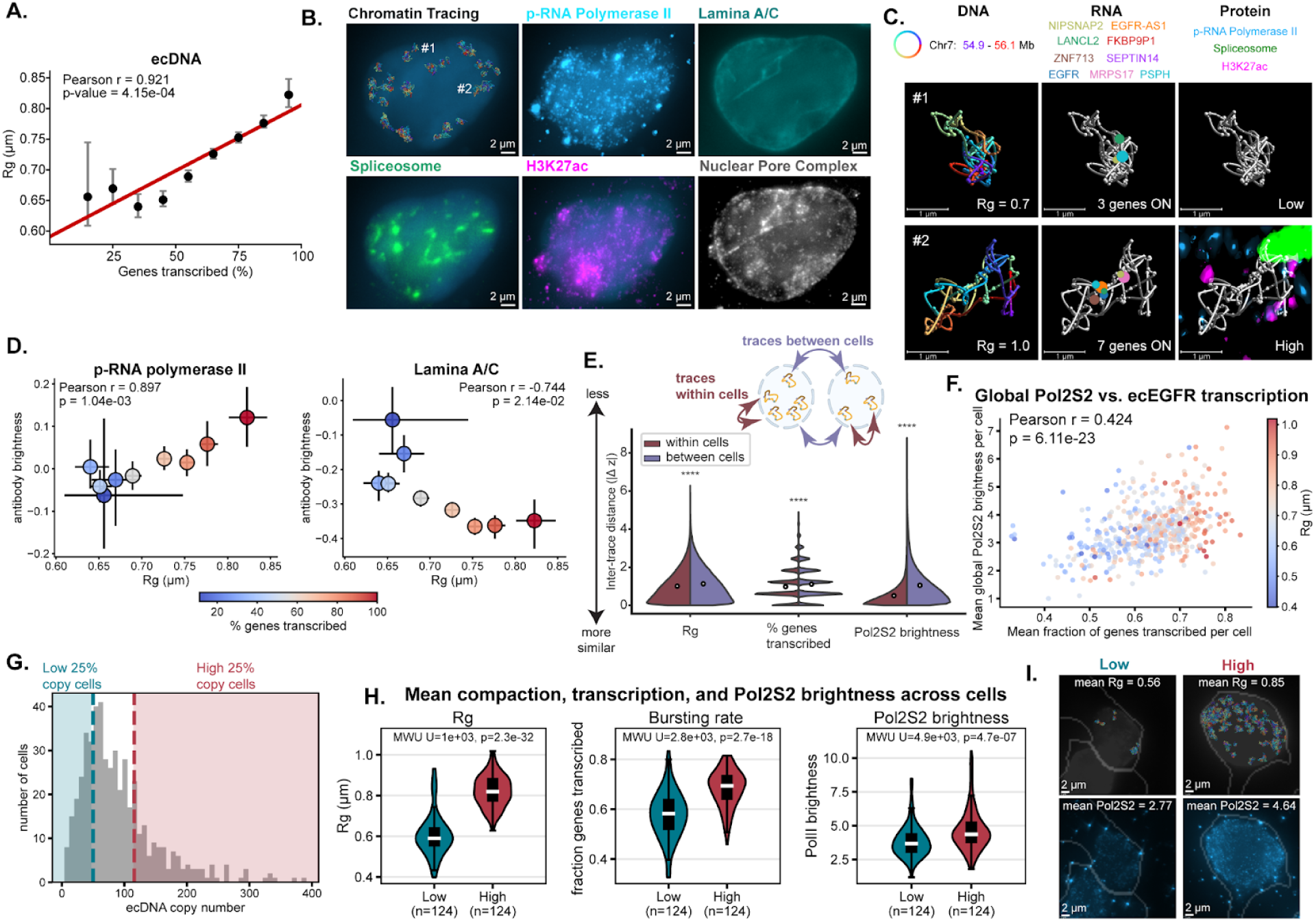
Coordinated physical expansion, transcriptional activity, and deposition of transcriptional machinery underlie heterogeneity across single ecDNA molecules. **(A.)** Stratified scatter plot of the percentage of genes transcribed and the radius of gyration (Rg) for ecDNA. Each black dot represents the median per group, and the bar represents the 95% confidence interval. The red line marks the ordinary least squares line. N = 34, 349 ecDNA traces total. From left to right, each group has N = [248, 754, 1986, 3936, 6598, 8341, 7203, 4069, 1214] traces. **(B.)** Images of a representative cell showing ecDNA traces and immunostaining by mm-MERFISH of phosphorylated-RNA Polymerase II (Pol2S2), spliceosome (SC35), H3K27ac, Lamina A/C, and Nuclear Pore Complex (NUP98). **(C.)** Representative low Rg (top row) and high Rg (bottom row) single molecule traces in the context of multi-modal measurements. **(D.)** Correlation between radius of gyration and the normalized Pol2S2 brightness across ecDNA traces with different percentages of transcribed genes. Dots mark medians and error bars mark 95% confidence intervals. N is the same as A). **(E.)** Split violin plot comparisons of z scored differences across ecDNA from within the same cell vs. from different cells. N = 438,277 simulations. White circles mark medians. A two-sided Mann-Whitney U test was used to compute statistical significance. **** - p < 1e-4. **(F.)** Scatter plot comparing the mean transcription and global Pol2S2 deposition per cell, colored by the mean compaction per cell. N = 493 cells. **(G.)** Histogram of ecDNA copy number distribution. N = 493 cells. **(H.)** Violins comparing the mean transcription, compaction and Pol2S2 deposition between the low copy and high copy cells. A one-sided Mann-Whitney U test was used to compute statistical significance. **(I.)** Representative images of cells with low copy (left column) and high copy (right column). Top row: Single molecules represented by the rainbow configuration. Nucleus represented by DAPI stain (gray). Bottom row: immunofluorescence image of Pol2S2 (blue). Gray contour marks the cytoplasm boundaries.

To ensure the reliability and accuracy of ecDNATracer, we compared our measured traces to simulated polymer structures and to population average ecDNA structure from sequencing-based assays. Specifically, we applied ecDNATracer to simulated linear (chromosomal) and circular (ecDNA) polymers with and without noise and compared the results with measured ecEGFR 3D structures (**Extended Data Fig.1B-D**). Although circular, ecEGFR molecules structurally do not resemble simple ring-like structures due to the high degree of chromatin folding in both simulated and experimental data at 5-Kb resolution (**Extended Data Fig.1B-D**).^23,24,39^ To investigate the population average structure of ecDNA, we calculated the median spatial distance between each pair of 5-Kb loci (**Extended Data Fig.2C**). This average representation facilitated comparisons with an orthogonal approach that measures contact frequency between chromatin loci using proximity ligation-based sequencing (e.g., Hi-C). The inter-loci distance revealed chromatin domains and “loop” features that correlated with HiC contact probability in GBM39^19^ (Pearson correlation coefficient = 0.75, **Extended Data Fig.2C-E**). The inter-loci distance and TAD-like pattern was reproducible across technical replicates (**Extended Data Fig.2F-G**).

Together, these results establish that single ecDNA molecules can be directly visualized and reconstructed in three dimensions within individual cells, revealing organized chromatin structures. This capability enables direct interrogation of the relationship between ecDNA chromatin organization and transcriptional activity.

### ecEGFR is more physically expanded and transcriptionally active than its chromosomal counterpart

We compared the 3D organization of the EGFR locus on ecDNA and its chromosomal counterpart by measuring the average distance between the constituent 5-Kb segments (**Fig.1D, Extended Data Fig.2H**). On average, the two groups have similar 3D conformation features, including matching domain structures (**Fig.1D, Extended Data Fig.2H**), suggesting that similar architectural proteins are binding both configurations. A notable exception is the enriched proximity between the beginning and end of ecEGFR (**Fig.1D**, top-right corner insets), supporting its circular topology. Despite these similar features, we found that ecEGFR was more physically expanded with larger interdistances compared to matched chromosomes (**Fig.1E, Extended Data Fig.2I**). Relative expansion of ecDNA was further supported by a larger average radius of gyration (Rg),^25,41^ defined as the average distance of the 5-Kb loci from the centroid of the chromatin fiber (**Fig.1F, Extended Data Fig.2J**). A trace with low Rg (**Fig.1G, top**) will be more condensed towards the centroid, while a trace that has high Rg will be more expanded (**Fig.1G, bottom**). This result defies expectations of simulated polymer physics models which predict linear polymers to occupy more volume than circular polymers, suggesting that non-polymeric mechanisms mediate ecEGFR expansion (**Extended Data Fig.1E**).

To further explore the interplay between ecEGFR copy number, structure, and transcription, we designed and co-imaged multiplexed probes that target the intronic and exonic regions of the genes that are on the 1.2-Mb amplicon. Specifically, we measured nascent RNA for EGFR and eight additional genes present within the 1.2-Mb *EGFR* locus and compared transcriptional bursting on individual ecEGFR and chromosome 7 molecules (**Extended Data Fig.3A**). Consistent with transcriptional bursting of RNA polymerase,^22,42^ only a subset of DNA traces were actively transcribing (**Fig.1H**). We assessed bursting on a single-molecule basis for each gene (**Fig.1H**) and quantified gene-specific bursting rates, validated by their corresponding exon abundance (**Fig.1I, Extended Data Fig.3B**). Consistent with their overall structural similarity, the transcriptional activity of genes on ecEGFR and chromosomes was highly correlated (Pearson’s correlation coefficient 0.99, p-value = 1.67e-07), suggesting that genes on ecDNA largely retain the regulatory features of their chromosomal counterparts. Notably, however, most genes (8/9) exhibited a modest but significantly higher bursting frequency on ecEGFR compared with the chromosomal loci (**Fig. 1I**), indicating that ecEGFR molecules are more transcriptionally active than their chromosomal counterparts at single-molecule resolution.

**Figure 3.**
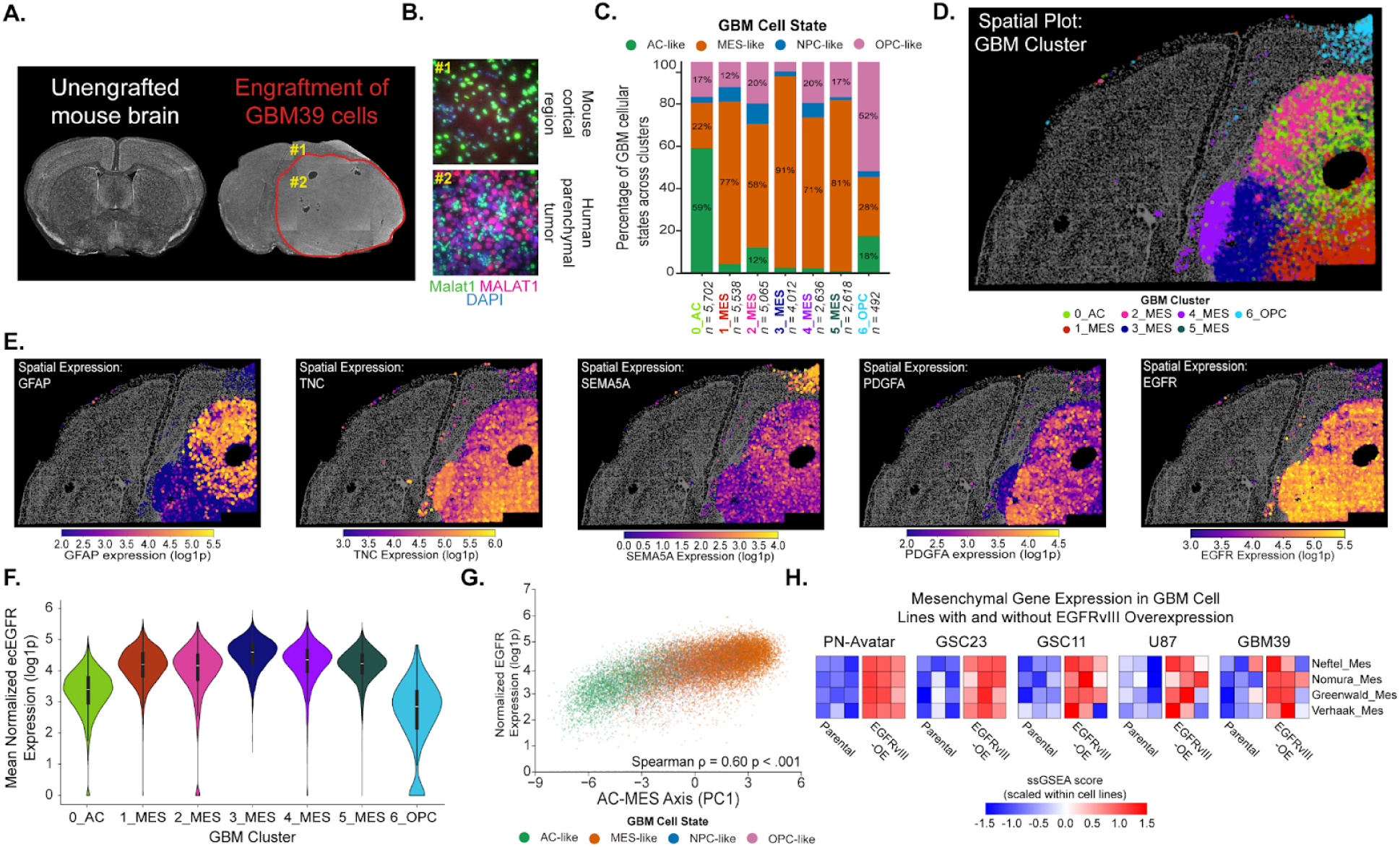
GBM transcriptional states are spatially organized and influenced by EGFR expression. **(A.)** Dapi-stained coronal sections of an unengrafted mouse (left) and a GBM39 engrafted mouse after 3.5 months. The red outline marks the GBM39 parenchymal tumor. **(B.)** Representative smFISH images of *MALAT1* (human) and *Malat1* (mouse) in the cortical region of the brain (#1) and intra-tumoral region (#2) of the GBM39 orthotopic xenograft. Insets are highlighted on the right section from Fig.3A. **(C.)** Stacked bar plot depicting the proportions of different GBM cellular states across the Leiden clusters defined based on RNA-MERFISH data for the human gene library (343 genes). The dominant state was used to annotate the GBM clusters. **(D.)** Image with the spatial location of the GBM clusters identified. Mouse cells are annotated in grey. **(E.)** Image of the single-cell RNA expression measured by RNA-MERFISH across all human cells for different marker genes of the GBM clusters: *GFAP* (0_AC), *TNC* (MES clusters), *SEMA5A* (6_OPC), *PDGFA* (3_MES), and *EGFR*. Mouse cells are annotated in grey. Expression range is indicated for each gene in the corresponding colorbars. **(F.)** Violin plot of *EGFR* expression across GBM clusters. **(G.)** Correlation of Principal Component 1 (PC1 - from single-cell expression data) and *EGFR* expression across single-cells. Cells were colored by GBM cellular state. *N* = 26,063 cells. **(H.)** Heatmap of the single-sample Gene Set Enrichment Analysis (ssGSEA) scores for four^33,37,60,67^ different GBM mesenchymal gene sets across five GBM cell lines with EGFRvIII overexpression (EGFRvIII-OE) and without EGFRvIII overexpression (parental). ssGSEA scores were scaled within cell lines indicating relative MES-like gene expression following EGFRvIII overexpression.

Together, these results demonstrate that ecDNA adopts a globally expanded chromatin configuration with higher transcriptional activity relative to endogenous chromosomes, revealing ecDNA as a physically and functionally distinct chromatin entity rather than a simple amplification of genomic sequence.

### Physical expansion of ecDNA correlates with transcriptional activity

We asked whether physical expansion varies with transcriptional output on a per-molecule basis. Stratifying traces by the number of genes transcribed, we found that more expanded chromatin structures (i.e., larger Rg) transcribed more genes, indicating a strong correlation between Rg and transcriptional activity for both ecDNA and chromosomal loci (**Fig.2A, Extended Data Fig.3C**). These data reveal functional heterogeneity among genomically identical ecDNA molecules and link global chromatin volume to transcriptional activity, which may reflect local openings on chromatin for transcription factors and regulatory proteins to bind.

We next examined whether chromatin expansion and transcription correlate with the local enrichment of regulatory proteins using the immunofluorescence modality of mm-MERFISH. We quantified transcription-associated proteins,^22,43^ such as phosphorylated RNA polymerase II (Pol2S2, associated with transcriptional elongation), active histone mark H3K27ac, spliceosome component SC35, and nuclear structures associated with transcriptional repression, such as nuclear lamina A/C (**Fig.2B**). Integrating these measurements (**Fig.2C**), we observed that chromatin expansion (Rg) and the number of genes transcribed were positively correlated with the deposition of active transcription-associated marks, such as Pol2S2 and H3K27ac, and negatively correlated with the nuclear lamina (**Fig. 2D, Extended Data Fig. 3D**).

Together, these results indicate that coordinated variation in chromatin expansion, transcription, and active regulatory machinery underlies the pronounced heterogeneity among ecDNA molecules.

### The interplay between single-molecule regulation and copy number of ecDNA

We next explored whether ecDNA states are coordinated within individual cells or vary independently across molecules, reflecting stochastic versus cell-intrinsic regulation of ecDNA. We compared z-scored differences in ecDNA features—including transcriptional activity, chromatin expansion (Rg), and enrichment of regulatory protein marks—across all ecEGFR molecules and stratified these comparisons within and between cells. ecEGFR molecules within the same cell exhibited significantly smaller differences across these metrics than molecules from different cells (**Fig.2E, Extended Data Fig.3E**), indicating that cellular context contributes to ecEGFR regulation.

We found that single-molecule associations between chromatin expansion, transcription, and active mark enrichment (**Fig. 2D**) reflected a more transcriptionally active cellular state, defined by mean global brightness of active protein marks. Higher mean ecDNA transcriptional output and Rg were positively correlated with increased nucleus-wide deposition of Pol2S2 and H3K27ac (**Fig. 2F, Extended Data Fig 3F**), indicating that intercellular variation in global transcriptional activity is associated with coordinated shifts in ecEGFR expansion and activity.

Additionally, we examined the relationship between ecDNA copy number and the magnitude of this high Pol2S2 state across cells. First, consistent with prior findings,^11,19,44^ ecEGFR copy number varied substantially across cells (**Fig. 2G**) and mean ecDNA counts per cell correlated with exon counts for all ecDNA-encoded genes (**Extended Data Fig 3G-I**), underscoring the important role of copy number in regulating ecDNA expression. Stratification into low- and high-copy number groups revealed that ecDNA molecules from high-copy number cells exhibited increased Rg, higher bursting rates, and elevated active protein marks (**Fig. 2H-I, Extended Data Fig.3J**). In contrast, there was no difference in lamin deposition across these two groups (**Extended Data Fig.3K**).

Collectively, these findings show that ecDNA structure and transcriptional activity are coordinated within individual cells, revealing a cell-intrinsic layer of ecDNA regulation that operates alongside copy number variation. Specifically, our analysis revealed that one state determinant of GBM––global Pol2S2 within the cell––is associated with physical expansion and transcriptional activity of ecDNA. Notably, these features of ecDNA were also higher in cells with more ecDNA molecules, suggesting an interplay between copy number and single-molecule regulation of ecDNA.

### Subnuclear localization and micronuclear residence modestly influence ecDNA

Beyond measuring DNA, RNA, and protein, mm-MERFISH retains sub-micron spatial information. The genome is organized at several scales within nuclei: chromosomes occupy a defined volume within the nucleus, called chromosomal territories; at the subchromosome scale, Mb-sized DNA stretches are arranged in A (active, or euchromatin) or B (repressed, or heterochromatin) compartments; and at a finer scale (100’s of Kb - Mb), DNA is organized into topologically associating domains (TADs).^45^ While our single-molecule chromatin tracing revealed chromatin domain structures, we wanted to explore whether other layers of genome organization apply to ecDNA. We observed that active and repressive mark deposition displayed a reciprocal pattern along the 5-Kb segments of ecEGFR (**Extended Data Fig. 4A**), recapitulating the previously shown spatial segregation of active and repressive chromatin states.^22,45^

Additionally, lamina associated domains (LADs), chromosomal domains that are in proximity with the nuclear lamina, tend to be more transcriptionally repressed and associated with heterochromatin compared to those that are not.^46^ Investigating this regulatory logic for ecDNA, we classified each trace as either ‘lamina associated’ (within 750 nm, or the average Rg for each trace) or not (>750 nm), based on the distance from the centroid of each ecDNA to nuclear lamina (**Extended Data Fig.4B-C**). While only a small fraction of the total ecEGFR molecules are lamin-associated (7.5%), they tended to be less transcriptionally active compared to those that are not (**Extended Data Fig.4D**). On the contrary, speckle associated ecDNA molecules showed the opposite trend (**Extended Data Fig.4E-G**), consistent with prior work that showed loci near the speckle reflect more active transcription.^46^

**Figure 4.**
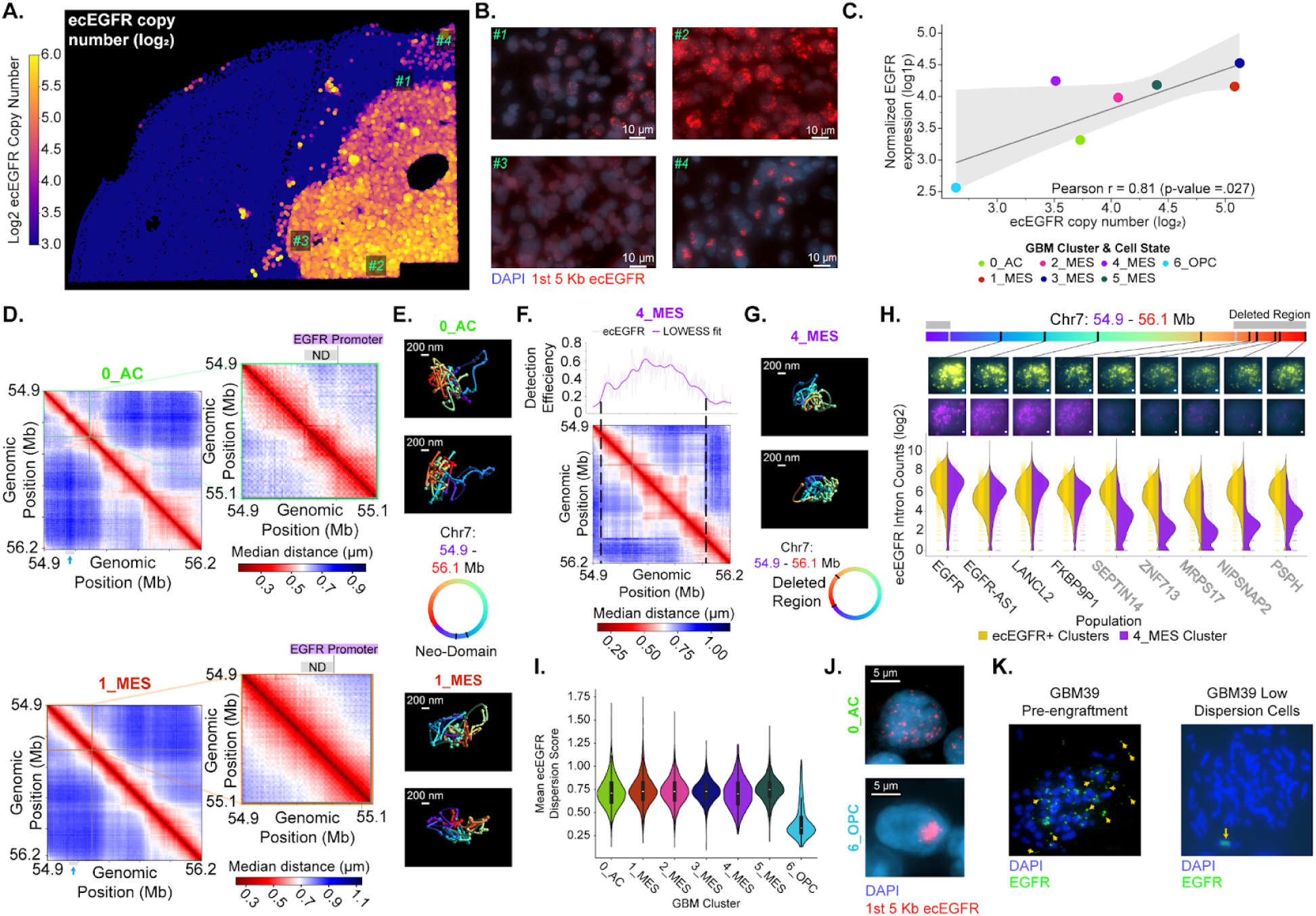
ecDNA exhibits diverse structural and regulatory phenotypes across spatially distinct GBM clusters. **(A.)** Image quantifying number of ecEGFR molecules per cell using DNA-MERFISH. **(B.)** Images of DNA-MERFISH of the first 5 Kb segment (red) of ecEGFR across phenotypically diverse regions: #1 - low copy number, #2 - high copy number, #3 - distinct ecDNA structural variant, and #4 - chromosomal reintegration. DAPI - blue. **(C.)** Correlation between mean ecEGFR copy number and mean *EGFR* expression across each GBM cluster. Points are colored by their GBM cluster. Pearson linear regression with confidence intervals (light grey bands), correlation coefficient, and p-value are indicated within the panel. **(D.)** Median distance matrices of all ecEGFR traces within 0_AC (top; green; *N* = 18,161 traces) and 1_MES (bottom; red; *N* = 55,900 traces) clusters. The neodomain (grey box with blue arrow) is annotated at the bottom of the matrices. A grey line marks EGFRvIII deletion. Insets magnify the *EGFR* locus from the start of the ecDNA to the EGFRvIII deletion with the neo-domain (grey) and *EGFR* promoter (purple) annotated on top. **(E.)** Representative ecEGFR traces from 0_AC (top) and 1_MES (bottom). The rainbow color key annotates each ecEGFR trace’s genomic locus from the beginning (purple) to end (red) and the location of the neo-domain (blue-cyan) identified in Fig.4D. **(F.)** The median distance matrix for all ecEGFR traces from 4_MES cells. The detection efficiency (lavender) and Locally Weighted Scatterplot Smoothing (LOWESS) line (purple) were computed and plotted for each bin above the distance matrix. Dotted black lines indicate where detection efficiency is low, representing a potential intra-ecEGFR deletion. *N* = 27, 253 traces. **(G.)** Representative ecEGFR traces from 4_MES. The rainbow color key annotates each ecEGFR trace’s genomic locus from the beginning (purple) to end (red), and the location of the deleted region identified in Fig.4F. **(H.)** Split violin plots of intronic counts for all ecEGFR genes across GBM clusters, excluding 4_MES and 6_OPC, (left; gold; 22,935 cells) and the 4_MES cluster (right; purple; 2,636 cells). Images of nascent transcription of ecEGFR genes from a representative cell containing full length ecEGFR (gold) and a representative cell from 4_MES cluster (purple). The scale bars represent 1 µm. The genomic locations of genes are annotated (black bar) on a rainbow colored ecDNA schematic. The structural variant deletion is marked by a grey bar. Black lines connect each gene to its nascent transcription image. **(I.)** The distribution of ecEGFR dispersion scores across clusters. Differences between clusters were assessed using a Kruskal-Wallis test (H_6_ = 546, *p* = 1.31 × 10⁻¹¹⁴). **(J.)** A representative cell from 0_AC and 6_OPC showing high (top) and low (bottom) ecDNA dispersion scores, respectively. DAPI - blue and 1st 5 Kb of ecEGFR - red. **(K.)** Metaphase spreads of pre-engraftment GBM39 *in vitro* (left) and isolated extraparenchymal tumor-derived GBM39 cells *in vitro* (right). EGFR FISH - green and DAPI - blue. Yellow arrows indicate *EGFR* signal on ecDNA molecules from a GBM39 cell (left) and chromosomal reintegration of ecDNA from an extraparenchymally-derived GBM39 cell (right).

Beyond the nucleus, ecDNA molecules have been reported to reside within micronuclei, small aberrant nuclear structures that arise from and contribute to genome instability.^47^ We observed micronuclei in roughly 20% of all imaged cells (**Extended Data Fig.4H**), aligning with a recent study of GBM39.^48^ The micronuclei distribution was heterogeneous across cells and was logarithmically distributed with a small population of cells having up to 4 micronuclei (**Extended Data Fig.4H**). 30% of micronuclei contained ecEGFR, the majority of which were actively transcribing *EGFR* (**Extended Data Fig.4I-J**). Bursting rates of ecEGFR molecules within micronuclei significantly correlated to those within the cell’s main nucleus (Pearson r = .986, **Extended Data Fig.4K**). Interestingly, the presence of micronuclei significantly correlated with ecEGFR copy number and transcription (**Extended Data Fig.4L**), raising the possibility that increased copy number and/or expression may drive micronuclei formation.

Together, these findings indicate that while subcellular contexts such as lamina-, speckle-, or micronuclear-association can modulate ecDNA activity, they do not account for the dominant sources of ecDNA heterogeneity. Instead, ecDNA regulation primarily reflects coordinated cell-level programs, providing a framework for understanding how ecDNA structure and activity may be shaped by broader biological contexts, including the tumor microenvironment.

### Spatially segregated GBM subpopulations with distinct transcriptional states emerge in the murine brain

After characterizing ecEGFR *in vitro*, we investigated how the brain environment shapes transcriptionally-defined states of GBM cells in an orthotopic xenograft model. GBM39 cells were engrafted into the striatum of an athymic nude mouse (**Fig.3A**), and once the mouse was moribound (∼3.5 months), the brain was extracted, sectioned, and hybridized to multiple mm-MERFISH libraries: ecEGFR RNA/DNA libraries (as in Fig.1-2), a mouse CNS cell type library (275 genes), and human CNS cell type library (343 genes) (Supplementary Table 1). In this chimeric system, probes targeting mouse and human transcripts, including *MALAT1*/*Malat1*, were used to distinguish non-malignant (mouse) and malignant (human) cells (**Fig.3B, Extended Data Fig.5A-C**). While the ipsilateral cortical region of the brain consisted mainly of mouse cells (**Fig.3B, top, green**), the tumor mass was a complex environment composed of both non-malignant and malignant cells (**Fig.3B, bottom, green and red**). The malignant cells were clustered based on their single-cell RNA expression into seven Leiden^49^ clusters, which possessed different proportions of four GBM cellular states^33^: MES-like, AC-like, NPC-like, and OPC-like (**Fig.3C, Extended Data Fig.5D-E)**. Overall, most clusters were predominantly MES-like except for the largest (0_AC) and smallest (6_OPC) clusters, which were predominantly AC-like and OPC-like, respectively (**Fig.3C)**.

**Figure 5.**
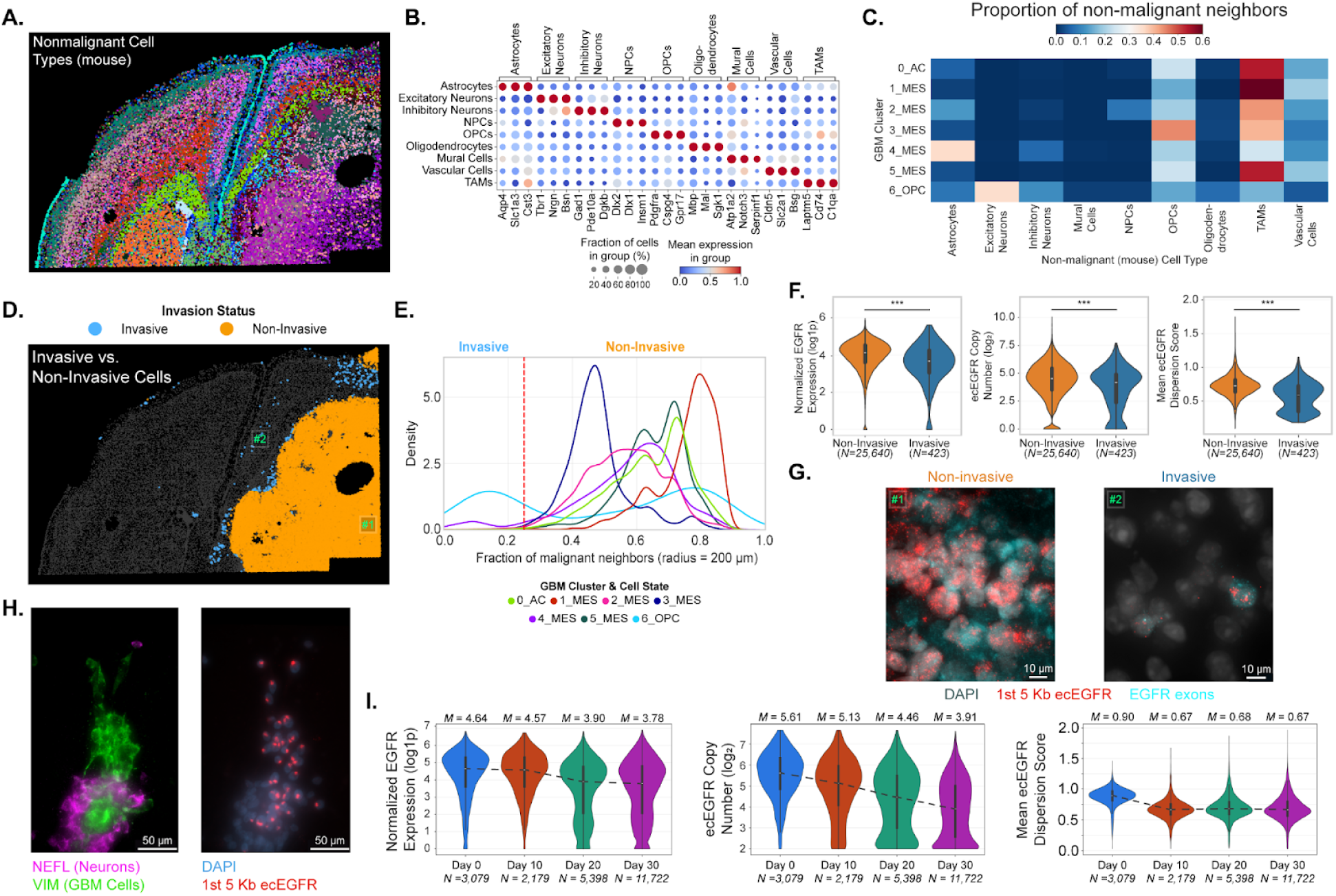
Microenvironmental selection of ecEGFR phenotypes across GBM transcriptional and functional states. **(A.)** Spatial distribution of leiden clusters across non-malignant (murine) cells. **(B.)** Dot plot marking differential expression across major cell types. **(C.)** Heatmap quantifying the cellular composition of non-malignant cells neighboring GBM clusters. For each GBM cell, the 50 nearest non-malignant neighbors were pooled across GBM clusters. The relative proportion of each non-malignant cell type is shown. **(D.)** Spatial distribution of invasive (blue) and non-invasive (orange) malignant cells. The fraction of malignant cells within a 200 µm radius of each GBM cell was used to define invasive (< 0.25 malignant fraction) and non-invasive (>0.25 malignant fraction) cells. **(E.)** Distribution of fraction of malignant neighbors for each GBM cluster. The dashed red line marks the 0.25 threshold used to define invasive status. **(F.)** Violin plots showing the normalized EGFR exon counts (left), ecEGFR copy number (middle), and ecEGFR Dispersion Score (right) across non-invasive (orange) and invasive (blue) cells. Groups were compared using two-sided Welch’s t-tests with Benjamini–Hochberg FDR correction. *** FDR < 0.0001. **(G.)** Representative images of invasive and non-invasive regions annotated in Fig.5D. DAPI - grey, the 1st 5 Kb region of ecEGFR - red, and EGFR exons - cyan. **(H.)** Representative images of GBM39 cell and neuron co-cultures after 10 days. The left image shows smFISH signals for *NEFL* (magenta) and *VIM* (green), which mark neurons and GBM cells, respectively. The right image shows DAPI (blue) and the first 5-Kb of ecEGFR (red). **(I.)** Violin plots showing the EGFR exon counts (left), ecEGFR copy number (middle) and ecEGFR dispersion score (right) across four timepoints of coculture: day 0, 10, 20, and 30. Median values lie above each violin plot, and a dotted black line connects each median value for each timepoint. Differences between time points were assessed using a Kruskal-Wallis test for each variable. *EGFR* expression: H = 3284.83, p < 1e-6. ecEGFR copy number: H = 1133.3, p = 2.163e-245. Dispersion score: H = 3190.63, p < 1e-6. Additional two-sided Mann-Whitney U tests with Benjamini–Hochberg multiple test corrections were used between the Control and the Day 30 group. *EGFR* expression: MWU = 29249361.0, p < 1e-6. ecEGFR copy number: MWU = 23466895.5, p = 7.918 e-245. Dispersion score: MWU = 5800160.0, p < 1e-6.

GBM transcriptional clusters had distinct spatial organization within the tumor and largely neighbored each other in a homotypic fashion (**Fig.3D**). For instance, 0_AC cells were located in the mid-superior portion of the tumor (**Fig.3D**, green), 6_OPC cells were located in a cortically infiltrative region away from the tumor (**Fig.3D**, cyan), and MES-like clusters were located in different regions throughout the tumor (**Fig.3D**). While some genes, such as *OLIG1*, were ubiquitously expressed by all cancer cells throughout the tumor (**Extended Data Fig.5F**), the classification of GBM clusters and their spatial positions were reaffirmed by the expression of several marker genes, such as *GFAP* for AC-like cells and *TNC* for MES-like cells (**Fig.3E, Extended Data Fig.5F-I**). 6_OPC cells were marked by high expression of *SEMA5A* (**Fig.3E**), a gene recently associated with OPC-like cells.^37^ Across published sequencing data of GBM tumors, 6_OPC marker genes like *SEMA5A*, *SOX8*, and *LINGO1* were co-expressed, enriched in Proneural (i.e., OPC- and NPC-like tumors) tumors, and significantly correlated with genes associated with nervous system development, synapse organization, and OPCs (**Extended Data Fig.6**).^33,50,51^ Lastly, while there were several clusters dominated by MES-like cells, many possessed distinct gene expression patterns such as *PDGFA* in 3_MES cells, *JUN* in 4_MES cells, and *CXCL14* in 2_MES cells (**Fig.3E, Extended Data Fig.5J**), motivating our approach to define GBM subpopulations based on their expression of all 343 genes rather than GBM cellular state marker genes alone.

**Figure 6.**
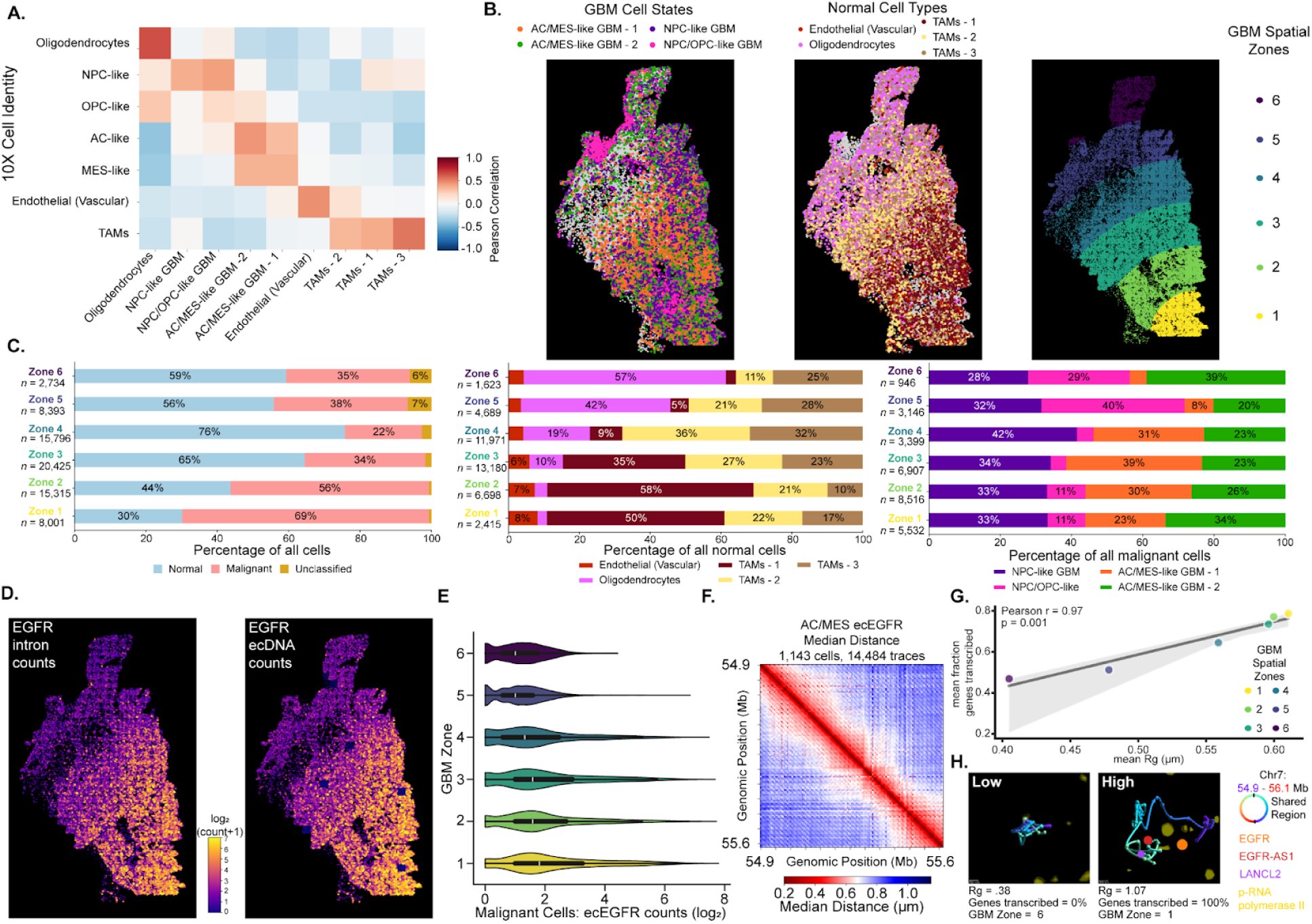
ecEGFR structure, copy number, and transcriptional activity are organized at multiple scales in patient-derived GBM tissue. **(A.)** Heatmap illustrating the correlation of gene expression betweenLeiden clusters from mm-MERFISH+ and cell types from 10X-Multiome. **(B.)** Spatial distribution of malignant (left) cell states and non-malignant (middle) cell types for a primary GBM sample. The right panel marks the definition of GBM spatial zones by dividing the sample into equally sized bins along the vertical axis. **(C.)** Stacked bar plots highlighting different proportions of cell identity across GBM zones: non-malignant vs. malignant (left), normal cell types (middle), and GBM cellular states (right). **(D.)** Spatial distribution of ecEGFR counts and introns per cell across a primary GBM tumor. **(E.)** Violin plot showing the distribution of ecEGFR counts across malignant cells across GBM zones. **(F.)** Median distance matrix for *EGFR* locus traces across ecDNA from AC/MES cells in this sample. Each bin represents the pair-wise distance between two loci where red and blue represent closer and further distances, respectively. **(G.)** Correlation of the mean Rg and mean fraction of transcribing ecDNA genes (*EGFR*, *EGFR-AS1*, and *LANCL2* were measured) from AC-/MES-like cells across each GBM zone. Points are colored based on their GBM zone. From Zone 1 to 6, N = [3882, 4302, 4015, 1801, 378, 106] traces. **(H.)** Representative EGFR locus traces from GBM Zone 6 (left) and 1 (right) with phosphorylated-RNA polymerase II antibody staining (yellow) and introns for *EGFR* (orange), *EGFR-AS1* (red), and *LANCL2* (purple). Intron sizes were scaled by brightness. The rainbow annotation key from the ecEGFR species in GBM39 was kept with the beginning (purple) and end (red) annotated accordingly. Black bins note the shared region that was traced in the patient-derived GBM sample. Rg, genes transcribed, and GBM Zone are annotated below each trace.

We next asked whether the transcriptional diversity we observed *in vivo* preexisted *in vitro* prior to implantation. GBM39 expression of 343 genes significantly correlated across *in vivo* and *in vitro* contexts (**Extended Data Fig.7A**, Pearson r = 0.63, p-value = 1.0e-39), including, for instance, classical GBM marker genes such as *EGFR* and *SOX9* (**Extended Data Fig.7B**). We next compared their transcriptional programs with the GBM39 cells *in vivo* using Ingest^52^, a software that maps single-cell RNA data across datasets (**Extended Data Fig.7C**). Despite their overall correlation, we observed that the *in vitro* GBM39 cells occupied a smaller, more compressed region in transcriptional space with some GBM39 *in vivo* clusters fully absent (i.e., 6_OPC) or greatly underrepresented (i.e., 3_MES) *in vitro* (**Extended Data Fig.7C-D**). Notably, the most differentially expressed genes across GBM clusters *in vivo* had markedly higher expression compared to *in vitro* cells (**Extended Data Fig.7A,E**). Altogether, these analyses suggest that, upon engraftment, GBM39 cells tend to become more transcriptionally heterogeneous consistent with prior studies.^33,53,54^

Next, we considered whether GBM transcriptional and spatial heterogeneity was reproducible across independent mouse replicates. The average gene expression across mouse replicates strongly correlated (Pearson r = 0.73, p-value < 0.0001, **Extended Data Fig.8A**). Upon integrating (Ingest) the GBM cells from one mouse replicate sample into the transcriptional space of the other we observed a more complete coverage of the transcriptional clusters compared to GBM39 *in vitro* cells (**Extended Data Fig.8B-C**). The spatial distribution of the GBM clusters was more variable. For instance, the second mouse replicate exhibited a smaller tumor near the meninges (“extraparenchymal tumor”) in addition to its main parenchymal tumor (“parenchymal tumor”) (**Extended Data Fig.8D**). Nevertheless, the two replicates included many conserved spatial features: MES-like clusters (e.g., 1_MES) were abundant in the tumor cores, 2_MES cells were enriched at the tumor edges, and 6_OPC cells were infiltrating into cortical brain regions (**Extended Data Fig.8D**, **Fig.3D**). Spatial gene expression patterns were also largely retained, with some genes ubiquitously expressed like *OLIG1* and others spatially variable, such as *TNC* in the tumor core or *PCLAF* in infiltrative cells (**Fig.3E, Extended Data Fig.8E, 5F-I)**. We re-defined GBM Leiden clusters and cellular states on mouse replicate 2 and recapitulated cellular state proportions (MES >> AC > OPC ≥ NPC) (**Extended Data Fig.8D, F-H, 5D**). While the parenchymal tumor was predominantly MES-like, the extraparenchymal tumor exhibited an AC-like state marked by genes such as *GFAP* (**Extended Data Fig.8H-I**).

Altogether, these results show that the brain microenvironment promotes transcriptionally diverse GBM cell populations that are spatially organized within the tumor, motivating further investigation into the drivers of this intratumoral heterogeneity.

### EGFR expression influences transcriptional heterogeneity in GBM

We next characterized the relationships between *EGFR* expression, cellular state, and spatial organization. *EGFR* expression significantly varied across GBM clusters in a spatially distinct manner (**Fig.3E-F**). Specifically, 0_AC cells had lower *EGFR* expression than all MES-like clusters, and 6_OPC cells had the lowest mean *EGFR* expression in the entire tumor (**Fig.3F**). We performed PCA on the RNA expression of malignant cells and found that Principal Component 1 (PC1) defined a continuous transcriptional axis spanning AC- to MES-like states for both mouse replicates (**Fig.3G, Extended Data Fig.8J**). This AC- to MES-like score correlated strongly with *EGFR* expression *in vivo* in both mouse replicate 1 (Spearman correlation coefficient = 0.60, **Fig. 3G**) and mouse replicate 2 (Spearman correlation coefficient = 0.85, **Extended Data Fig.8J**) but only modestly *in vitro* (Spearman correlation coefficient = .36, **Extended data Fig. 7F**). The genes that correlated most strongly with *EGFR* expression were also the strongest drivers of this AC- to MES-like axis in both mice replicates (**Extended Data Fig.7G, 8K**).

We hypothesized that increasing expression of *EGFR* could shift GBM cells toward a MES-like state. To test this, we overexpressed *EGFRvIII* in GBM39 and four other transcriptionally distinct GBM cell lines and performed bulk RNA sequencing (**Extended Data 9A-D**). For all cell lines, we found that *EGFRvIII* overexpression led to higher expression of MES-like genes (**Fig.3H**) and promoted AC- or MES-like states (**Extended Data Fig.9F**). Interestingly, GBM39—the only cell line with high basal levels of *EGFR* expression—reduced AC-like gene expression following *EGFRvIII* overexpression, consistent with our *in vivo* data (**Extended Data Fig.9A, E-F**). Conversely, treatment of GBM39 with the *EGFR* inhibitor erlotinib (GBM39-ERL) reduced ecEGFR copy number,^12,19^ *EGFR* expression, and the MES-like score in a correlated manner *in vitro* (**Extended Data Fig.9G**).

Collectively, our analysis linked spatially and transcriptionally distinct GBM subpopulations with particular levels of *EGFR* expression. Genetic or pharmacologic modulation of *EGFR* dosage was sufficient to shift global gene expression programs, suggesting that *EGFR* levels influence GBM states and their spatial organization *in vivo*. These results prompted us to investigate how different levels of EGFR expression is regulated on the ecDNA across different GBM clusters.

### ecDNA exhibits diverse structural and regulatory configurations across spatially distinct GBM subpopulations *in vivo*

We observed multiple mechanisms acting on ecDNA, including variation in copy number, 3D structure, genetic composition, and chromosomal reintegration across tumor compartments (**Fig.4A-B**). ecEGFR copy number and EGFR expression varied in a correlated manner across spatially distinct GBM clusters: highest in the tumor core (MES-like clusters), lower in superior tumor (0_AC), and lowest in cortically infiltrating cells (6_OPC), suggesting that copy number variation is a key modulator of ecDNA expression (**Fig.4A-C**). Mean Rg and transcription of all ecEGFR-encoded genes were also correlated and varied across GBM subpopulations: lowest in 0_AC and highest in MES-like clusters, such as 1_MES (**Extended Data Fig.10A**). These results suggest copy-number independent mechanisms of ecEGFR regulation *in vivo* and align with our *in vitro* analysis in which higher transcriptional activity of ecEGFR correlated with a globally expanded structure (**Fig.2A**).

We compared the 3D structure of ecEGFR across the two largest GBM clusters: 0_AC and 1_MES (**Fig.4D**). While the overall chromatin domain structures were similar across the two subpopulations, a ∼750 Kb region neighboring the *EGFR* promoter possessed unique structural differences (**Fig.4D**). Notably, a new smaller chromatin domain formed upstream of the EGFR promoter (chr7:55.100-55.175-Mb) in 0_AC cells (**Fig.4D**). This 0_AC-specific neo-domain was associated with higher intra-region contact (**Extended Data Fig.10B**). Visually appreciable at the single-molecule level, the chromatin neo-domain (colored blue) is physically separated from the ecDNA molecules in the characteristic traces of 0_AC cells but not 1_MES cells, as quantified by its centrality score^55^ (**Fig.4E, Extended Data Fig.10C**). EGFR amplifications have been shown to possess a 5’ skew due to the presence of cis-regulatory elements upstream of its promoter whose activities are crucial for *EGFR* expression and GBM cell viability.^56^ Population average epigenomic data of GBM39 cells *in vitro*,^15^ which did not display the neo-domain (**Fig.1D**), showed significant interactions between the EGFR promoter and putative regulatory regions (H3K4me1) within the neo-domain (**Extended Data Fig.10D**). These results show chromatin reorganization in specific GBM clusters of regions expected to modulate *EGFR* expression.

We observed that 4_MES cells possessed a unique structural variation of ecEGFR (SV-ecEGFR, **Fig.4B, inset #3**). Specifically across traces in these cells, there was low detection of 5-Kb segments at the beginning (chr7: 54.925 - 55.000-Mb) and end (chr7: 55.977 - 56.207-Mb) of ecEGFR, suggesting an incomplete structural variant (**Fig.4B, inset #3, 4F**). This was also manifested in the median distance matrix showing an inner block with smaller interdistances between retained loci. Representative traces lacked DNA segments from this region (chr7:55.000 - 55.977-Mb, **Fig.4F-G**). Consistent with an intra-ecEGFR deletion, nascent transcription of the five genes that fell within this region was nearly undetectable in these cells (**Fig.4H**). In accordance with previous work,^17,57,58^ this analysis underscores sequence evolution as another mechanism by which cancer cells can alter ecDNA expression and highlights the ability of mm-MERFISH to provide direct visualizations of different structural variants of ecDNA, which may be obscured by ensemble-averaged approaches.

Cortically infiltrative 6_OPC cells harbored tightly clustered ecEGFR molecules within their nuclei, with a size of ∼2 µm, the size of chromosomal territories (**Fig.4B, inset #4**). For each cell, we calculated an ecDNA dispersion score: the average distance between the ecEGFR molecule and their center of mass, normalized by what is expected from a uniform sampling distribution (Methods). 6_OPC cells exhibited a significantly lower ecDNA dispersion score (**Fig.4I-J**). This phenotype appeared prominently in the extraparenchymal tumor of mouse replicate 2 (**Extended Data Fig.11**): In addition to having less *EGFR* expression and copy number relative to the parenchymal tumor (**Extended Data Fig.11A-C**), the extraparenchymal tumor also exhibited tightly clustered ecEGFR molecules with a significantly lower ecDNA dispersion score similar to cortically infiltrating 6_OPC cells of mouse replicate 1 (**Extended Data Fig.11D, E**). Extraparenchymal tumors were a recurrent feature of this xenograft model and consistently exhibited clustered ecEGFR, observed across several independent replicates (**Extended Data Fig.11F**). They formed in mice treated with erlotinib as well as in non-treated mice, indicating that their occurrence was independent of drug treatment (**Extended Data Fig.11F**). Even upon matching ecEGFR copy number, *EGFR* expression varied between parenchymal and extraparenchymal tumors, suggesting epigenetic modulation of ecEGFR (**Extended Data Fig.11G**). We dissected and cultured cells from an extraparenchymal tumor and observed lower average ecEGFR copy number, *EGFR* RNA, and EGFR protein levels relative to pre-engrafted GBM39 cells by qPCR and FACS (**Extended Data Fig.11H-K**). EGFR FISH of extraparenchymal cells recapitulated tightly clustered ecEGFR signals in interphase (**Extended Data Fig.11L**) and, notably, revealed ecEGFR chromosomal re-integration in metaphase spreads (**Fig.4K**).

Together, these results demonstrate that ecEGFR is structurally and functionally plastic, and its copy number, chromatin organization, sequence composition, and reintegration configurations vary across GBM subpopulations and tumor regions. This spatial heterogeneity prompted us to investigate how interactions with neighboring non-malignant cells shape the emergence of distinct ecDNA regulatory mechanisms across GBM states.

### The microenvironment shapes the phenotypic diversity of ecDNA and GBM states

GBM subpopulations were spatially associated with distinct microenvironments defined by non-malignant cell type composition and degree of invasion (**Fig.5**). Mouse cells were clustered based on their single-cell expression of 275 marker genes (Supplementary Table 1). We defined the major cell types of murine cells which had characteristic marker gene expression and spatial distributions^59^ (**Fig.5A, Extended Data Fig.12A-B**). Interestingly, some cell types, including tumor-associated macrophages (TAMs), vascular cells, and astrocytes, had distinct transcriptional clusters within and outside the tumor, reflecting tumor-associated expression differences (**Extended Fig.12C-E**). For instance, relative to endothelial cells outside of the tumor, intratumoral endothelial cells had higher expression of *Cldn5*, an important component of blood brain barrier tight junctions (**Extended Data Fig.12F**). Other genes, such as *Junb*, *Cq1a*, and *Mki67*, were broadly upregulated in non-malignant cells within the tumor, indicating cell-type independent gene expression changes in response to GBM cells (**Extended Data Fig.12F-I**). For all GBM clusters, we defined the average composition of neighboring non-malignant cells (Methods), revealing distinct microenvironments for several GBM subpopulations (**Fig.5C**). The predominant non-malignant neighbor of GBM cells was TAMs (**Fig.5C, Extended Data Fig.12E**), consistent with prior reports in human GBM samples.^33,37,60,61^ OPCs were the second most frequent non-malignant neighbor and were also preferentially enriched within the tumor (**Fig.5C, Extended Data Fig.12E**). Notably, 3_MES cells had the highest proportion of OPC neighbors **(Fig.5C, Extended Data Fig.12B)** and expressed the highest levels of *PDGFA* in the tumor (**Fig.3D-E)**––the ligand for the OPC receptor *PDGFRA*––suggesting that the tumor milieu may promote OPC proliferation and/or migration. Infiltrative 6_OPC cells with chromosomally re-integrated ecEGFR had the lowest amount of TAM neighbors and predominantly colocalized with cortical excitatory neurons, consistent with their high expression of genes related to neuronal development and function such as *SEMA5A* and *LINGO1* (**Fig.3D-E, 5C, Extended Data Fig.6**). 4_MES cells, which contained SV-ecEGFR, were uniquely positioned near astrocytes **(Fig.5C)** and disrupted striatal medium spiny (*Pdea10a*+) neurons flanking this GBM subpopulation **(Extended Data Fig.13A-D**). We detected a small number of 1_MES cells with SV-ecEGFR that were spatially and transcriptomically connected to 4_MES cells (**Extended Data Fig.5E, 13E-F**). 1_MES cells had progressively higher copy numbers of SV-ecEGFR as they radiated out from the tumor core and converged toward 4_MES cells, suggesting selection of SV-ecEGFR in a spatially and transcriptionally distinct subpopulation (**Extended Data Fig.13E-F**). Collectively, this analysis highlights the spatial and transcriptional relationships between the non-malignant microenvironment and GBM cellular states.

We next characterized GBM functional states based on their capacity to invade beyond the tumor. GBM cells were classified as invasive based on the fraction of malignant neighbors within a 200 µm radius (**Fig.5D-E, Extended Data Fig.14B**). Invasive GBM cells were enriched for 6_OPC cells and NPC-/OPC-like states relative to non-invasive cells (**Fig.5D-E, Extended Data Fig.14D**). Across all GBM cells, *EGFR* expression was significantly lower in invasive cells relative to non-invasive cells across both mouse replicates (**Fig.5F-G, Extended Data Fig.14A-C**). These changes in *EGFR* expression coincided with decreased ecEGFR copy number and more chromosomal re-integration, highlighting the cooperativity of different regulatory mechanisms. These results link *EGFR* dosage, ecDNA regulation, and transcriptional states to malignant cell behaviors like invasion.

We wondered whether additional properties such as extracellular matrix (ECM) adhesion varied across GBM states with different *EGFR* expression and ecDNA phenotypes. Previous studies have shown that reduction in ECM adhesion can serve as a proxy for metastatic potential and invasion.^62–64^ We performed a spinning disc assay,^63^ which subjects cells to a radially increasing shear stress, on two populations of GBM39 with distinct levels of *EGFR* expression: GBM39 and GBM39-ERL (**Extended Data Fig.14E**). Across multiple ECM conditions, greater force was required to detach GBM39 cells compared to GBM39-ERL cells (**Extended Data Fig.14F**). These results suggest that erlotinib-mediated loss of ecEGFR copy number, reduced *EGFR* expression, and acquisition of NPC-/OPC-like states^19^ decreases cancer cell-ECM adhesion. Altogether, this analysis links GBM cellular states harboring distinct ecEGFR phenotypes with particular bio-physical properties (e.g., adhesion) *in vitro* and cancer cell behaviors (i.e., invasion) *in vivo*.

We tested whether the microenvironment can directly affect *EGFR* expression and associated ecDNA regulatory properties by co-culturing iRFP-labeled GBM39 cells with iPSC-derived excitatory neurons. GBM cells grown within a neuronal environment exhibited distinct morphologies such as enhanced arborization with close spatial association with excitatory neuron (*MAP2*+) dendrites (**Extended Data Fig.15A-B**). After 8 days, co-cultured GBM cells trended toward lower ecEGFR copy number measured by qPCR compared to cells grown in identical conditions without neurons (**Extended Data Fig.15C-E**). To better understand ecEGFR dynamics across longer time intervals, GBM39 cells were sparsely co-cultured with neurons and analyzed by mm-MERFISH at days 10, 20, and 30. Marker genes such as *NEFL* and *VIM* were imaged to distinguish neurons and GBM cells, respectively (**Fig.5H**). Overall, ecEGFR copy number and expression consistently decreased over time (**Fig.5I**). The ecEGFR dispersion score significantly dropped by Day 10 after which it largely stabilized, suggesting an environmental selection for chromosomal re-integration (**Fig.5H-I**). These observations indicate that extrinsic cues from excitatory neurons can shift ecEGFR copy number and re-integration, GBM cell morphology, and *EGFR* expression.

Collectively, these results demonstrate that GBM molecular and functional states are associated with distinct cellular ecosystems within tumors. Invasive GBM cells exhibited lower ecEGFR copy number and *EGFR* expression, and neuron co-culture experiments induced these phenotypes over time. These findings support a model in which GBM states and ecDNA behavior are plastic and shaped by tumor–microenvironmental interactions.

### Spatial gradients of *EGFR* dosage and GBM cellular states in patient-derived glioblastoma

We next characterized the phenotypic diversity of ecEGFR and GBM cellular states in primary GBM patient samples. MRI-guided sampling was performed on the tumor core, rim, and infiltrative margin of four newly diagnosed GBM tumors, and extracted nuclei were profiled by 10X-Multiome (**Extended Data Fig.16A**). One tumor, P3, possessed a disproportionate amount of ATAC fragments that mapped to a 2 MB locus encompassing EGFR suggesting the presence of ecEGFR (**Extended Data Fig.16B**), which was confirmed by Droplet HiC and EGFR DNA FISH.^19^ After characterizing cells as malignant or non-malignant using copy number aberration analysis (**Extended Data Fig.16C**),^32,65^ non-malignant cell types and malignant cell states^33^ were assigned using their single cell gene expression profiles (**Extended Data Fig.16D-F**). The tumor rim and infiltrative margin were more similar than the tumor core: the tumor core possessed more malignant cells, more TAMs, and more AC- and MES-like GBM cells, while the tumor rim and margin possessed more non-malignant cells, oligodendrocytes, and OPC-like cells (**Extended Data Fig.16G**), consistent with recent characterizations of GBM tumor organization.^60,66^ Aligning with our xenograft GBM model, ecEGFR expression and copy number was non-uniform across tumor regions, with higher copy number and expression in the the tumor core (**Extended Data Fig.16H**). Agnostic to region, ecEGFR copy number and expression were enriched in AC- and MES-like states relative to NPC- and OPC-like states (**Extended Data Fig.16I**). Altogether, this analysis highlights the interplay between ecEGFR copy number and transcriptional variation in human GBM.

To assess whether the spatial and state distribution of ecEGFR copy number and expression are general features of tumors, we re-analyzed a cohort of 26 GBM tumors previously profiled by 10X-Visium, a spatial transcriptomics platform that sequences RNA from 55 μm spots.^60^ Although 10X-Visium only captures RNA, and thereby does not directly measure focal DNA amplifications of *EGFR*, we estimated the presence of ecDNA in malignant spots based on a statistical enrichment of transcripts coming from a 2-Mb window centered on the *EGFR* locus (**Extended Data Fig.17A**, Supplementary Table 3, Methods). 17/26 samples contained spots classified as malignant, and of these, six (35%) were classified as ecEGFR+, consistent with the prevalence of ecEGFR across different GBM cohorts (**Extended Data Fig.3B**).^17^ These samples were marked by high abundance of ecEGFR locus transcripts, including *NIPSNAP2* and *MRPS17*, as observed in GBM39 (**Fig.1I**). Counts from all genes within the 2-Mb *EGFR* locus were aggregated, normalized, and scaled within samples, revealing spatial heterogeneity of *EGFR* locus expression (**Extended Data Fig.17B - top**). In Greenwald et al,^60^ 10X-Visium spots were classified into metaprograms based on gene expression, GBM layers spanning the hypoxic core (L1) to non-malignant pranchema (L5), and infiltrative environment based on an enrichment of neuron, oligodendrocyte, and astrocyte transcripts from neighboring spots.^60^ Consistent with our mouse xenografts and 10X-Multiome analysis, we found that the regional variation in ecEGFR counts varied across these different classifications.^60^ Specifically, neurodevelopmental metaprograms (e.g., NPC) had the lowest counts across all samples whereas hypoxic metaprograms (e.g., MES-Hyp) possessed the highest (**Extended Data Fig.17C**). Aligning with these findings, estimated *EGFR* locus counts were spatially distributed across GBM layers: highest in the hypoxic core layers (L1/2) and lowest in the more infiltrative layers (L3/4, **Extended Data Fig.17D**). Consistent with our previous analysis on invasiveness, malignant infiltrative spots had significantly less counts deriving from the ecEGFR locus compared to malignant non-infiltrative spots (**Extended Data Fig.17B, E**), including across two ecEGFR+ samples that were collected from the core and infiltrative margin of a single patient tumor (**Extended Data Fig.17G-H**).

Collectively, these results reinforce that EGFR dosage and the phenotypic diversity of ecEGFR is associated with transcriptomically, functionally (i.e., invasive), and spatially distinct GBM subpopulations in a broad cohort of primary human GBM.

### The multi-scale organization of ecDNA at single-molecule resolution in patient-derived GBM

To investigate the relationship between ecDNA organization and transcription at single-molecule resolution in patient-derived GBM, we employed mm-MERFISH on an ecEGFR+ primary GBM sample. Using 343 human cell type marker genes, we first clustered cells on their single-cell gene expression profiles and obtained a high correlation of expression with matching cell types defined on the 10X-Multiome data from the same sample (**Fig.6A**). Gene expression measurements were also reproducible across technical replicates (**Extended Data Figure.18A**). Malignant and non-malignant cell clusters had distinct spatial distributions and gene expression (**Fig.6B, Extended Data Figure.18B-D**). We partitioned the sample into six equally sized zones from the bottom of the sample to the top (GBM zones 1-6, **Fig.6B-C**). Collectively, zones 1-4 contained more malignant cells, more TAMs, and more AC/MES-like GBM cells relative to zones 5-6, which consisted of more non-malignant cells, oligodendrocytes, and NPC/OPC-like cells (**Fig.6C**). Consistent with our previous results, the absolute counts of ecEGFR and EGFR expression progressively decreased from the tumor core towards the infiltrative areas (**Fig.6D-E**) and varied with GBM transcriptional states (**Extended Data Fig.18E-H**). PCA across malignant cells defined the main transcriptional axis as a NPC-/OPC-to-AC-/MES-like spectrum (PC1). Along this axis, AC/MES-like cells possessed ecEGFR at higher copy numbers relative to NPC-/OPC-like cells (**Extended Data Fig.18E-H**), underscoring the relationship between intratumoral EGFR dosage and transcriptional heterogeneity in GBM.

We traced the chromatin structure of a 640-Kb region (chr7: 54.945 - 55.582-Mb) of ecEGFR in this primary GBM sample (**Extended Data Fig.19A-B**) and characterized the DNA structure in AC/MES-like cells, which harbored the highest numbers of ecEGFR (**Extended Data Fig.19E-F**). The population average median distance across ecDNA recapitulated the domain-level organization observed in Hi-C (Pearson r = 0.60; **Fig.6F, Extended Data Fig. 19B–D**). Upon analyzing the single-molecule data, we reaffirmed results observed in our analysis of GBM39: (1) ecDNA and chromosomes exhibited nearly identical chromatin domains, but ecDNA was globally expanded relative to its chromosomal counterpart (**Extended Data Fig.19E-G**), and (2) transcriptional activity of ecEGFR significantly correlated with global chromatin expansion (Rg) (**Extended Data Fig.20A**) and enrichment of active transcription markers (**Extended Data Fig.20B**). At the larger scale, within the AC/MES-like state, we observed that ecEGFR molecules in core regions (GBM Zones 1-3) were more globally expanded and transcriptionally active relative to those in infiltrative areas (GBM Zones 4-6, **Fig.6G, Extended Data Fig.20C**), which was visually apparent in representative chromatin traces from Zones 1 and 6 (**Fig.6G-H, Extended Data Fig.20D**). These results support both copy number–dependent and copy number–independent regulatory principles of ecEGFR transcription, even within transcriptionally matched AC-MES-like cells.

Collectively, these results establish that *EGFR* dosage varies across GBM states and is non-uniform across tumor regions in patient-derived GBM tissue. Our single-molecule analysis of ecEGFR revealed that these shifts in *EGFR* expression coincided with changes in copy number and chromatin structure (i.e., physical expansion), which were spatially organized across the broader tumor architecture.

## Discussion

### Defining the organization, regulation, and heterogeneity of ecDNA at single-molecule resolution

We leveraged mm-MERFISH to resolve the 3D structure and transcriptional activity of individual ecDNA molecules within their native multi-scale spatial context. Importantly, this approach integrates microenvironmental context, GBM transcriptional states, and the single-molecule organization of ecDNA. Chromatin tracing and related technologies have primarily been applied to systems with diploid genomes to study genome organization and regulation.^22–26^ A recent pioneering study used chromatin tracing to characterize the evolution of chromatin structure in genetically engineered mouse models, revealing the reorganization of the genome at different stages of tumor progression.^26^ How copy number aberrations, structural variants, and other features of complex karyotypes reshape chromatin organization and transcriptional activity at single-molecule resolution has, until now, remained understudied. Given the prevalence of ecDNA in GBM,^9,17^ we traced ecDNA harboring *EGFR* at 5-Kb genomic resolution in GBM cells *in vitro*, in intracranial xenograft models, and in patient-derived GBM tissue.

### Copy number and chromatin structure cooperate to regulate ecDNA expression

At single-molecule resolution, we found that ecDNA and its matched chromosomal locus exhibited nearly indistinguishable domain structures. Despite these similarities, ecDNA was more physically expanded, defying results from polymer simulations. By simultaneously imaging nascent RNA alongside active transcription markers, we uncovered a strong relationship between ecDNA expansion (Rg), transcriptional activity, and recruitment of regulatory proteins such as phosphorylated RNA polymerase II. This epigenetic regulation aligns with prior work showing enhanced chromatin accessibility and transcriptional activity of ecDNA by comparing tumors or isogenic cell lines with and without ecDNA.^15,20^ Long-read sequencing of ecDNA isolated with CRISPR-CATCH also revealed hypomethylation of ecDNA relative to its matched chromosomal counterpart.^38^ While consistent with our results, these studies largely relied on sequencing-based assays that obscure native spatial context and measure the average ensemble of ecDNA molecules and/or matched chromosomal sequences within a sample (bulk) or, at best, within single nuclei/cells.^15,19,20,38^ By contrast, our single-molecule imaging approach enabled direct measurement of these regulatory principles on individual ecDNA molecules *in situ*.

Despite multiple features of ecDNA transcriptional regulation converging at the single-molecule level, we found that transcript abundance of all ecDNA-encoded genes largely depended on copy number, consistent with prior studies.^13,44^ Interestingly, we observed an interplay between copy number and single-molecule ecDNA regulation: cells in a state with globally higher Pol2S2 levels had increased ecDNA copy number and contained more expanded, transcriptionally active ecDNA. These findings raise the possibility that increasing ecDNA copy number and oncogene dosage may progressively reshape the chromatin of ecDNA itself. These results highlight the cell-intrinsic regulation of ecDNA and foreshadow the coordination of ecDNA within GBM states *in vivo*.

### *EGFR* dosage influences GBM cellular states

Across xenografts and patient-derived GBM tissue, *EGFR* expression markedly varied across distinct GBM subpopulations and strongly correlated with genes that confer GBM cell state identity. In a simplified model, we found that EGFR expression had the following relationships with GBM states: MES-like > AC-like > OPC-like ≥ NPC-like. Genetic and pharmacologic perturbation of EGFR expression shifted GBM states, supporting a model in which *EGFR* can serve as a cellular state “rheostat.” Specifically, *EGFR* overexpression shifted gene expression programs, predominantly enhancing MES-like states, whereas erlotinib treatment reduced *EGFR* expression and suppressed MES-like gene expression. Contextualizing malignant cell behavior with *EGFR* dosage, we show that erlotinib-treated GBM39 cells have decreased ECM adhesion relative to untreated GBM39 cells, suggesting that NPC- and OPC-like states^19^ with lower *EGFR* expression possess bio-physical properties that serve as a proxy for malignant cell behavior like invasion.^62,63^ While *EGFR* amplifications have previously been associated with GBM transcriptional subtypes^67^ and states,^33,37^ our work establishes that the relative levels of *EGFR* expression among malignant cells actively influences several GBM states. Notably, *EGFR* amplifications were initially associated with an AC-like state, and its overexpression in non-malignant neural progenitor cells directly drove a higher AC-like gene expression program.^33^ However, there was no statistical association found between *EGFR* amplifications and AC-like state abundance in an expanded follow-up study.^37^ Given that *EGFR* expression varies substantially within tumors, our “rheostat” framework may help reconcile these previous studies, as *EGFR* levels may vary in a graded manner across different states: from NPC-/OPC-like to AC-like to MES-like. Interestingly, *EGFR* amplifications have recently been associated with a newly defined glial progenitor-like (GPC) state with the potential to differentiate along multiple trajectories into all other GBM states,^37^ raising the possibility that modulating *EGFR* dosage contributes to the plasticity of GPC cells.

Additional cell-intrinsic and microenvironmental mechanisms^33,37,60,64^ also contribute to GBM heterogeneity—e.g., macrophage interactions promoting MES-like states^33,61^—indicating that multiple factors must be considered to account for GBM cellular states and their spatial organization. Whether *EGFR* dosage similarly regulates cellular states in tumors lacking ecDNA remains to be determined. By examining *EGFR*-containing ecDNA, we defined this rheostat model in the context of one of the most prevalent genomic events in newly diagnosed GBM (33-63%), where *EGFR* expression can be dynamically tuned through multiple ecDNA-specific mechanisms.

### Multiple mechanisms enable plastic regulation of ecDNA across scales

ecDNA exhibited pronounced heterogeneity across molecules, cell states, and tumor regions, enabling transcriptional regulation by several mechanisms: evolution of DNA sequence, 3D chromatin reorganization, chromosomal reintegration, and copy number variation. Consistent with the role of ecDNA evolution and heteroplasmy in cellular adaptation,^16,17,19^ we observed the formation and selection of a structural variant ecDNA in a transcriptomically and spatially distinct GBM subpopulation, which led to nearly undetectable levels of expression of the five genes that fell within the deleted region. Beyond sequence variation, the 3D structure of ecDNA also varied across GBM cell populations, reinforcing the relationship between physical expansion and transcriptional activity of ecDNA. In an extreme case, we found chromosomal reintegration of ecDNA in a GBM subpopulation with the lowest levels of *EGFR* expression in the entire tumor. Finally, consistent with *in vitro* experiments, we also observed pronounced cell-to-cell variation in ecDNA copy number across distinct GBM subpopulations *in vivo*, which strongly correlated with *EGFR* expression. Our observations are consistent with prior reports showing non-uniform ecDNA copy number across tumor regions^17^ and variation in ecDNA abundance and chromosomal reintegration across metastatic sites.^18^ While these insightful studies highlight how ecDNA phenotypes can vary in a context-dependent manner, they do not link this phenotypic diversity to malignant cellular states and the broader microenvironment.

### The phenotypic diversity of ecDNA and GBM states depend upon the local microenvironment

Distinct transcriptional and functional GBM states emerge depending on the cellular microenvironment. Mapping GBM states and ecDNA phenotypes in space revealed a striking microenvironmental organization. Specifically, AC- and MES-like GBM cells neighbored TAMs in the tumor core, whereas infiltrative regions were enriched with NPC- and OPC-like GBM cells neighboring neurons and oligodendrocytes. Accordingly, *EGFR* expression was higher in the tumor core relative to infiltrative regions, and multiple, complementary regulatory mechanisms acting on ecDNA modulated *EGFR* expression. Generally, ecDNA was more expanded, transcriptionally active, and present at higher copy numbers in tumor core regions, whereas infiltrative regions were characterized by lower ecDNA copy number, reduced transcriptional activity, and more physically compact chromatin structures with additional regulatory mechanisms like chromosomal reintegration in distinct populations. Consistent with a causal role of heterotypic inputs influencing ecDNA phenotypes, longitudinal co-culturing of GBM39 cells with excitatory neurons led to reduced *EGFR* expression, decreased ecDNA copy number, and increased chromosomal reintegration. Aligning with our findings, single-cell RNA sequencing,^66^ intravital imaging,^66^ and spatial transcriptomic^60,64^ approaches have defined a clustered and layered organization of GBM tumors^60^ and highlighted the role of non-self neighbors in driving GBM plasticity and behaviors like invasion.^33,61,64,66^ However, these studies lack direct genomic context. With multi-modal spatial genomics, we add an ecDNA dimension to these findings, revealing the interplay between genetic and microenvironmental influences in shaping cellular states, malignant cell behaviors, and the broader tumor organization.

### Clinical and technological implications

Intratumoral heterogeneity and cancer cell plasticity drive therapy resistance and remain fundamental obstacles in the treatment of cancer.^5,30^ *EGFR* inhibitor clinical trials have failed in GBM despite their success in other cancer types,^30,68^ raising the possibility that the prevalence of ecDNA in GBM and its capacity to drive cellular adaptation may contribute to poor patient outcomes. We show that ecDNA contributes to intratumoral heterogeneity and plasticity by modulating oncogene expression through multiple, complementary mechanisms, thereby influencing cellular states. Therapeutically targeting *EGFR* signaling may select for particular ecDNA phenotypes (e.g., low copy number or chromosomal reintegration); shift GBM cellular states (e.g., from AC-/MES- to NPC-/OPC-like); and reconfigure the tumor architecture (e.g., promoting diffusely infiltrating cells). As such, beyond directly targeting ecDNA,^20,69^ anticipating how GBM cells change in response to targeted therapy may enable rational combination therapies. For example, targeting NPC-/OPC-like cells in combination with *EGFR* inhibitors may be an effective therapeutic approach, consistent with the “state-selective lethality”^70^ paradigm. In addition to its implications for GBM therapy, this work lays the foundation for future technology development in cancer spatial genomics. Specifically, we envision extending mm-MERFISH to target hundreds of structural variants, enabling comprehensive characterization of cancer genomes and their impact on tumor organization and evolution. Because of the heritability of DNA, we expect improved library complexity would enable clonal and tumor evolution mapping, as foreshadowed by the emergence and selection of a structural variant ecDNA in our xenograft model. In summary, these findings provide an unprecedented window into intratumoral heterogeneity, revealing how genomic variation across scales shapes cell identity and malignant behavior within the broader tumor architecture.

## Methods

### Cell Culture

GBM39 cells were cultured as described previously.^12,19^ Briefly, GBM39 cells were grown in DMEM/F12 media (Gibco) supplemented with B27 without vitamin A (Gibco/Life Technologies), 20 ng/mL human recombinant EGF (Fisher), 20 ng/ml bFGF (Stemcell Tech), and 1% PenStrep (Fisher) in incubators at 37 °C and 5% CO_2_. GBM39 cells grown as neurospheres were dissociated every 7-10 days, and media was replaced every 3-4 days. GBM39 cells grown adherently were cultured on Geltrex (Fisher) according to manufacturer’s guidelines. GBM39-ERL were grown in 5 µM of Eroltinib (Fisher) reconstituted in DMSO for at least 30 days before assaying. Following 30 days, GBM39-ERL was maintained in 5 µM of Eroltinib with drug replacement every 3 days.

For EGFRvIII overexpression studies, four additional glioblastoma cell lines were used: one established cell line (U87), two glioma stem cell lines (GSC11 and GSC23), and one engineered glioma model (Proneural-Avatar). U87 was obtained as described previously.^71^ GSC lines were obtained as described previously.^72^ The Proneural Avatar (PN-Avatar) glioma neurosphere cell line was generated from an iPSC line engineered to harbor glioma driving mutations. Once edited, cells were then differentiated into neural progenitor cells and serially engrafted into mice brains where they formed tumors. This cell line underwent two rounds of engraftment before being cultured indefinitely *in vitro*. The methods to generate this line were outlined previously.^73^ The GSC and avatar lines were grown with the same protocol as GBM39. U87 was grown adherently on tissue culture plates or within tissue culture flasks using DMEM/F12 media (Gibco) supplemented with 10% FBS (Benchmark) and 1% PenStrep (Fisher).

iNPC12 is a iPSC-derived neural progenitor cell line that was used as a diploid control for qPCR experiments. Differentiation and maintenance of this cell line was performed as described previously.^73^

### hIPSC-derived excitatory neurons and co-culture

WTC11 iNGN2 iPSCs (a gift from Yin Shen’s lab, UCSF) were cultured in Essential 8 Flex medium (Thermo Fisher, A2858501) on Matrigel-coated dishes (Corning, 354277) at 37°C and 5% CO_2_ as previously described.^74^ Cells were passaged when they reached ∼70% confluence or every 3 days. Media was supplemented with 1X CultureCEPT (Thermo Fisher, A56799) during passaging.

Neuronal differentiation was done as previously described.^74^ Briefly, one million iPSCs were seeded in 10 cm dishes in KnockOUT DMEM/F12 medium (Thermo Fisher, 12660-012) containing 1X N2 supplement (Thermo Fisher, 17502048), 1X NEAA (Thermo Fisher, 11140-050), 10 ng mL-1 BDNF (STEMCELL, 78005.1), 10 ng mL-1 NT-3 (PeproTech, 450-03), 1 μg mL-1 mouse laminin (Thermo Fisher, 23017015), and 2 μg mL-1 doxycycline (Sigma, D9891). Media was replaced daily for two days, and on day 3 of differentiation cells were dissociated with Accutase (STEMCELL, 07920). Up to two millions cells were then seeded onto 40 mm #1.5 coverslips (Bioptechs, 40-1313-0319) coated in poly-L-ornithine (Sigma, P3655) in a 1:1 mixture of Neurobasal-A Medium (Thermo Fisher, 12349015) and DMEM/F12 (Thermo Fisher, 13330-032) containing 1X B27 supplement (Thermo Fisher, 17504044), 1X NEAA, 0.5X N2 supplement, and 0.5X GlutaMAX (Thermo Fisher, 35050061). Throughout differentiation, 50% of the media was replaced every 3 days.

For co-culture experiments, GBM39 neurospheres were dissociated per methods described above, re-suspended in Neuronal media, and added to hIPSC cultures at low cell numbers. Specifically, for iRFP imaging and FACs experiments, approximately 50,000 cells were added per 10 cm plate containing 1 million Neurons (1:20 ratio) at least 10 days post-differentiation. Additional GBM39 cells were grown separately in 10 cm dishes with the same culture conditions above as a control. iRFP was monitored daily using a BZ-X800 microscope (Keyence) for eight days. At day 8, cells were dissociated, FACS was performed using the methods above. DNA was isolated from GBM39 cells, and qPCR was performed, as described above, to measure relative ecDNA copy numbers.

For MERFISH and IF experiments, approximately 5,000 cells were added per coverslip containing approximately 100,000 Neurons (1:20 ratio). Cells were fixed and assayed by IF (see Methods below) 8-10 days post co-culture. For mm-MERFISH experiments, after the appropriate duration in time, coverslips were aspirated, washed with PBS, and gently fixed with 4% PFA with RNAse inhibitors (New England Biosciences, M0314L). Coverslips were then hybridized to probes immediately, or frozen in PBS with 10% glucose solution.

### Orthotopic Xenografts

As described previously,^12,75^ animal studies were performed in compliance with UC San Diego Animal Care Program under the approved protocol S00192M. For tumor implantation, female BALB/c nude mice aged 4-8 weeks were stereotactically engrafted with 100,000 - 200,000 cells suspended in 3-5 uL 1.0 mm anterior and 2.0 mm lateral to the bregma at a depth of 2 mm from the inner skull surface. Mice were monitored for symptoms (weight loss, motor dysfunction, behavioral changes) and euthanized according to institutional animal welfare guidelines once moribound. For *in vivo* drug studies (Extended Data Fig.11), erlotinib was reconstituted at 100mg/ml in water with 5% DMSO, 0.5% methyl cellulose, and 0.5% Tween-80 and administered at 100mg/kg dose at a 5 days on 2 days off schedule until moribound.

### Flow cytometry

Flow cytometry was performed on a Sony SH800S cell sorter (Sony Biotechnology). For all experiments, cellular debris was excluded based on forward- and side-scatter area (FSC-A and SSC-A), and singlets were identified using FSC-A versus FSC-H gating. Data were analyzed using the Sony SH8000S software and Flowjo v10.10 (BD Biosciences).

For co-culture experiments, GBM39 cells were lentivirally transduced with iRFP720 to distinguish malignant cells from neurons. Cells were washed with PBS, lifted with Accustase (Innovative Cell Technologies), and resuspended in cold PBS supplemented with 1% FBS. Single cell suspensions were passed through 40 μm cell strainers (Fisher, 22363547) into FACS tubes prior to analysis. iRFP720 fluorescence was detected using the 700 nm laser line. Gates were established using non-transduced GBM39 cells as negative controls.

To measure single cell EGFRvIII protein levels, cells were incubated with a vIII-specific EGFR antibody according to the manufacturer’s guidelines. Briefly, cells were incubated with primary antibody (Anti-EGFRvIII, clone DH8.3, Millipore Sigma, MABS1915, 1:100 dilution), washed with PBS + 1% FBS, and incubated with anti-mouse secondary antibody (ThermoFisher, A21202, 1:1000 dilution) at room temperature for one hour on a shaker and protected from light. Gates were established using GBM39 cells incubated with the secondary antibody without primary antibody incubation. Fluorescence was detected using the FITC (488 nm) laser line.

### Immunofluorescence

For IF, media from co-cultures was aspirated, and cells were washed with 1X PBS and subsequently fixed with 4% PFA for 15 minutes. Samples were then washed with 1X PBS three times and incubated in 0.5% Triton X-100 in PBS for 10 minutes at room temperature to permeabilize cells. Following permeabilization, cells were washed three times 1X PBS and incubated in blocking buffer (5% BSA in PBS containing 0.1% Tween-20; PBTA) for 60 minutes at room temperature. After blocking, primary antibodies—MAP2 (Proteintech, 17490-1-AP, 1:100 dilution), VIM (Cell Signaling, 5741S, 1:100 dilution), and NeuN (AbCam, ab104224, 1:100 dilution)—were diluted in PBTA and incubated with cells overnight at 4°C. The following morning, the primary antibody solution was washed three times with 1X PBS. Secondary antibodies—anti-mouse secondary (Thermo, A-21037, A-31571, A-31570, 1:1000 dilution) and anti-rabbit (Invitrogen, A-11034, A-31573, A-21039, 1:1000 dilution)—were diluted in PBTA for one hour at room temperature and were protected from light. Samples were then washed three times 1X PBS, and coverslips were mounted with ProLong Gold Antifade with DAPI (Invitrogen, P36966) and imaged on a BZ-X800 microscope (Keyence).

### Lentiviral transduction

293T cells were transfected with iRFP720^76^, psPAX2, and pMD2.G packaging constructs using TransIT-VirusGen (Mirus). High-titer lentivirus laden supernatant was collected 48 hours after transfection, filtered through a 0.45 µm cellulose acetate filter, and purified with a LentiX concentrator (Takara Bio). GBM cells were infected with virus for 24 hours at 37C, after which the media was aspirated, the cells were washed with 1X PBS, and fresh standard growth media was added. Infected cells were selected in puromycin, and iRFP expression was determined by fluorescence imaging and FACS.

### Quantitative PCR (qPCR)

For reverse transcription (RT-) qPCR, RNA was extracted using the RNeasy Plus kit (Qiagen) according to the manufacturer’s guidelines. RNA was reverse transcribed using cDNA EcoDry Premix (Takara), and 1 uL of cDNA (10 ng of starting RNA) was amplified using the iTaq Universal SYBR Green Supermix (Bio-Rad) and readout with the Bio-Rad CFX96 qPCR system. Gene expression levels were calculated using the ΔΔCt method by first normalizing target gene Ct values to *GAPDH* within each sample and then comparing to the cell line iNPC12.

For qPCR, DNA was extracted from cells using the DNeasy blood and tissue kit (Qiagen) according to manufacturer’s guidelines. 10 ng of DNA was amplified using the iTaq Universal SYBR Green Supermix (Bio-Rad) and readout with the Bio-Rad CFX96 qPCR system. Relative DNA copy number was calculated using the ΔΔCt method by normalizing target gene Ct values to *GLI3* (a chromosome 7 reference locus) within each sample and then comparing to the diploid cell line iNPC12.

### smFISH on metaphase spreads or interphase cells

As described previously,^19^ one million cells were seeded into a single well of a 6-well plate coated with Geltrex (Corning, 354277) and, after one day, incubated in 100 ng/mL of Karyomax (Gibco) in standard growth media (see Methods above) for 10 hours. After incubation, cells were washed with PBS, lifted with Accutase (Innovative Cell Technologies), and spun down in the centrifuge at 1000 rpm for 5 minutes. The supernatant was aspirated, and the cell pellet was dislodged by gently flicking the conical tube. Cells were resuspended and incubated in 1 mL of warm 0.075 M KCl for ten minutes, after which 200 uL of cold Carnoy’s Fixative (3:1 mixture of methanol and acetic acid) was added and mixed by gentle inversion. Cells were then spun down at 1000 rpm for 5 minutes, the supernatant was aspirated, and the pellet was resuspended in 1 mL of cold Carnoy’s Fixative kept on wet ice. The cell suspension was transferred to a 1.5 mL Eppendorf tube and re-spun at 1000 rpm for 5 minutes. After aspirating the supernatant, fixed mitotic cells were resuspended in 40 uL of Carnoy’s fixative and were stored at -20C or used immediately for metaphase spreads. For each metaphase spread, 4 uL of cells were added to glass coverslips in a dropwise manner at a height of 6 inches.

For smFISH, fixed cells in interphase or arrested in metaphase were equilibrated in 2X SSC and dehydrated in a 70%, 85%, and 100% ethanol series for 2 minutes each. FISH probes were reconstituted in hybridization buffer (Empire Genomics) per the manufacturer’s guidelines, added to the slide directly atop cells, and covered with a coverslip sealed with clear nail polish. The glass slide containing the cells and FISH probes was placed on a 75C hotplate for 3 minutes to denature DNA and then placed into a humidified 37C incubator overnight (16-20 hours) for hybridization. The coverslip was removed, and cells were washed 2X for two minutes: once with 0.4X SSC buffer containing 0.3% IGEPAL and once with 2X SSC containing 0.1% IGEPAL. After washing, cells were mounted with ProLong Diamond Antifade Mountant with DAPI (Invitrogen) and a coverslip before imaging with a BZ-X800 microscope (Keyence).

### Spinning disc assay

Spinning disc assay was performed as described previously.^63^ 25 mm glass coverslips were coated with 2 μg/cm^2^ human fibronectin (isolated from serum), or GelTrex (Corning, 354277) according to manufacturer’s guidelines for 1 hour at room temperature or overnight at 4 °C. Cells were seeded onto the coated coverslips at a density of 2,037 cells/cm² to minimize cell-cell interactions. After plating, cells were allowed to attach for 48 hours at 37 °C. After the attachment period, the coverslips were mounted on a custom-built spinning disk device and submerged in D-PBS (Dulbecco’s Modified Eagle Media; Gibco, 11966025) supplemented with 4.5 g/L dextrose (Fisher Scientific, BP350-1), pre-warmed to 37°C. This spinning buffer lacked cations. The mounted cells were subjected to varying levels of fluid shear stress, determined by the rotational speed of the disk, for a duration of 5 minutes. After spinning, cells were fixed using 3.7% formaldehyde (Polysciences, 18814-10). Nuclei were then stained with 4′,6-diamidino-2-phenylindole (DAPI; ThermoFisher Scientific, D1306; 1:2000) and mounted on slides using Fluoromount-G (Southern Biotech, 0100-01).

Imaging was performed using a 10X objective lens on a BZ-X800 microscope (Keyence). Stitched images of the nuclei were generated using the default Keyence image capture software. A custom Python script was employed to analyze these stitched images and calculate the average cell adhesion strength (τ_50_). The τ_50_ value was defined as the shear stress at which 50% of the initially attached cell population was detached, following the equation:

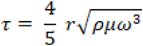

where *r* is the radial distance from the disk’s center, ρ is the density of the buffer, μ is the buffer viscosity, and ω is the rotational velocity.^52^ The Python code used for image analysis is available on the Engler Lab GitHub page (https://github.com/englea52/Englerlab).

### 10X-Multiome Library Generation

Collection of GBM specimens was approved by the Institutional Review Board at the University of Minnesota (IRB #STUDY00012599). Tumor samples were flash-frozen in liquid nitrogen immediately after surgical resection and stored at −80 °C until processing for 10x Genomics Multiome analysis. Samples were thawed and nuclei were isolated as previously described.^77^

Isolated nuclei were processed using the 10X Genomics Chromium Single Cell Multiome ATAC + Gene Expression assay following the manufacturer’s protocol, targeting recovery of 5,000–10,000 nuclei per sample. Sequencing library quality was assessed on an Agilent 4200 TapeStation using High Sensitivity ScreenTape, and library concentrations were quantified by real-time PCR. Libraries were sequenced on an Illumina NextSeq 2000 platform.

### mm-MERFISH

#### Oligonucleotide Probe design

For the mm-MERFISH experiments, we designed probes targeting the genome and transcripts for sequential imaging. Probes targeting the transcripts of central nervous system cell type markers were designed for combinatorial imaging.

##### 1) Sequential probes

To enable chromatin tracing, we designed sequential probes targeting 252 regions that are 5-Kb segments for the ecEGFR amplicon (chr7:54.9 - 56.2 Mb, genome build T2T-CHM13), based on public sequencing data.^15^ To delineate the endogenous chr7 locus, we targeted three additional regions that are specific to the endogenous chr7 locus. These regions are 1) 50kb downstream (chr7: 56,209,203 - 56, 259, 203) and 2) 100kb downstream (chr7: 56, 259, 203 - 56, 309, 203) of the ecEGFR amplicon, and 3) the 27kb EGFR vIII (exon 2-7 deletion) (chr7: 55,287,159 - 55,314,911).

In addition to harboring the oncogene *EGFR*, we designed probes to target eight additional coding genes encoded by the ecEGFR amplicon: *EGFR*, *EGFR-AS1*, *LANCL2*, *FKBP9P1*, *SEPTIN14*, *ZNF713*, *MRPS17*, *NIPSNAP2*, *PSPH*. We designed probes that target the intronic and exonic regions of each target region for ecDNA-encoded genes.

Encoding probes were designed using a publicly available pipeline (https://github.com/deprekate/LibraryDesigner), as described previously.^78^ For each region of interest, roughly 100 probes were designed per 5-Kb locus. Each probe has 40-nt complementary to the ecEGFR amplicon genome or transcripts, and the 40-mer region was screened to remove regions that are highly homologous to other targets or binding to a highly repetitive region on the genome. Each 40-mer was appended with three 20-nt adapter sequences unique to each 5-Kb locus, along with universal PCR priming sites at both ends.

A full list of encoding probes can be found in Supplementary Table 1 and 2.

##### 2) cell type markers

Additionally, to characterize the cell types within brain tumor tissue, we included an additional gene panel of 343 genes in the human central nervous system and 250 murine central nervous system cell type markers for the mouse xenograft models (Supplementary Table 1).

##### 3) Codebook design

For combinatorial imaging of murine cell type marker genes, we used a 48-bit, 4-ON-bits, hamming weight 4 barcoding scheme (Supplementary Table 1).

For combinatorial imaging of human cell type marker genes, we used a 60-bit, 4-ON-bits, hamming weight 4 barcoding scheme (Supplementary Table 1).

#### Probe Synthesis

The probes were generated from pooled oligonucleotides and processed in a manner similar to previously reported protocols.^78^ In brief, the custom oligonucleotide pools (Twist Biosciences) were subjected to limited-cycle PCR amplification, during which a 5′-acrydite–modified forward primer was incorporated to facilitate subsequent integration into an acrylamide gel. The resulting amplified DNA served as the template for in vitro transcription using T7 polymerase (NEB), followed by reverse transcription with Maxima RT (Thermo Scientific) and hydrolysis of the RNA strand to produce single-stranded DNA probes. These probes were then purified using Zymo Research D4003 and D4006 kits.

#### Readout and adaptor probe preparation

Adaptor probes and readout sequences are prepared in a similar fashion, as previously described,^22,78^ with the full list available in Supplementary Table 2. In addition to binding the encoding probe, each 60-nt adapter probe will bind two readout probes through a 20-nt complimentary sequence. Each readout probe is labeled with fluorescent dyes specific to the imaging channels: Cy3 (561nm), Cy5 (638nm), and Alexa Fluor 750 (750nm). These were purchased from IDT with high-performance liquid chromatography (HPLC) purification.

#### MERFISH sample preparation and hybridization

Sample preparation largely followed the protocol used for MERFISH+.^78^ Briefly, OCT-embedded tissues—mouse brains, patient-derived tumors—were sectioned at −20 °C using a Leica CM3050S cryostat to obtain serial coronal sections of 10-20 μm thickness. Samples were then lightly fixed in 4% PFA for 10 minutes and either frozen in 1x PBS with 10% glucose solution or immediately used for hybridization. Thawed or fresh samples were washed 2x with 1X PBS and then permeabilized using 0.5% Triton X-100 in 1× PBS for 10 minutes at room temperature. The tissues were then preincubated in a hybridization wash buffer (40% (vol/vol) formamide in 2× SSC) for 10 minutes at room temperature. After permeabilization, coverslips were transferred to a clean 60-mm petri dish and incubated with 0.1 M hydrochloric acid for exactly 5 minutes at room temperature, then washed three times for 5 minutes each with 1× PBS Tissue sections were then overlaid with 100 μL of encoding probe hybridization buffer containing 2× SSC, 50% (vol/vol) formamide, 10% (wt/vol) dextran sulfate, and a total encoding probe concentration of 1 μg/μL. Petri dishes with the sample and hybridization buffer-probe solution were then sealed with pressure adhesive (Blu-tack), and incubated in a water bath at 90°C for exactly 3 minutes and 30 seconds. Samples were then immediately transferred to a humidified oven at 47 °C for 18–24 hours. The next day, samples were washed in 40% (vol/vol) formamide in 2× SSC with 0.1% Tween-20 for 30 minutes at room temperature and then embedded into a polyacrylamide gel.

To immobilize the RNAs, the encoding-probe–hybridized samples were embedded in a 4% polyacrylamide gel for 6 hours, post-fixed with 4% (vol/vol) paraformaldehyde in 2× SSC, and washed with 2× SSC. Samples were then photobleached for 6-10 hours using a MERSCOPE Photobleacher (cat. #10100003, Vizgen). mm-MERFISH imaging was performed on a home-built microscope system as previously described and detailed below.

If RNA-MERFISH was being performed all solutions and buffers contained RNAse inhibitors until after the gel embedding step.

#### Image acquisition and experimental setup

The experimental setup comprised multiple integrated components, all of which are the same as described previously.^78^

Briefly, Image acquisition was performed using a custom-built fluorescence microscope. The core of the setup was a ASI microscope body fitted with a Nikon CFI Plan Apo Lambda 60x oil immersion objective (NA 1.4).

Automated buffer exchanges on the microscope stage were performed using a custom-built fluidics system. This system comprised a syringe pump (Hamilton), two 24-port valves (VICI C25G24524UMTA), a flow chamber (Bioptechs 060319–2), and tubing sealed with pressure adhesive (Blu-tack). Each valve was connected to a separate buffer source, with dedicated lines designated for imaging, stripping, and washing. Buffers were delivered into the sample chamber, and waste was collected downstream in an open-loop configuration.

Custom software was employed to coordinate and control these components and can be found at https://github.com/ZhuangLab/storm-control.

#### multi-modal MERFISH imaging

We first carried out sequential hybridization of fluorescent readout probes, followed by sequential imaging of the targeted genomic loci and the transcripts of genes on the ecEGFR amplicon. Then, we imaged the mRNAs for CNS cell type markers.

Finally we performed a series of antibody staining and imaging steps. For each sample, we imaged in the following order, using a 3 color imaging scheme:

1. 84 rounds of hybridization for chromatin tracing imaging, sequentially targeting 252 loci on the ecEGFR amplicon
2. 1 round of additional hybridization, sequentially imaging chr7 specific probes
3. 9 rounds of sequential imaging for the intron and exons of the gene on ecEGFR
4. 20 rounds of combinatorial imaging for 343 human CNS genes
5. (mouse xenograft only) 16 rounds of combinatorial imaging for 250 mouse CNS genes
6. 3 rounds of antibody imaging. Described more in depth below.

#### Immunofluorescence staining for mm-MERFISH

Antibody imaging was carried out immediately following completion of the DNA and RNA imaging. Three buffers/solutions were used for IF staining: (1) a blocking buffer consisting of 0.5% Triton-X in 1X PBS, (2) primary antibody solution of 0.3% Triton-X in 1X PBS with a 1:500 dilution of the targeted antibody, and a (3) secondary antibody solution 0.3% Triton-X in 1X PBS with a 1:500 dilution of the secondary antibody. Samples were initially incubated for 4 hours at room temperature with two primary antibodies derived from the host species (mouse/rabbit). After washing in 2× SSC for 15 minutes, the samples were then incubated for 2 hours with species-specific secondary antibodies conjugated to fluorescent dyes.

Primary antibodies used:

Pol2S2: Rabbit Anti-RNA polymerase II CTD repeat YSPTSPS (phospho S2) antibody (Abcam, Cat# ab193468, RRID:AB_2905557)

SC35: Mouse monoclonal [SC-35] to SC35 - Nuclear Speckle Marker (Abcam, Cat# ab11826, RRID:AB_298608)

H3K27ac: Rabbit Histone H3K27ac antibody (pAb) (Active motif, Cat# 39133, RRID:AB_2561016)

Lamina A/C: Mouse Lamin A/C (E-1) (Santa Cruz Biotechnology, Cat# sc-376248, RRID:AB_10991536)

Nup98: NUP98 (C39A3) Rabbit mAb (Cell Signaling Technology, Cat# 2598, RRID:AB_2267700)

Secondary antibodies used:

Anti-mouse secondary: Alexa Fluor® 790 AffiniPure Donkey Anti-Mouse IgG (H+L) (Jackson ImmunoResearch Labs, Cat# 715-655-150)

Anti-rabbit secondary: Alexa Fluor® 647 AffiniPure Donkey Anti-Rabbit IgG (H+L) (Jackson ImmunoResearch Labs, Cat# 111-605-144)

### Overview for mm-MERFISH analysis pipeline

The image analysis pipeline used for this was implemented and can be found at https://github.com/BogdanBintu/NMERFISH and https://github.com/deprekate/MERMAKE/

#### Flat-field correction

Images were initially processed to correct for nonuniform illumination across the field of view (FOV), particularly at the edges and corners, using flat-field correction. This was achieved by computing the per-pixel median intensity from the first imaging round for each color channel separately. Each image was then normalized by inversely scaling it according to the corresponding median brightness.

#### Deconvolution

The images were next deconvolved using a Wiener algorithm with a custom point spread function (PSF) computed for the specific microscope used for imaging, in order to reduce noise and sharpen fluorescent spot signals. A high-pass filter with a blur sigma of 30 was then applied to further enhance the signal-to-noise ratio.

#### Localization of fluorescent spots

Following image pre-processing, local intensity maxima within a 1-pixel radius were detected and subsequently filtered to require a minimum brightness of 3600 and a correlation of at least 0.25 with the point spread function. The remaining local maxima were retained for decoding.

#### Sequential FISH quantification

For transcripts imaged individually as smFISH targets, the fluorescent spots identified in the previous step were further filtered based on a minimum brightness threshold relative to their local background. This threshold was determined independently for each color channel and manually adjusted to eliminate dim and noisy signals while preserving bright puncta.

#### Chromatin tracing chromatic aberration

Chromatic aberrations between imaging channels were quantified and corrected by independently labeling the same set of genomic loci in each color channel. Signals corresponding to identical loci were then compared across channels to estimate channel-specific offsets and distortions, which were subsequently applied to correct inter-channel misalignment.

#### RNA-MERFISH decoding

To reconstruct RNA molecules from fluorescent spots detected across imaging rounds, spots were first grouped across images into clusters within a 2-pixel radius after drift correction. Because the MERFISH codebook was designed with a Hamming weight of 4, only clusters containing at least three spots were retained to permit single–bit error correction, while clusters with one or two spots were excluded. For each retained cluster, a brightness vector was assembled from the spot intensities in each round, L2-normalized, and compared against the codebook for decoding.

To remove noise from the decoded transcripts, a quality score was computed for each transcript using three criteria: (1) the mean brightness of the fluorescent spots, (2) the distance between the brightness vector and the closest entry in the MERFISH codebook, and (3) the average distance of each fluorescent spot from the cluster median. These metrics were combined into a single score by calculating a joint Fisher p-value based on the distributions of each metric across all decoded transcripts. The score distribution for transcripts assigned a blank barcode was then compared with that for transcripts assigned a gene identity, and a threshold was selected that separated the peaks of the two distributions.

#### Nuclei segmentation

For all experiments, the nuclei were segmented in three dimensions (3D) using Cellpose v2.^79^ For each z-slice, a 2D segmentation mask was first generated from the DAPI channel using Cellpose’s “dapi” model, following flat-field correction and deconvolution of the DAPI images as described above. Cells were then linked across neighboring z-slices to construct a 3D segmentation.

#### Cytoplasm segmentation

For the *in vitro* experiments, additional cytoplasm segmentations were generated based on a cumulative RNA signal. We pulled together drift-corrected images from the sequential smFISH for genes encoded on the ecDNA. The max projection of all rounds were used as the ‘cyto’ channel, while the DAPI remained as the ‘dapi’ channel. The cytoplasm segmentation was done using Cellpose.^79^

#### Assigning transcripts and chromatin traces to cells

Transcripts detected by both smFISH and MERFISH were assigned to individual cells by first correcting the coordinates of each molecule using the computed drift between the segmentation images and the images in which the RNA were acquired. The adjusted coordinates were then rounded to the nearest integer, and each transcript was assigned to a cell according to the value of the segmentation mask at its corresponding coordinates.

### **EcDNATracer:** Reconstruction and assignment of single-molecule ecDNA traces

We developed a computational pipeline ‘ecDNATracer’ to reconstruct ordered single-molecule ecDNA traces from multiplexed DNA-MERFISH spot detections and assign these traces to endogenous chr7 or ecDNA based on chr7 probes, producing standardized **linear** and **circular** trace tensors of shape (N, M, 6) with default M = 252 loci.

For each cell, the input consisted of a spot table X ∈ R^{P × 6}, where each row corresponded to a detected locus with 3D position (z, x, y), log-intensity log H, color channel c, and locus (round) index r. Traces were reconstructed as ordered sequences of detections spanning locus indices and subsequently assembled into standardized tensors in which each trace occupied a fixed M × 6 array with missing loci filled by NaNs.

To generate candidate traces, we first constructed a proximity graph over detections using a KD-tree radius query with threshold d_th (default d_th = 0.75um). Edges were formed between detections i,j satisfying ||x_i − x_j||_2 ≤ d_th and annotated with spatial distance d_ij and genomic gap Δr_ij = |r_i − r_j| (wrapped for circular tracing). Edges were weighted by a decreasing function w(d, Δr) and processed in descending order using a union–find structure that merged components greedily under a locus-uniqueness constraint, favoring components resembling valid polymer traces.

Each coarse trace was locally refined by augmenting nearby detections within radius D, retaining the brightest candidates per locus, and gating by locus-specific intensity thresholds. Refined candidates were re-traced using a layered-graph formulation in which nodes represented detections at each locus and edges connected spatially proximal detections across loci. Edge costs combined spatial distance and genomic gap via a log-score function, with optional brightness priors. Greedy extraction of disjoint minimum-cost paths yielded one or more reconstructed traces per candidate set.

To assign traces to genomic anchor points, we evaluated each trace against three predefined anchor sets (default loci 73, 253, and 254) by combining spatial and genomic distances using a scoring function. One-to-one anchor–trace matching was solved using the Hungarian algorithm. Anchor superpoints were constructed by intersecting consistent anchor matches across sets, and joint ASP–trace assignment was solved by Hungarian matching with optional distance gating.

Selected traces were assembled into standardized linear and circular tensors (N, 252, 6), constituting the final output for downstream structural and transcriptional analyses.

See **[Code Availability]** for the ecDNATracer github link containing all code to perform ecDNA chromatin tracing.

#### Simulator

ecDNATracer comes with a companion simulator. To model experimental noise in reconstructed traces, we implemented a generative simulator that perturbs an underlying “true” trace using empirically motivated stochastic processes to mimic experimental noise. Starting from a set of true 3D locus coordinates X = { x₁, …, x_N }, the simulator generates an observed trace Y by introducing three independent noise components: (i) locus dropout, (ii) spurious detections, and (iii) spatial localization error. First, each true locus is retained with probability p_det, modeling missed detections as a Bernoulli process. Second, a random number of spurious loci is added, with the count drawn from a Poisson distribution to represent background or misassigned detections. Third, all retained and spurious loci are perturbed by additive isotropic Gaussian noise to model localization uncertainty. The resulting observed trace Y thus reflects realistic detection inefficiency, background contamination, and coordinate jitter, while preserving the large-scale geometry of the original trace.

### Detection efficiency analysis

Detection efficiency for each imaging round was defined as the percentage of traces in which the corresponding 5-Kb locus was detected for that experiment.

### Copy number analysis

Copy number was computed by taking the average number of spots over a set number of rounds with different, progressively inclusive strategies: (1) The first two loci exhibited higher detection efficiency and reduced cumulative signal loss compared to later imaging rounds. As such, we computed the mean number for the first two loci. This was used in patient-derived tissue. (2) In addition to the first two loci, we computed the mean number of spots across the promoters of EGFR and EGFR-AS1 (four additional loci total), which would normally be targeted with standard EGFR DNA FISH. This was used for *in vitro* and Mouse Replicate 2 experiments. (3) For an exhaustive method, we used the mean number of spots across all 252 loci. This method was used for Mouse Replicate 1. All three methods strongly correlated with each other across replicates; e.g., For GBM39 *in vitro*, (1) vs. (2): Pearson r = 0.94; (1) vs. (3): Pearson r = 0.97; and (2) vs. (3): Pearson r = 0.94.

### In vitro analysis

#### Chromatin tracing *in vitro*

We jointly profiled 655 cells across two technical replicates of GBM39 cells, capturing 48,460 ecDNA traces and 1,063 endogenous chromosomal EGFR loci. For each experiment, roughly 40 field of views (FOVs) were selected, avoiding densely seeded areas. Starting from the acquired images, we ran through the entire mm-MERFISH processing pipeline for preprocessing, and used ecDNATracer to identify individual ecDNA and matching endogenous chromosomes.

#### Proximity Frequency Matrices from Imaging

We then computed the median pairwise distance across all traces to get the representative domain structures in the ecDNA amplicon. For validation, we computed the contact frequency from published HiC data in the same cell line in the corresponding genomic region.^19^ To compute the proximity frequency for any given pair of loci, we first counted the number of measured distances for that locus pair that were below a defined cutoff distance (250 nm in this study, unless otherwise specified). This count was then normalized by dividing it by the total number of distances measured for that pair of loci to produce a contact frequency rate. The proximity frequency matrix was used in comparison with the 5-Kb contact frequency matrix produced by HiC.

#### Single trace visualization

Individual assembled traces were visualized in three dimensions using a ball-and-stick representation with a continuous rainbow colormap assigned along genomic order. Each detected 5-Kb locus was rendered as a spherical node, and consecutive loci were connected by a smooth interpolating curve obtained by fitting a spline to the ordered 3D coordinates. Missing intermediate loci were bridged implicitly by the spline interpolation to produce a visually continuous trajectory. For circular ecDNA traces, a straight closing edge was added between the last and first loci with a linearly interpolated color gradient to indicate circular continuity. The first locus (chr7: 54.925–54.930-Mb) was colored purple and the last locus (chr7: 56.207–56.212-Mb) was colored red. All visualizations were rendered in Napari.

#### Radius of gyration

To quantify the spatial compaction of each assembled trace, we computed the radius of gyration (Rg) from the 3D coordinates of detected loci. For a trace containing N loci, we first subtracted the center of mass of the trace from each coordinate and then calculated the root mean square distance of loci from this center. Rg was computed separately for linear and circular traces using the same definition. Missing loci were excluded from the calculation.

#### Theoretical radius-of-gyration modeling

To model the radius of gyration (Rg) under realistic experimental noise, we generated synthetic circular (ecDNA) and linear (chr7) polymers using the ecDNATracer simulator, incorporating detection dropout, Poisson-distributed false positives, and localization noise. Rg was computed from each simulated trace using the same definition as for experimental data. We simulated equal numbers of circular and linear traces, respectively, to match the empirical sampling, yielding a balanced, noise-aware null baseline for interpreting observed compaction differences.

#### Linking chromatin traces to nascent transcription

The nascent RNA of the 9 genes were quantified the same way as described above in‘smFISH signal quantification’. For each cell and intron species, detected intron molecules were assigned to reconstructed traces by enforcing a one-to-one spatial matching. For each intron–trace pair, we computed the mean Euclidean distance between the intron molecule and all detected loci of the trace, ignoring missing loci, yielding a distance matrix Di,t. Assignments were obtained using the Hungarian algorithm (linear sum assignment) with the constraint that each intron molecule could be assigned to at most one trace and each trace to at most one intron molecule. A hard spatial gate Rmax (default 2.0 µm) was applied such that intron–trace pairs with Di,t > Rmax were disallowed, allowing intron molecules to remain unassigned if no proximal trace was present. Assignments were performed jointly across circular (ecDNA) and linear (chr7) traces and reported separately for each type.

#### Bursting rate quantification

For each intron species, we classified whether a reconstructed trace was transcriptionally bursting based on the brightness of the associated intron molecules, after correcting for color-specific intensity differences. Baseline brightness statistics were first estimated separately for each hybridization color by subsampling cells and pooling all intron-molecule brightness values within each color, yielding a per-color median m_c and standard deviation s_c. For a given intron species g (mapped to color c) and trace t, the trace-level intron brightness H_{t,g} was converted to a standardized score

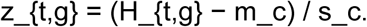

A trace was classified as bursting for intron g if z_{t,g} exceeded a predefined color-specific threshold t_c (t_c = 1 for color 0 and t_c = 2 for colors 1 and 2), yielding a binary indicator per intron species. The total number of bursting introns per trace (‘total_genes_transcribed’) was computed as the sum across intron species.

#### Local antibody brightness quantification

For each antibody channel and field of view (FOV), raw immunofluorescence images were flat-field corrected and a high-pass filter with a blur sigma of 20 was then applied to further enhance the signal-to-noise ratio. Corrected images were then normalized within each FOV by z-scoring using the image-wide median and standard deviation. For each reconstructed trace, normalized intensities were sampled at each locus position of each trace; loci with missing coordinates or out-of-bounds indices were not considered in the quantification. A single antibody brightness per trace was defined as the mean across all detected loci, yielding a per-trace normalized brightness value comparable across FOVs and pooled across chr7 (linear) and ecDNA (circular) traces.

#### Global antibody brightness quantification

For each antibody channel and FOV, raw immunofluorescence images were flat-field corrected and rescaled to preserve the global intensity level. Corrected images were then normalized within each FOV by z-scoring using the image-wide median and standard deviation. For each cell, the average global brightness per antibody was calculated as the mean across all pixels that lied in the nucleus segmentation for that cell.

#### Quantifying Intermolecular heterogeneity

Intermolecular heterogeneity across reconstructed traces was quantified using pairwise distances of standardized feature values. For each trace t and feature f (total genes transcribed, radius of gyration, Pol II brightness), values were globally z-scored across all traces. For each feature,

<u>within-cell distances</u> were defined as d_within = | z_{t1,f} − z_{t2,f} |

for all trace pairs (t1, t2) originating from the same cell, and

<u>between-cell distances</u> were defined as d_between = | z_{t1,f} − z_{t2,f} |

for trace pairs originating from different cells. To avoid over-representation of cells with many traces, the number of within-cell pairs per cell was capped by random subsampling, and the number of between-cell pairs was matched to the within-cell sample size. For each feature, within- and between-cell distance distributions were compared using a two-sided Mann–Whitney U test, testing the hypothesis that within-cell distances are smaller than between-cell distances.

#### Active vs. repressive marks across ecDNA

For each antibody mark, we first computed the local brightness level for each 5-Kb locus for each trace. We then computed the relative brightness for each trace by z scoring the brightness within each trace. To compare active versus repressive chromatin compartments, we first summarized each antibody by computing a per-locus average across traces and then aggregated marks within functional groups by taking the per-locus median across active marks (PolII, H3K27Ac, spliceosome) and across repressive marks (lamin, Nup98).

#### Lamina association analysis

To estimate distances to the nuclear lamina, we used the nuclear segmentation boundary as a geometric proxy for the lamina surface. Specifically, inner boundary voxels were aggregated into a per-cell boundary point cloud. The distance to the lamina for each trace was defined as the Euclidean distance from the trace centroid to the nearest boundary point, computed using a KD-tree nearest-neighbor search. To be considered “lamina associated”, the distance to lamina has to be smaller than 750 nm, which is the average radius of gyration of a single ecDNA molecule.

#### Speckle association analysis

Nuclear speckles were defined from normalized SC35 immunofluorescence images by Gaussian smoothing (σ = 0.1 µm), thresholding at intensity > 8, and removal of connected components smaller than 10 voxels. The binary speckle mask was converted to a point cloud in physical space after anisotropic voxel scaling. For each ecDNA trace, centroids were drift-corrected and the minimum Euclidean distance to the nearest speckle voxel was computed and used as the distance-to-speckle metric. The association analysis is the same as ‘lamina association analysis’.

#### Micronuclei analysis

The micronuclei (MN) segmentation is done with a human-in-the-loop Cellpose 2.0^79^ training, where the data was compiled as the max projection of the lamina and DAPI image. The final segmentation was improved upon human review.

A cell was considered containing MN (MN+ cell), if the MN segmentation falls within the cytoplasm but outside the main nucleus.

A MN+ cell is considered having ecEGFR DNA, if the selected 6 loci (2nd copy number method) were all considered to be in the cell.

A MN+ cell is considered having EGFR intron, if the smFISH EGFR intron signal falls in the MN segmentation.

### Mouse analysis

#### Species identification using MALAT1/malat1 and species-specific gene expression

Mean brightness values for *MALAT1* and *Malat1* were computed for each cell, and cluster-level mean brightness values were calculated for each Leiden cluster, revealing two groups corresponding to human and mouse cells. For initial species classification, clusters with mean Malat1 brightness values <4500 were classified as human, whereas clusters with values >4500 were classified as murine. Mean human and murine gene counts were assessed for each cluster, which aligned with initial Malat1/MALAT1-based annotations, with the exception for a small number of edge cases. To account for contributions from neighboring cell types due to technical reasons (e.g., segmentation biases), we performed background subtraction by identifying, for each cell, the 50 nearest neighboring cells belonging to other clusters within a 30 μm radius and calculating their mean gene expression profile. This neighbor-derived expression profile was scaled to the total number of neighbors within the radius and subtracted from each cell’s gene expression vector. Cluster-level mean human and murine gene expression was then reassessed, resulting in improved separation between human and murine clusters. Clusters were subsequently subsetted based on their species, genes corresponding to the opposite species were removed, and cells were reprocessed independently within each species (e.g., dimensionality reduction and clustering).

#### GBM cell state assignment

GBM cellular states were computed and assigned as done previously.^19,33^ Briefly, we quantified the gene enrichment score for each GBM state gene set.^33^ Each score was calculated as the relative average expression of the gene set compared to a control gene set matched for similar baseline expression levels. After obtaining scores for all four cellular states, cells were first stratified into NPC-/OPC-like or AC-/MES-like using a differential score: D = max(SC_OPC_, SC_NPC_) - max(SC_AC_, SC_MES_). For distinguishing a singular state, an identity score was calculated as C = log_2_(|SC_OPC_ - SC_NPC_| + 1) for NPC-/OPC-like cells and C = log_2_(|SC_AC_ - SC_MES_| + 1) for AC-/MES-like cells. Each cell was then plotted within a four-quadrant scatter plot with D on the y-axis and C on the x-axis. For bulk RNA-sequencing, ssGSEA scores were used to calculate D and C instead of single-cell gene enrichment scores.

#### RNA sequencing analysis of public GBM cohorts

We compared *SEMA5A*, *SOX8*, and *LINGO1* expression across a cohort of 10 GBM tumors that were multi-regionally sampled using the 3D GBM dataset and interactive browser.^51^ TCGA data was accessed through cBioPortal.^80–82^ Briefly, gene expression values were obtained for 145 GBM tumors from the TCGA PanCancer Atlas.^83^ Each gene was correlated with *SEMA5A*, *SOX8*, and *LINGO1* gene expression. Genes were then ranked by their Spearman correlation coefficient, and the top 100 positively correlated genes were used as input for Enrichr^84^ gene set enrichment analysis to identify biological processes associated with genes strongly correlated with 6_OPC marker genes. All genes detected by RNA sequencing were used as a background gene set for Enricher analysis.

#### PCA and rank-rank analysis

Malignant cells were subsetted, and PCA was recomputed using the algorithm from Scanpy.^52^ We correlated the expression of each gene in our 343 MERFISH gene panel with EGFR expression and/or ecDNA copy number across all cells, as indicated in the figure legend. All genes were then ranked based on their Pearson correlation such that genes with strong positive correlations (high positive Pearson R values) received low rank values, whereas genes with strong negative correlations received high rank values. Similarly, genes were ranked according to their PC1 loading scores, with strongly positive PC1 loadings assigned low rank values and strongly negative PC1 loadings assigned high rank values. For every gene, both ranks––i.e., the EGFR correlation rank and the PC1 loading rank––were plotted against and correlated with each other.

### Comparison of *in vitro* and *in vivo* GBM39 datasets

We used the Ingest function in Scanpy^52^ to project query datasets (GBM39 *in vitro* and GBM39 mouse replicate 2) into a shared reference transcriptional space defined by GBM39 mouse replicate 1 using default parameters. Prior to mapping, query datasets were re-normalized to the same number as Mouse Replicate 1 (median total counts of Mouse Replicate 1) to ensure consistent scaling and prevent normalization-induced bias during PCA projection. The reference dataset was preprocessed and embedded using principal component analysis (PCA), followed by neighborhood graph construction and UMAP visualization. The learned PCA loadings from the reference were used to project query cells into the same principal component space, and nearest-neighbor relationships were assigned relative to the reference. Query cells were subsequently embedded into the precomputed UMAP space without modifying the reference embedding.

#### ecDNA dispersion score

To quantify the spatial dispersion of the ecDNA molecules within each cell, we computed the root mean square (RMS) distance of the 1st 5kb loci from their centroid in three-dimensional space. To control for cell-specific geometry and sampling density, we generated a null expectation by randomly sampling n positions from the set of candidate genomic loci within the same cell and computing the same RMS dispersion metric; this was repeated 100 times and averaged. The dispersion score was defined as the ratio of the observed RMS dispersion to the simulated mean dispersion, such that values greater than 1 indicate broader spatial spread than expected by random sampling within the cell.

### Chromatin tracing *in vivo*

#### Chromatin tracing

The raw processing was the same as described in the section ‘Chromatin tracing in vitro’. After cell type identification, we randomly sampled 1500 cells from each cell cluster and performed chromatin tracing. The input traces were then used as input to ‘ecDNATracer’ with the same default parameters described above. A median distance matrix was computed for each GBM cluster.

#### Centrality Score

For each ecDNA trace, we quantified radial organization by computing a centroid-based centrality score for every detected locus. The 3D centroid of the trace was first calculated using all detected loci, and the Euclidean distance from each locus to this centroid was measured. Each locus was then assigned a normalized centrality score defined relative to the mean centroid distance across the trace, such that loci closer to the centroid than average receive higher scores. Traces were excluded if fewer than 60% of loci were detected. In this formulation, values approaching 1 indicate strongly central positioning within the ecDNA molecule, values near 0 indicate approximately average radial position, and negative values indicate loci that are more peripherally positioned than the trace average.

#### Structural variant identification

Based on the structural variant we discovered in the 4_MES population, we re-examined all ecDNA traces by screening reconstructed ecDNA polymers for contiguous genomic regions lacking detectable loci. For a specified locus window, in this case, chr7:54.925-55.000 Mb and chr7:55.977-56.207 Mb, we quantified the fraction of 5kb segments whose spatial coordinates were missing and classified a trace as structurally altered if ≥95% of loci within that window were undetected.

### Next generation sequencing analysis

#### Hi-C analysis

Single-cell Hi-C data were processed as previously described.^19^ Reads were mapped to the T2T reference genome to match the chromatin tracing probe coordinates. Briefly, data was mapped using bwa mem with flag ‘-SP5M’. Aligned Hi-C pairs were parsed and sortedw ith pairtools with parse flag ‘--min-mapq 40 --walks-policy all --max-inter-align-gap 30’. PCR duplicates were subsequently identified and removed using pairtools dedup with consideration of single cell barcodes. Contact matrices were then generated at 5-Kb resolution using cooler, and multi-resolution .mcool files were created and balanced with ‘cooler zoomify’ at resolutions of 5-Kb, 10-Kb, 25-Kb, 50-Kb, 100-Kb, 250-Kb, 500-Kb, 1-Mb, and 2.5-Mb. Cells were then aggregated to generate a pseudo-bulked contact matrix at 5-Kb resolution for comparison with ensemble chromatin tracing measurements of ecDNA.

#### RNA sequencing analysis

Paired-end 150 bp unstranded RNA-seq was performed by Novogene Corporation Inc. (Sacramento, CA). FASTQ file quality was assessed using FastQC, and reads were aligned to the hg38 reference genome (GENCODE v49 annotation) using STAR (v2.7.11a). Alignment quality was evaluated using STAR log files. BAM files were indexed with samtools (v1.19) and manually inspected in IGV (v2.19.7) to confirm mapping quality and transcript structure. Gene-level counts were generated using FeatureCounts from the Subread package (v2.1.1). Differential gene expression (DEG) analysis was performed in R (v4.5.2) using DESeq2. Genes with fewer than 10 total counts across all samples were excluded prior to analysis. Statistical significance was determined using Benjamini–Hochberg adjusted P values (Padj < 0.05). For gene ontology (GO) analysis, DEGs were further filtered to include genes with an absolute fold change ≥2. GO enrichment was performed using Enrichr. Heatmaps were visualized using the pheatmap package, and volcano plots were generated using the EnhancedVolcano package. Gene set variation analysis (GSVA) was performed using the GSVA package in R. Variance-stabilizing transformed (VST) counts generated by DESeq2 were used as input. Gene sets were derived from MSigDB or previously published gene signatures. Gene identifiers were standardized to uppercase, and gene sets containing fewer than 10 genes or fewer than 10 genes overlapping the expressed dataset were excluded. Single-sample gene set enrichment analysis (ssGSEA) was performed with normalization enabled. GSVA enrichment scores were Z-scored across or within cell lines, as indicated in the figure legends. Three replicates were included for each experimental condition.

#### Epigenomic sequencing analysis

Publicly available epigenomic sequencing data for GBM39 were accessed from Wu et al.^15^ and analyzed as follows. The sequenced ChIP-seq reads were aligned to the T2T assembly using Bowtie2 (v2.5.3) with default parameters. Samtools (v1.19.2) was used to sort and index the aligned BAM files. Peaks were called using MACS3 (v3.0.1), and normalized BigWig coverage files were generated using deepTools bamCoverage (v3.5.5) with an effective genome size of 3,117,292,070. 4C data analysis was performed using 4C-ker (https://github.com/rr1859/R.4Cker) with default parameters. Briefly, a reduced genome was built using the T2T genome and Csp6I restriction sites. FASTQ files were aligned to this reduced genome using Bowtie2 (v2.5.3). The resulting SAM files were used to generate BedGraph files, and self-ligated and undigested fragments were removed. A 4C-ker object was then created, and 4C interactions were called using the nearBaitAnalysis module with adaptive windows (k = 5). scanMotifGenomeWide.pl from HOMER was used to identify CTCF motifs genome-wide using relaxed odds ratio thresholds.

### Mouse neighborhood analysis

#### Nonmalignant cell type assignment

After subsetting nonmalignant cells based on species identification (see Methods above), cells were re-clustered using the Leiden^49^ algorithm and manually annotated based on canonical marker gene expression.^33,59,60^ Clusters with the same major cell type label were merged.

#### Invasion analysis

We computed the number of malignant and nonmalignant neighbors within a radius of 200 µm for each GBM cell and classified it as “invasive” if ≥75% of its neighbors were non-malignant (e.g., malignant fraction ≤25%).

### Co-culture analysis

#### Co-culture MERFISH experiment

GBM39 were co-cultured longitudinally with neurons and harvested at Day 0, 10, 20, and 30. At each time point, the same mm-MERFISH libraries described above were hybridized to the samples. We sequentially imaged selected markers in the following order: the first three 5-kb segments of ecEGFR; EGFR intron and exon transcripts; astrocytic markers VIM and TNC; and neuronal markers NEFL and NEFM.

#### Co-culture MERFISH preprocessing

For each DNA segment or gene, the preprocessing is the same as described in ‘Sequential FISH quantification’. Given that the GBM39 were sparsely seeded, we selected FOVs that have at least 3 GBM cells for downstream analysis.

#### Cell-type assignment and GBM cell selection

Cell types were assigned using a marker-based clustering approach combined with EGFR status. Marker expression counts were normalized within each cell (CPM-like normalization with a pseudocount), standardized across cells (z-scoring per gene), and clustered using k-means (k = 2) to separate major cellular populations. Cluster assignments were reconciled with EGFR-based classification to define neurons, GBM, and edge-case categories. For downstream analyses, we retained cells classified as GBM or GBM-like and further restricted the dataset to cells with ecDNA copy number between 3 and 200 and EGFR exon counts ≤ 1000 to harmonize dynamic ranges across datasets and reduce potential confounding from potentially technical outliers.

### Co-culture comparison metrics

*EGFR copy number*: Colocalization was evaluated using the first three 5-kb segments of ecEGFR across adjacent imaging rounds. Molecules detected within a 2-pixel radius in consecutive rounds were considered colocalized, and the total number of colocalized molecules was used as a proxy for ecDNA copy number.

*EGFR expression*: the nuclear EGFR exon counts.

*Dispersion score*: ‘ecDNA dispersion scorè was computed based on the colocalized points.

### 10X-Multiome analysis

#### Preprocessing

Raw sequencing data were demultiplexed and aligned to the human GRCh38 reference genome using Cell Ranger ARC (10x Genomics). Downstream analyses were conducted in R using Seurat^85^ and in Python using Scanpy.^52^ Briefly, low-quality cells were filtered based on the number of detected genes, total UMI counts, and mitochondrial gene content per cell. Data from all samples were then integrated and scaled while regressing out total UMI counts and the percentage of mitochondrial transcripts. Dimensionality reduction was performed using Uniform Manifold Approximation and Projection (UMAP) following principal component analysis (PCA) as implemented in Seurat.^85^ Unsupervised Leiden clustering was carried out using the graph-based clustering algorithm in Seurat.^85^

#### Cell type identification

Clusters were annotated based on expression of established marker genes and their copy number variation profiles, which were calculated using InferCNV^32^ and epiAneufinder^65^. Briefly, copy number variation profiles were assessed across clusters to infer malignant status. Specifically, canonical GBM genetic alterations (e.g., chromosome 7 gain and chromosome 10 losses) were used to distinguish malignant and non-malignant cells. Afterwards, marker gene expression was assessed from previously described sources^59,60^ to assign broad cell types for non-malignant cells. GBM states were assigned to malignant cells according to previously described methods.^19,33^

#### Identification of focal EGFR gains in 10X-Multiome data

To identify focal gains of EGFR, GC-corrected ATAC fragments were computed using epiAneufinder^65^ and aggregated within 500-Kb bins across each chromosome. For each bin, fragments were summed across all cells within each sample, and pseudo-bulked fragments for each bin were plotted for each chromosome and sample. Aggregating fragments across large genomic bins (≥500 kb) enables detection of broad copy number alterations rather than variability driven by local chromatin accessibility. Relative fragment distributions across bins within each chromosome were used to control for global ploidy shifts and arm-level copy number changes.

### 10X Visium analysis

#### Identifying focal gains of EGFR in 10X-Visium data

We used a similar approach to our 10X-Multiome analysis to identify samples with putative focal EGFR gains (consistent with EGFR-containing ecDNA). Specifically, genomic coordinates for each gene were obtained from Ensembl (GRCh38 assembly) using the BioMart package, and only genes mapping to chromosome 7 that were detected in the dataset were retained. We defined a ±1-Mb window centered on the EGFR transcription start site (2-Mb total span) and summed raw counts across all genes within this interval for each spot. To control for differences in sequencing depth, window-level counts were normalized by total counts per spot to compute counts per million (CPM), followed by log1p transformation. This yielded depth-normalized EGFR-locus counts per spot. For visualization and within-sample comparisons, aggregated EGFR-locus counts were additionally z-scored within each sample.

To identify samples with putative focal gains of EGFR, we restricted analysis to malignant spots as defined by CNA inference.^60^ For each sample containing spots classified as malignant (17/26), we summed counts across chr7 genes and computed the fraction of total chromosome 7 expression arising from the EGFR-centered window. Significance was assessed using a permutation-based null model in which contiguous genomic blocks of equal gene number were randomly positioned along chromosome 7 (25,000 permutations per sample). An empirical one-sided p-value was calculated as the fraction of permutations with window-level expression greater than or equal to the observed value. P-values were adjusted across samples using the Benjamini–Hochberg multiple-hypothesis testing correction procedure.

Nine of the 26 samples exhibited FDR-adjusted p-values < 0.05, but to ensure reliability of our results, we also considered absolute counts from that locus and the number of genes being highly expressed from the EGFR window. Specifically, to prevent single highly expressed genes from driving false positives, we also required that at least two genes from within the EGFR window were ranked within the top 1% of chromosome 7 genes by expression within that sample. Seven samples satisfying both permutation significance (FDR-adjusted) and focal-expression criteria were classified as harboring putative EGFR focal gains consistent with ecDNA amplification.

#### Defining invasion in 10X-Visium data

Adapting similar methods from Greenwald et al.,^60^ we defined 10X-Visium spots based on the identity of neighboring spots. Specifically, for each spot, we defined a neighborhood radius that spans approximately four spots (four times the median nearest-neighbor spacing within each sample). We then calculated the fraction of neighbors that derive from cell types from the brain parenchyma: neurons, oligodendrocytes, and (reactive) astrocytes. If these cell types comprised ≥30% of neighboring spots, then that spot was classified as infiltrative. All other spots were classified as non-infiltrative.

### Patient-derived GBM tissue analysis

#### Cell type and state assignment

First, cells were clustered using the Leiden algorithm and manually annotated based on canonical marker gene expression.^33,59,60^ While some clusters were assigned the same cell type and state, they possessed differences in gene expression and were labeled distinctly (e.g., TAMs - 1 and TAMs - 2); i.e., they were not merged into a broad classification. An adjacent tissue section profiled by 10X-Multiome provided sequencing-based annotations of cell types and transcriptional states, and correlation of mm-MERFISH clusters with 10X-Multiome populations largely confirmed our manual annotations. Two mm-MERFISH Leiden clusters that correlated poorly (Pearson r < |.3|) with 10X-Multiome populations were designated as unclassified. Some GBM populations correlated with multiple transcriptional states (e.g., AC-/MES- or NPC-/OPC-like). These populations were therefore annotated by multiple state labels to reflect this shared transcriptional identity.

#### GBM zone analysis

We started by defining the cell with the highest ecDNA copy number. Radial GBM zones were then assigned by computing the Euclidean distance from each cell centroid to this cell and partitioning these distances into linearly spaced concentric bins of equal distance. Each cell was thereby assigned to a discrete radial zone for downstream spatial analyses. Zone 1 defined the inferior edge of the tissue, while Zone 6 defined the superior edge of the tissue.

#### Chromatin tracing in patient-derived GBM tissue

Following identification of AC/MES-like GBM cells within the sample, chromatin tracing was performed on all cells in this subpopulation given their high ecDNA copy numbers. Because the probe set only partially tiled the ecDNA in this patient sample (chr7: 54.945 - 55.582-Mb), these loci were analyzed using the linear reconstruction mode in ecDNATracer. To delineate the endogenous chr7 locus, we targeted three additional 5-Kb loci downstream to the ecDNA. These regions are 1) chr7: 55,687,185-55,692,185 2) chr7:55,692,185-55,697,185 3) chr7:55,702,185-55,7071,85. Radius of gyration and other chromatin tracing analysis methods were implemented as described previously in the *in vitro* methods section.

### Quantification and statistical analysis

Sample sizes are reported in the manuscript, figure legends, or directly in the figure panels. The number of replicates was determined based on similar prior studies. Power studies were not performed because of unknown effect sizes during the experimental design phase of our study. All generated data was used for data analysis with the exception of single cell data in which we excluded based on technical features that indicate low quality cells: e.g., the number of detected genes, total UMI counts, and mitochondrial gene content per cell. Randomization and blinding was not performed for our studies.

## Data Availability

Data will be made available upon publication. Please reach out to the corresponding authors for data requests.

## Code Availability

Code will be made available upon publication. Please reach out to the corresponding authors for code requests.

## Acknowledgments

This work was supported by the National Human Genome Research Institute (4DN) UM1 HG011585 (B.R.); a SSCI Stellar Faculty Award, SSCI Brain Tumor Program and R01 NS134798 (F.F.); and the National Institute of Mental Health DP5-OD031878 and Chan Zuckerberg Initiative (CZI) AICP-0000000175 (B.B.). B.T. was supported by the following NIH grants: F30CA291021 and T32CA067754. W.D. was partially supported by NIH grant T32GM139790. Z.W. is a DDBrown Awardee of the Life Sciences Research Foundation.

## Author contributions

Study conception: B.T., W.D., T.K., B.R., F.F., B.B

Study supervision: B.R., F.F., B.B

mm-MERFISH data generation and analysis: B.T., W.D., T.K., K.M., M.N., B.B.

ecDNATracer: W.D., B.T., B.B.

GBM *in vitro* experimentation and analysis: B.T., T.K., Z.G., R.V., A.B., O.E., S.K., B.J., T.M.

GBM *in vivo* experimentation: B.T., Y.M., D.K.

Patient-derived GBM tissue collection: C.C. 10X

data generation: Z.W., T.L., L.C.

Next generation sequencing analysis: B.T., W.D., Z.W., J.B., B.S., Y.X., L.C.

Data interpretation: B.T., W.D., B.S., B.R., F.F., B.B.

Manuscript writing: B.T., W.D., B.R., F.F., B.B.

All authors edited and approved the final manuscript.

## Materials & Correspondence

Further information and requests for resources and reagents should be directed to corresponding authors. Lead contacts: Bogdan Bintu, Frank Furnari, and Bing Ren.

## Extended Data Figures

**Extended Data Figure 1.**
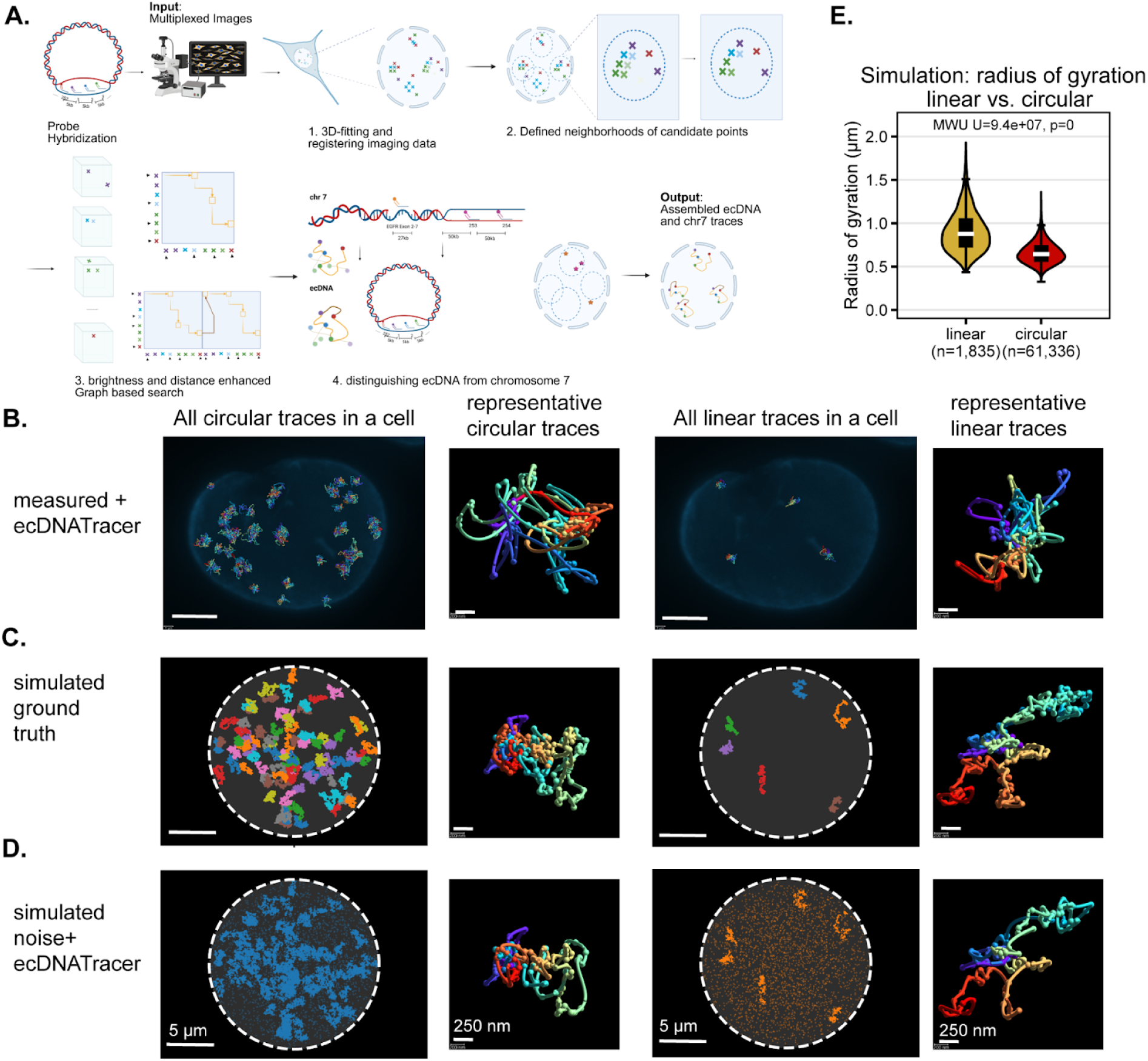
ecDNA chromatin organization defies polymer physics simulation results. **(A.)** Schematics of ecDNATracer. **(B.-D.)** Representative traces of ecDNA and chr7 obtained from GBM39 cells *in vitro* (B.), “ground truth” polymer simulations (C.), and reconstructions from ecDNATracer upon simulating noise and loss of detection efficiency (D.). **(E.)** Violin plots showing the radius of gyration of linear and circular simulated polymers. A two-sided Mann Whitney U test was used to test for statistical significance. P value < 0.0001.

**Extended Data Figure 2.**
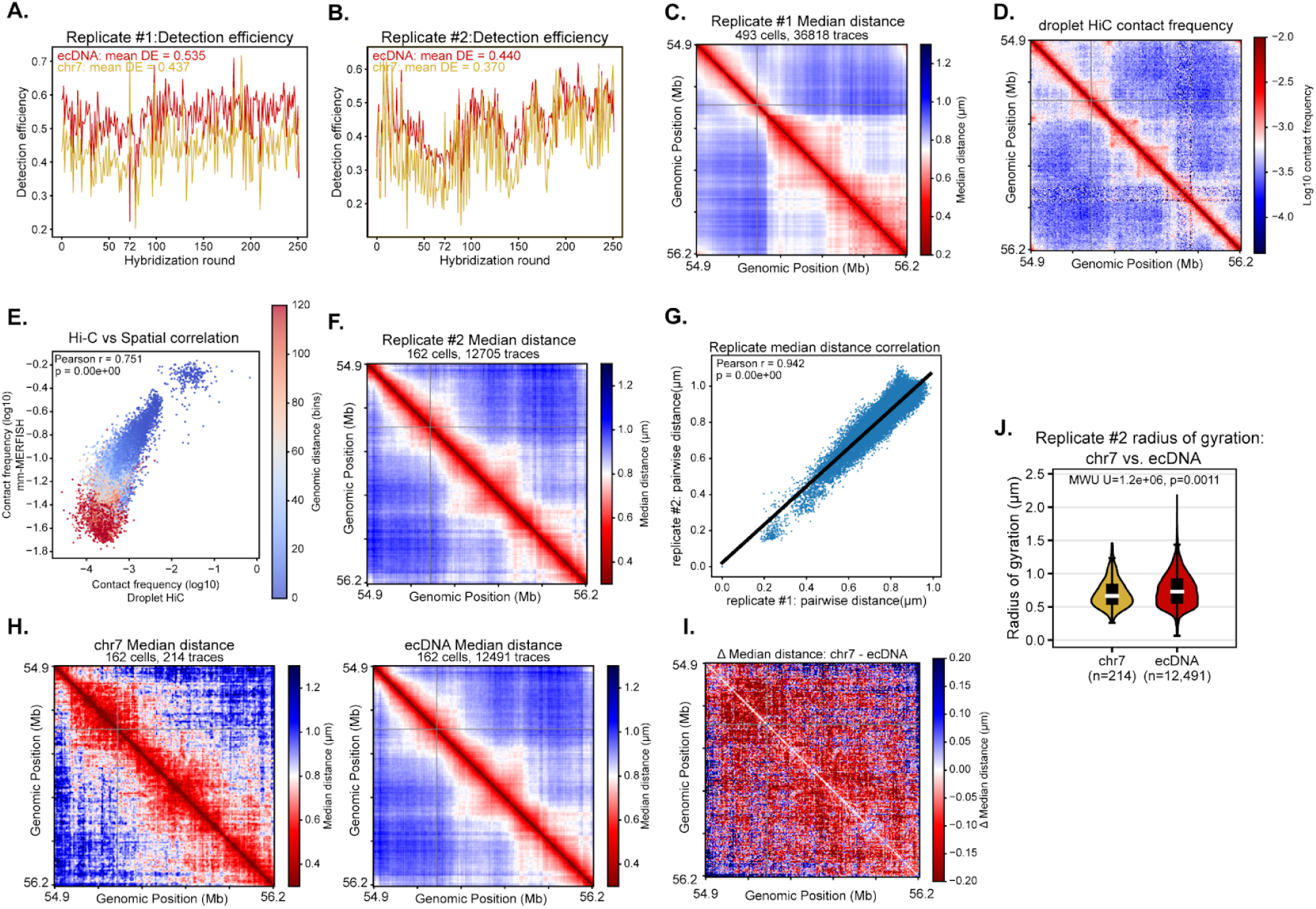
ecDNA is more globally expanded than the matching chromosomal locus across replicates. **(A,B)** Detection efficiency across 5-Kb segments in replicate #1 (A.) and in replicate #2 (B.) for ecDNA (blue) and chr 7 (orange) traces **(C.)** Median distance between all 5-Kb segments across all traces in replicate #1. **(D.)** Droplet HiC log10 contact frequency on the corresponding EGFR locus in GBM39. **(E.)** Correlation of HiC contact frequency and mm-MERFISH derived contact frequency. Contact is considered if the pairwise distance between loci is smaller than 250 nm. **(F.)** Median distance between all 5-Kb segments across all traces in replicate #2. **(G.)** Correlation of the median pairwise distance of 5-Kb segments between replicate #1 and #2. Black line marks y = x. **(H.)** Median distance matrix for the chr7 EGFR locus (left) and ecDNA (right) in replicate #2. **(I.)** Difference in median physical distance between chr7 EGFR locus and ecDNA in replicate #2. **(J.)** Violin plots of the radius of gyration of chr7 EGFR locus and ecDNA in replicate #2. A two-sided Mann Whitney U test was used.4 to test for statistical significance.

**Extended Data Figure 3.**
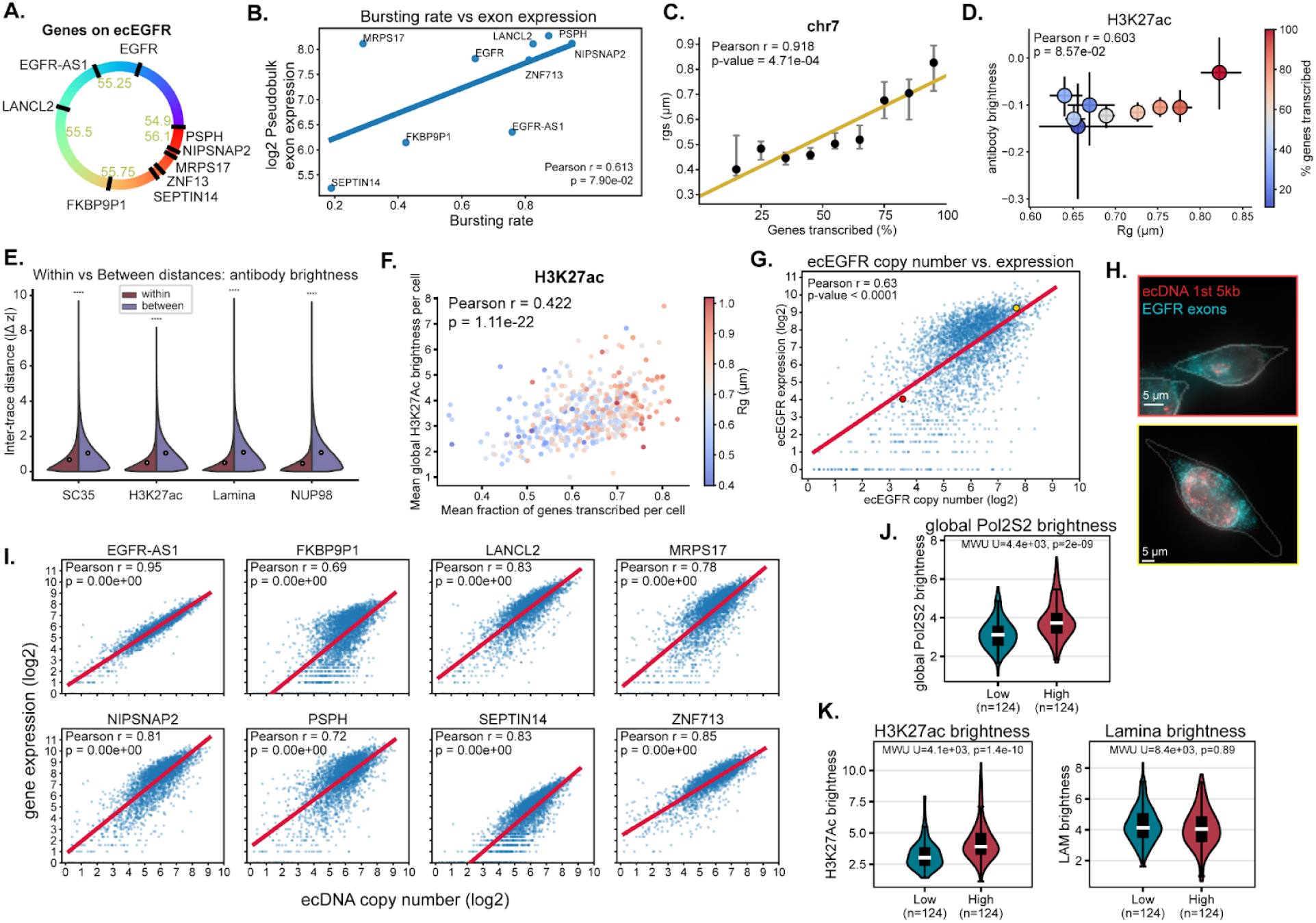
Physical expansion, deposition of regulatory machinery, cellular state, and copy number correlate with ecDNA transcriptional activity. **(A.)** Schematics showing the genes encoded on ecEGFR and their genomic location. **(B.)** Correlation between the average nascent transcription rate vs. average mRNA expression for genes on ecEGFR across 493 cells. mRNA expression is measured by counting exonic foci in the cytoplasm and nucleus (Methods). **(C.)** Stratified scatter plot of the percentage of genes transcribed and the corresponding radius of gyration for the chr7 locus. Black dot represents the median per group, and the bar represents the 95% confidence interval. Orange line marks the fitted OLS line. **(D.)** Correlation between radius of gyration and the normalized H3K27ac brightness across ecDNA traces with different percentages of transcribed genes. Dots mark medians and error bars mark 95% confidence intervals. **(E.)** Split violin plot comparisons of z scored differences across ecDNA from within the same cell vs. from different cells. White circles mark medians. N = 438,277 simulations. White circles mark medians. A two-sided Mann-Whitney U test was used to compute statistical significance. **** - p < 1e-4. **(F.)** Scatter plot comparing the mean transcription and global H3k27ac deposition per cell, colored by the mean Rg per cell. N = 493 cells. **(G.)** Scatter plot of ecDNA copy number (log2) and EGFR expression (log2). Red line marks fitted OLS line. The red circle marks a representative low copy cell and the yellow circle marks a representative high copy cell. **(H.)** Representative images of low ecDNA copy cell (top) and high ecDNA copy cell (bottom) highlighted in G). Red - first 5kb of ecDNA. Cyan - EGFR exon signal. Nucleus represented by DAPI stain (gray). Gray contour - cytoplasm boundaries. **(I.)** Scatter plot of ecDNA copy number (log2) and expression of each gene on ecEGFR (log2). The red line marks the fitted OLS line. **(J.)** Violin plot comparing the global Pol2S2 brightness between the low and high copy cells. A one-sided Mann-Whitney U test was used to test for statistical significance. **(K.)** Violins comparing the H3K27ac and Lamina A/C antibody deposition between the low copy and high copy cells. A one-sided Mann-Whitney U test was used to test for statistical significance.

**Extended Data Figure 4.**
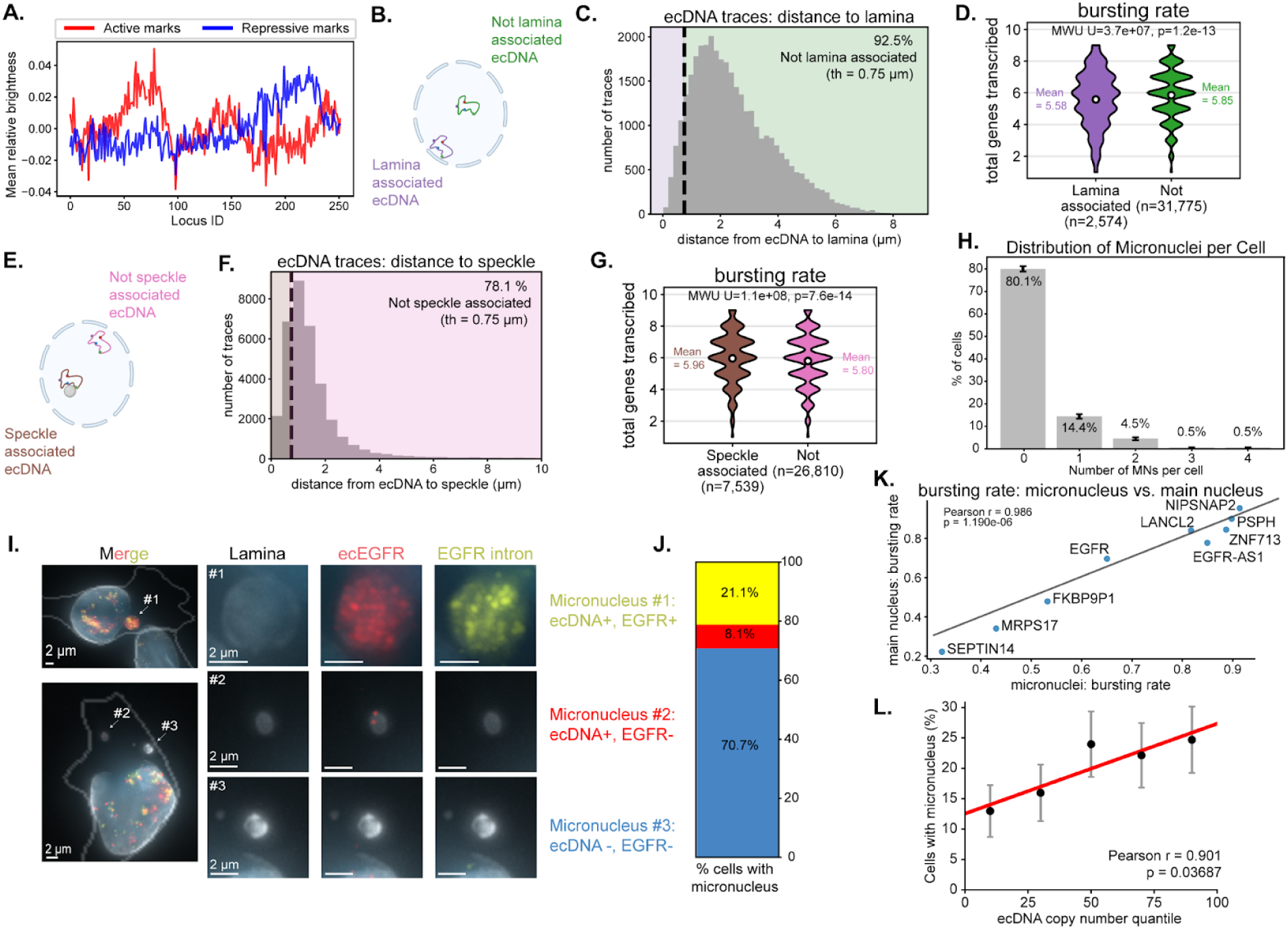
Subcellular localization of ecDNA and its relation to chromatin structure and expression. **(A.)** Mean relative brightness of active and repressive antibody marks across each 5-Kb segment on the ecDNA. Active marks: polII, SC35, H3K27ac. Repressive marks: Lamina, NUP98. **(B.)** Schematics defining lamina associated ecDNA. If the distance between the centroid of an ecDNA molecule to the nuclear lamina is smaller than 750nm (purple), the ecDNA is considered ‘lamina associated ecDNA’. **(C.)** Distribution of distance to lamina across all ecDNA molecules. **(D.)** Violin plots comparing the nascent transcription rate between lamina-associated ecDNA vs. not. White dot marks mean. A one-sided Mann-Whitney U test was used to test for statistical significance. **(E.)** Schematics defining speckle associated ecDNA. If the distance between the centroid of an ecDNA molecule to the nearest spliceosome is smaller than 750nm (brown), the ecDNA is considered ‘speckle associated ecDNA. **(F.)** Distribution of distance to speckle across all ecDNA molecules. **(G.)** Violin plots comparing the nascent transcription between speckle-associated ecDNA vs. not. White dot marks mean. **(H.)** Histogram of the number of micronuclei per cell. Bar represents the standard error of mean. N = 1193 cells. **(I.)** Representative images of cells with micronuclei (white arrow). #1: micronucleus with ecDNA and nascent EGFR transcription. #2: micronucleus with only ecDNA and no nascent EGFR. #3: micronucleus without ecDNA and nascent EGFR. **(J.)** Stacked bar graph showing the status of containing ecDNA and nascent EGFR in cells with micronuclei. N = 238 cells. **(K.)** Scatter plot showing the average bursting rate per gene across all measured ecDNA molecules when residing inside the main nucleus vs inside the micronucleus. Gray line marks y = x. **(L.)** Fraction of cells with micronuclei within different ecDNA copy number bins. Black dot represents the median per group, and the bar represents the 95% confidence interval. The red line represents the fitted OLS line.

**Extended Data Figure 5.**
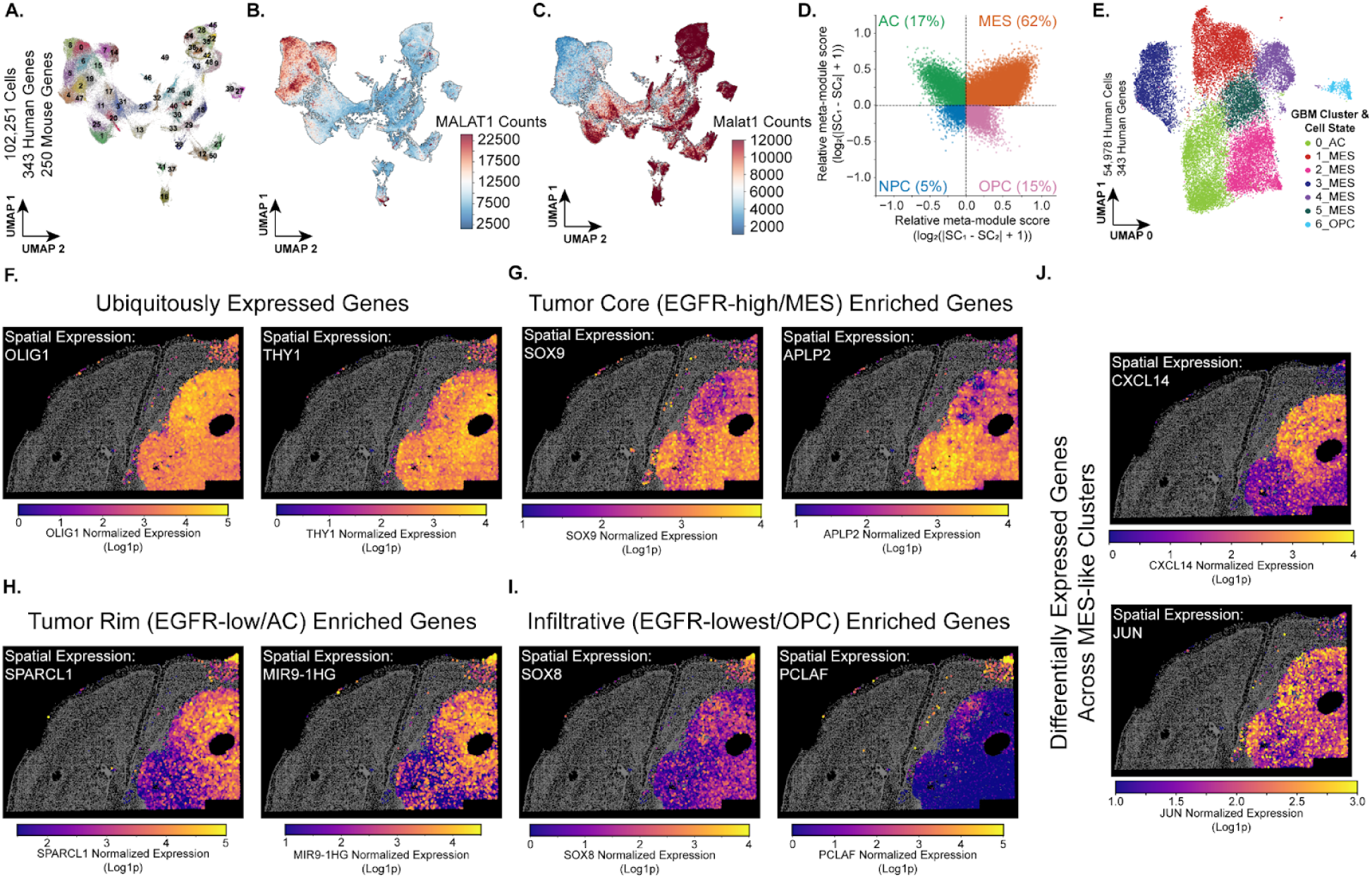
The transcriptional heterogeneity and spatial organization of GBM cells in a chimeric orthotopic xenograft model system. **(A.)** UMAP of both human and mouse cells from RNA-MERFISH data across 343 human genes and 250 mouse genes. Leiden clusters are annotated. **(B.-C.)** Human *MALAT1* (B.) and murine *Malat1* (C.) expression marked on the UMAP in A. **(D.)** Four-quadrant plot illustrating the distribution of GBM cellular states in the GBM39 orthotopic xenograft. Each cell was assigned a state score based on their enrichment of genes in each GBM cellular state gene set. Each quadrant in the plot represents the highest cellular state score for that particular cell and the percentage of cells of that particular state are labeled in each quadrant. **(E.)** UMAP of GBM39 cells annotated by GBM cluster (consistent with Fig.3C and Extended Data Fig.5D). **(F.-J.)** Image of the single-cell RNA expression measured by RNA-MERFISH across all human cells for different marker genes of the GBM subpopulations: (F.) *OLIG1* and *THY1* (ubiquitously expressed genes), (G.) *SOX9* and *APLP2* (tumor core and MES- enriched genes), (H.) *SPARCL1* and *MIR9-1HG* (tumor rim and AC- enriched genes), (I.) *SOX8* and *PCLAF* (infiltrative and OPC- enriched genes), and (J.) *CXCL14* and *JUN* (differentially expressed genes across MES-like clusters).

**Extended Data Figure 6.**
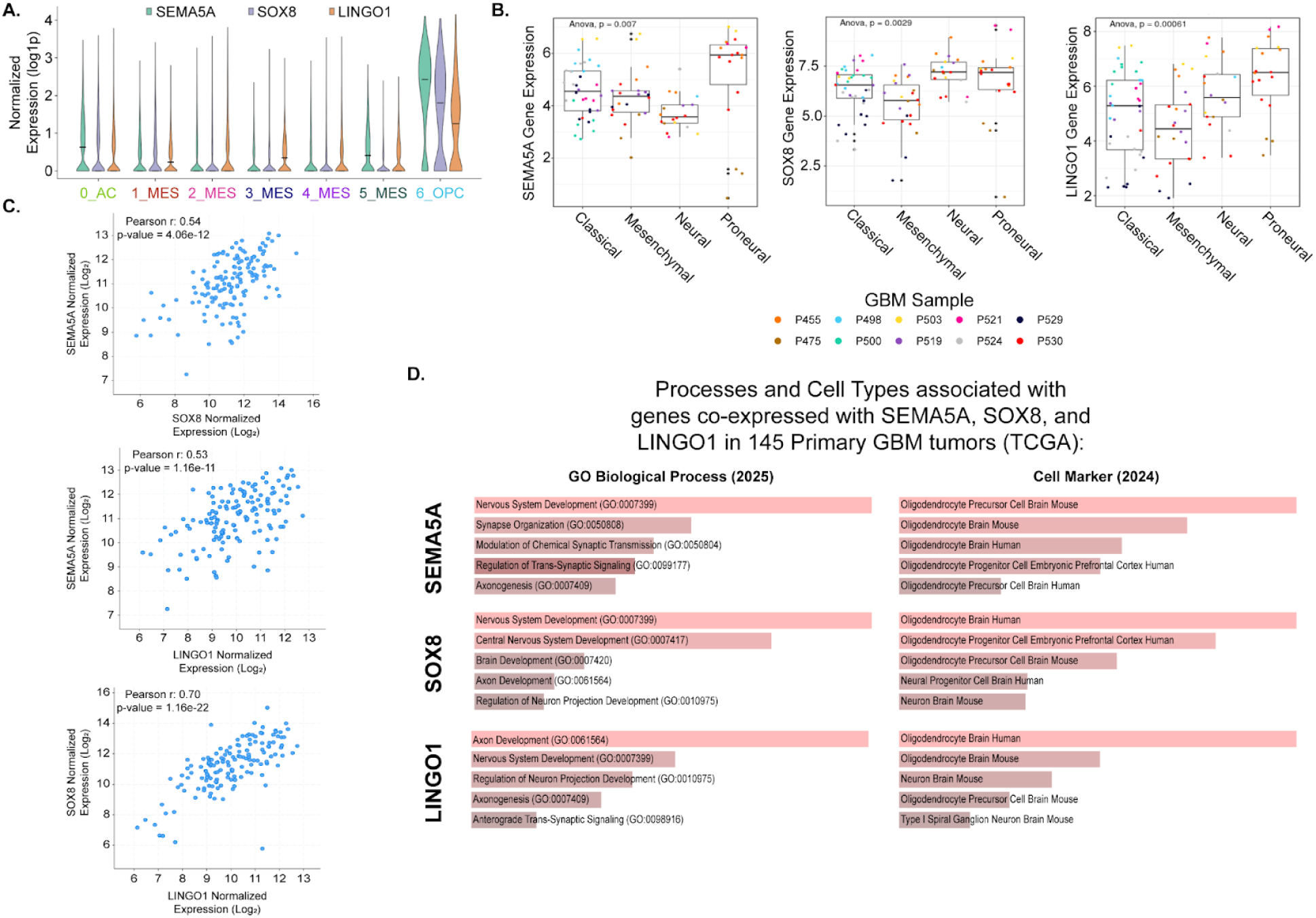
6_OPC marker genes are co-expressed, enriched in proneural GBM, and linked to OPC, neuronal, and synaptic developmental programs. **(A.)** Grouped violin plots of the normalized gene expression (log1p) of *SEMA5A*, *SOX8*, and *LINGO1* across GBM clusters. **(B.)** Normalized gene expression of *SEMA5A*, *SOX8*, and *LINGO1* from bulk RNA sequencing data of 10 primary GBM tumors that were multi-regionally sampled and classified into bulk GBM subtypes: Classical, Mesenchymal, Neural, and Proneural. Each color represents a different GBM tumor, and every datapoint represents a single region that was sampled (i.e., one tumor can have multiple samples that were sequenced). **(C.)** Scatterplots showing the correlation between normalized gene expression (Log2) of *SEMA5A*, *SOX8*, and *LINGO1* across bulk RNA sequencing data from 145 primary GBM samples from TCGA. **(D.)** Gene enrichment bar plots for Gene Ontology Biological Processes (left) and Cell Marker (right) genesets of the 100 genes that most strongly correlate positively with SEMA5A (top), SOX8 (middle), and LINGO1 (bottom) from 145 primary GBM samples.

**Extended Data Figure 7.**
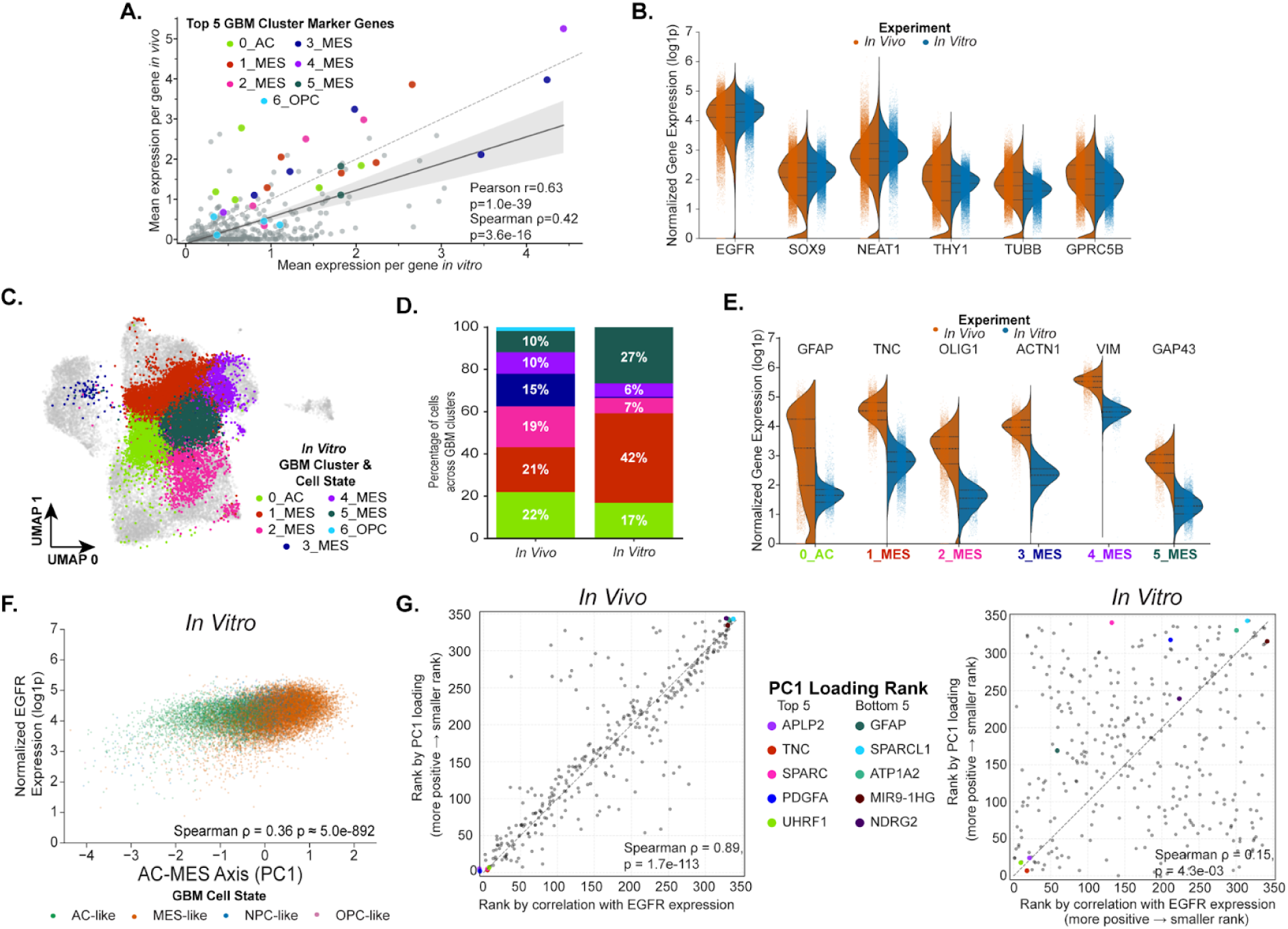
GBM cells adopt more heterogeneous transcriptional states *in vivo*. **(A.)** Correlation of average gene expression between GBM39 *in vitro* (X-axis) and *in vivo* (Y-axis). Dotted grey line marks x=y. The dark grey line shows the Pearson linear regression with confidence intervals (light grey bands) with corresponding correlation coefficient and p-value. The top five marker genes for each GBM cluster are colored. **(B.)** Split violin plots of normalized expression (log1p) across representative genes with similar expression *in vivo* (left; orange) and *in vitro* (right; blue) experiments. **(C.)** UMAP of GBM39 *in vivo* (grey) onto which in vitro GBM39 cells (colored) were integrated. Colors mark different GBM clusters. **(D.)** Stacked bar plots illustrating the proportional differences of GBM clusters across *in vitro* (right) and *in vivo* (left) experiments. **(E.)** Split violin plots with the expression of representative marker genes in matching clusters between *in vivo* (left; orange) and *in vitro* (right; blue) experiments. **(F.)** Correlation of Principal Component 1 (PC1 - from single-cell expression data) and *EGFR* expression across single-cells for GBM39 *in vitro*. Cells were colored by GBM cellular state. **(G.)** Correlation between each gene’s ranked PC1 loading score and their ranked correlation with *EGFR* expression. For both experiments, each gene’s expression was correlated with *EGFR* expression across all cells. All genes were then ranked based on their Pearson correlation by which strongly positively correlative genes were ranked lowly (e.g., 1, 2, … *N*) and strongly negatively correlated genes were ranked highly (e.g., *N*, *N* – 1, etc.). All genes’ PC1 loading scores were also ranked such that strongly positive PC1 loading scores were ranked lowly (e.g., 1, 2, … *N*) and strongly negative PC1 loading scores were ranked highly (e.g., *N*, *N* – 1, etc.). The top five genes for the positive (“Top 5”) and negative (“Bottom 5”) PC1 loading scores for GBM *in vivo* were annotated on both plots.

**Extended Data Figure 8.**
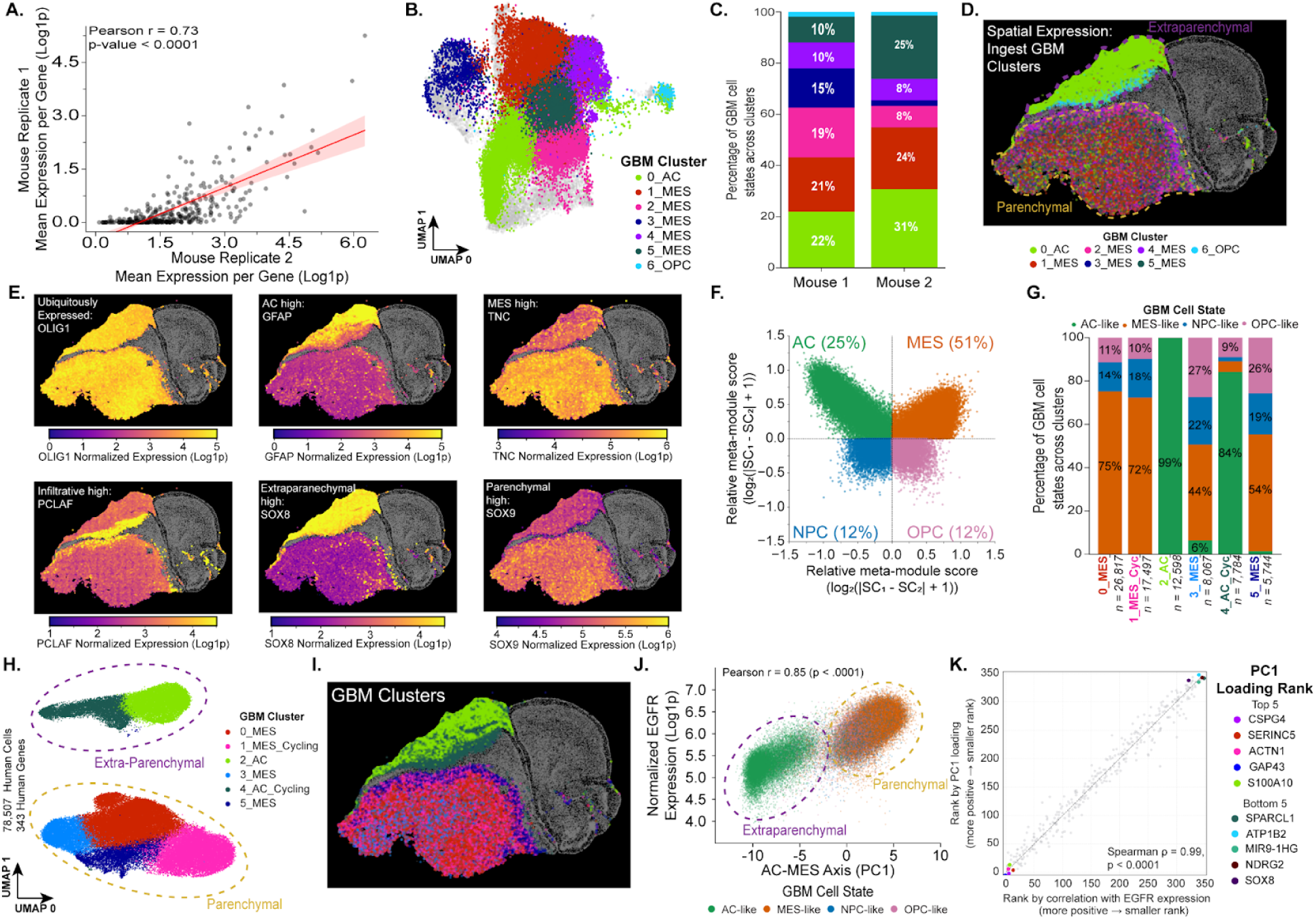
Independent mouse recapitulates general features of GBM spatial organization. (**A.)** Correlation of average gene expression of human genes across mouse replicates. The red line shows the Pearson linear regression with confidence intervals (light red bands), correlation coefficient, and p-value indicated. **(B.)** UMAP of GBM39 mouse replicate 1 (grey) onto which mouse replicate 2 cells (colored) were integrated. Colors mark different GBM clusters. **(C.)** Stacked bar plots illustrating the proportional differences of GBM clusters across mouse replicate 1 (left) and mouse replicate 2 (right) experiments. **(D.)** Spatial distribution for mouse replicate 2 of GBM clusters transferred from mouse replicate 1 (see panel B-C). All mouse cells are annotated in grey. Parenchymal (top) and extraparenchymal (bottom) tumors were annotated with dotted lines **(E.)** Image of single cell expression of representative genes with variable spatial patterns: *OLIG1* (ubiquitously expressed), *GFAP* (AC-enriched), *TNC* (MES-enriched), *PCLAF* (infiltrative-enriched), *SOX8* (extraparenchymal-enriched), and *SOX9* (parenchymal-enriched). All mouse cells are annotated in grey. **(F.)** Four-quadrant plot illustrating the distribution of GBM cellular states in the GBM39 orthotopic xenograft replicate #2. Each cell was assigned a state score based on their enrichment of genes for each GBM cellular state. Each quadrant in the plot represents the highest cellular state score for that particular cell and the percentage of cells of that particular state are labeled in each quadrant. **(G.)** Stacked bar plot depicting the proportions of different GBM cellular states across Leiden clusters in replicate #2. **(H.)** De-novo UMAP of GBM39 cells in mouse replicate #2. De-novo Leiden clusters annotated by the dominant GBM cellular state are colored. Parenchymal (gold) and extra-parenchymal cells encircled on the UMAP **(I.)** Spatial distribution of GBM clusters defined in (G). Mouse cells are annotated in grey. **(J.)** Correlation of Principal Component 1 (PC1 - from single-cell expression data) and *EGFR* expression across single-cells. Cells were colored by their GBM cellular state (see Extended Data Fig.8F). Parenchymal (gold) and extraparenchymal (purple) cell populations are marked with dashed outlines. **(K.)** As in Extended Data Fig.7G (see methods), correlation between each gene’s ranked PC1 loading score and their ranked correlation with *EGFR* expression.

**Extended Data Figure 9.**
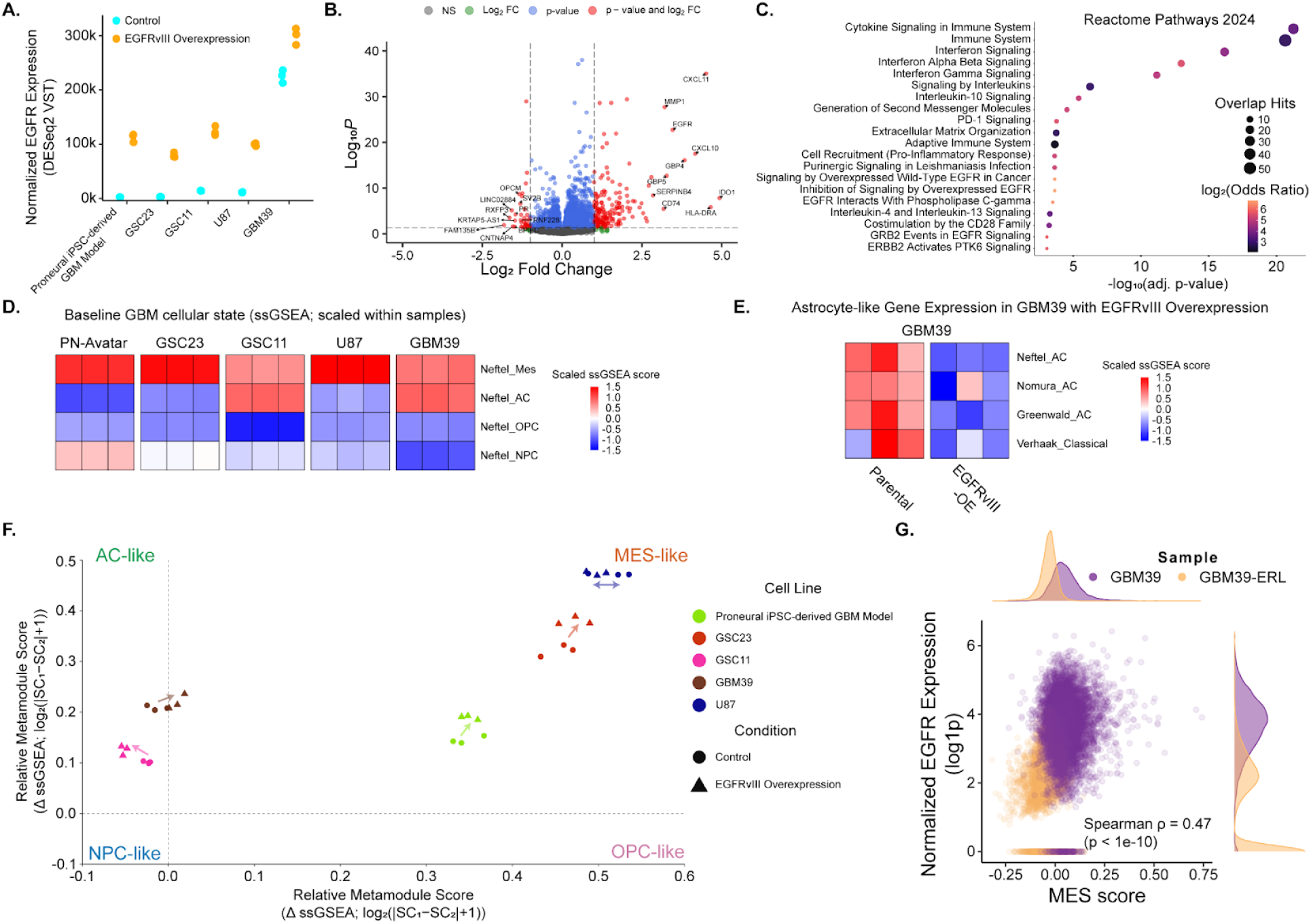
Genetic and pharmacologic perturbations of EGFR expression shift global gene expression and GBM cellular states. **(A.)** *EGFR* expression across five GBM cell lines with (blue) and without (orange) EGFRvIII overexpression. **(B.)** Volcano plot illustrating differentially expressed genes between *EGFRvIII*-overexpression and control conditions. Positive fold changes indicate genes that are more highly expressed in EGFRvIII-overexpression lines, while negative fold changes indicate genes that are more highly expressed at baseline. The 10 genes with highest magnitude in positive and negative fold changes are labeled on the plot. **(C.)** Gene enrichment analysis for the Reactome Pathways 2024 was performed with Enrichr<u>^84^</u> for all genes with a 2-fold increase and a Padj < 0.05 in expression following *EGFRvIII*-overexpression. The top 20 pathways are shown in a dot plot indicating the p-value (X-axis), overlapping hits (size), and the odds ratio (color). **(D.)** Heatmap of ssGSEA scores for Neftel et al. GBM cellular state gene sets<u>^33^</u> across five GBM cell lines at baseline. ssGSEA scores were scaled across cellular states, indicating relative expression of GBM cellular states within each cell line. **(E.)** Heatmap of ssGSEA scores for four<u>^33,37,60,67^</u> different GBM AC-like/Classical gene sets for GBM39 with (*EGFRvIII*-OE) and without *EGFRvIII* overexpression. ssGSEA scores were scaled across replicates (rows) indicating relative AC-like gene expression following EGFRvIII overexpression. **(F.)** Four-quadrant plot illustrating the distribution of GBM cellular states in five GBM cell lines with and without EGFRvIII overexpression. ssGSEA scores were computed for four GBM cellular states,<u>^33^</u> and each quadrant in the plot represents the highest cellular state ssGSEA score for that particular cell line. **(G.)** Single-cell correlation between *EGFR* expression and MES-like score for GBM39 (purple) and GBM39-ERL (orange). Annotated histograms illustrate the density of cells at particular *EGFR* expression levels and MES-like scores.

**Extended Data Figure 10.**
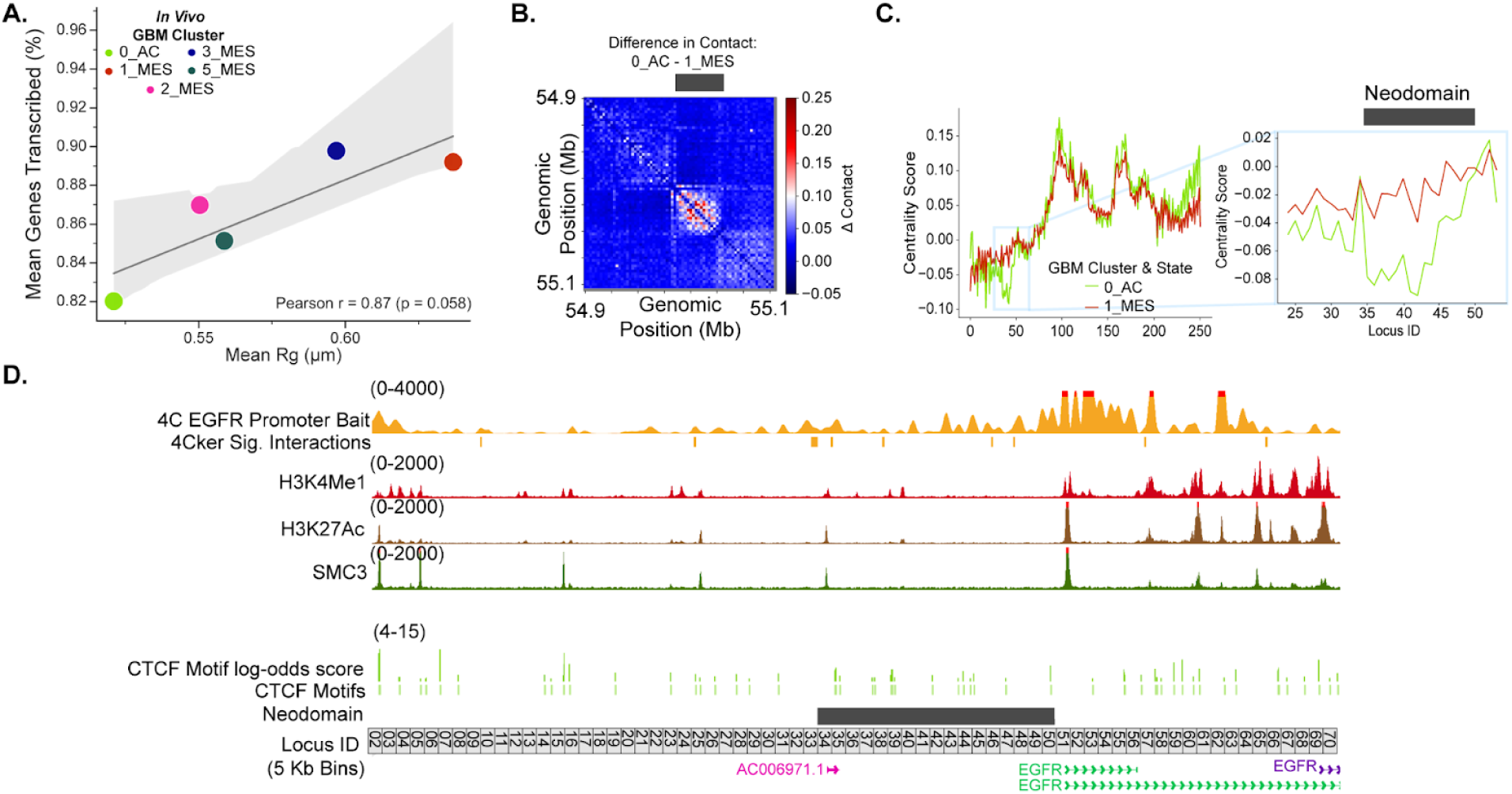
3D chromatin organization and activity of ecEGFR across GBM states. **(A.)** Correlation between the mean Rg and transcriptional activity across GBM clusters *in vivo*. Clusters 4_MES, containing a distinct ecEGFR structural variant, and 6_OPC, containing chromosomal re-integrated ecDNA, were omitted from this analysis. The Pearson linear regression (dark grey line) with confidence intervals (light grey bands), correlation coefficient, and p-value are indicated within the panel. **(B.)** Pairwise log contact differences between 0_AC and 1_MES GBM clusters. Contacts were defined as loci separated by <250 nm, and pairwise contacts were computed for each ecEGFR trace and averaged within clusters. White and red represent higher contact frequency in 0_AC, whereas dark blue represents higher contact frequency in 1_MES. The dark grey block at the top of the matrix marks the neo-domain present in 0_AC cells but not in 1_MES cells. **(C.)** Centrality scores for each 5-Kb locus on ecDNA. Centrality was computed (Methods) from the median distance matrices for 0_AC (green) and 1_MES (red) clusters. The light blue inset highlights the difference between the two clusters at the 0_AC-specific neo-domain region (bin ∼34-50). **(D.)** Genome track visualization of the following assays of GBM39 *in vitro*: 4C with the EGFR promoter as the anchor (bait) region (1st track), significant 4C interactions between the EGFR promoter (2nd track), active histone mark H3K4me1 ChIP-seq (third track), active histone mark H3K27ac ChIP-seq (fourth track), the cohesin component SMC3 ChIP-seq (fifth track), the log-odds score for candidate CTCF motifs (sixth track), and statistically significant CTCF motifs (seventh track). 0_AC-specific neo-domain identified in chromatin tracing data (Fig.4F-G) and each 5 Kb bin are the final two tracks, respectively.

**Extended Data Figure 11.**
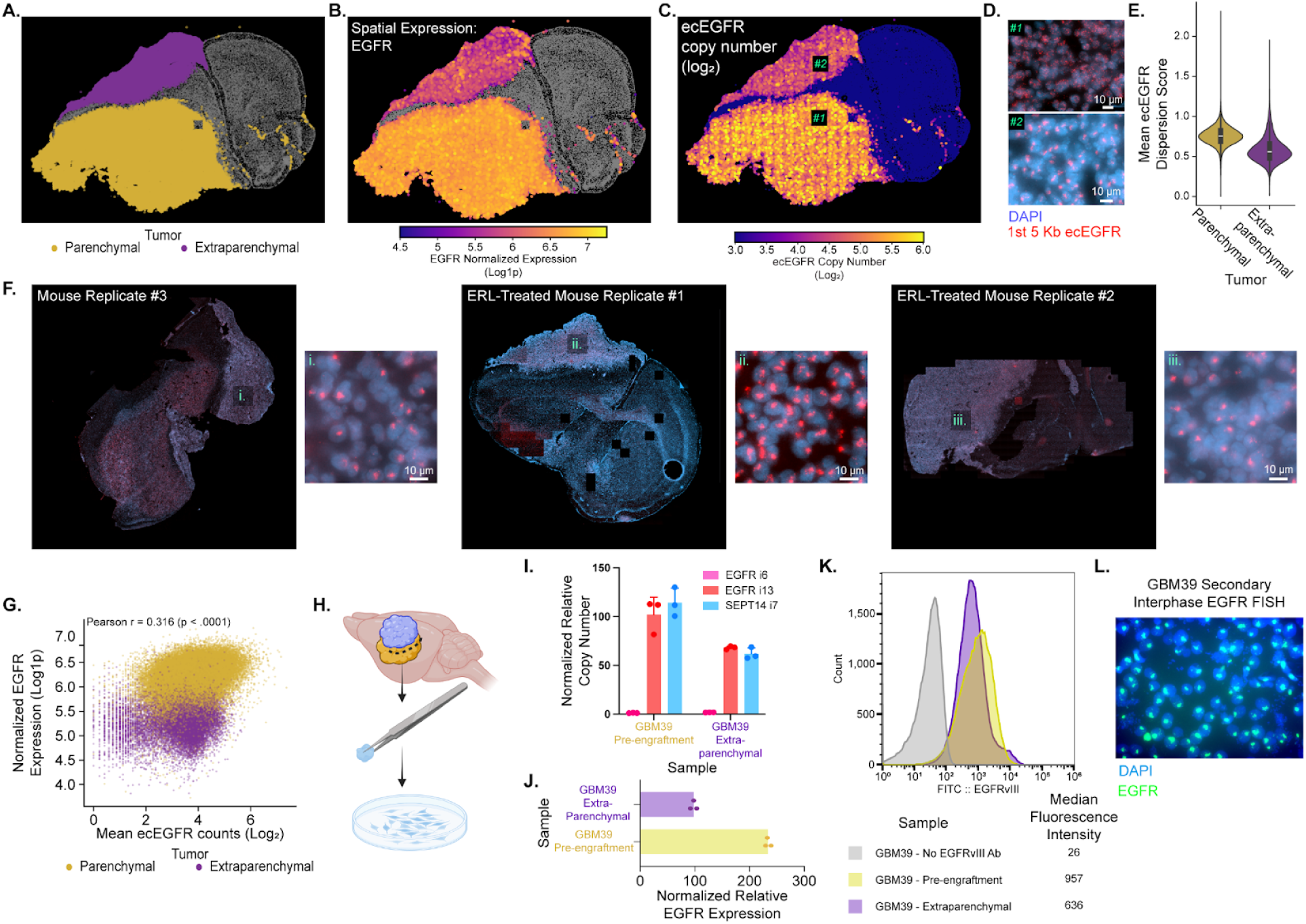
A multifocal GBM model reveals diverse ecEGFR phenotypes associated with distinct spatial contexts. **(A.)** Image of parenchymal (purple) and extraparenchymal (gold) tumors. Mouse cells are annotated in grey. **(B.-C.)** Images of EGFR expression (B.) and ecEGFR counts (C.). All mouse cells are annotated in grey. **(D.)** Representative images of DNA MERFISH of the first 5-Kb region of ecEGFR (red) across parenchymal (1) and extraparenchymal (2) regions. DAPI - blue. **(E.)** Violin plot of ecEGFR Dispersion Scores across parenchymal and extraparenchymal cells. **(F.)** Representative DNA-MERFISH images of the first 5-Kb region of ecEGFR (red) within parenchymal tumors across an additional untreated mouse replicate (left) and two mouse models treated with the EGFR inhibitor erlotinib (center, right). DAPI - blue. **(G.)** Single-cell correlation between ecEGFR counts and normalized EGFR expression annotated by their location. **(H.)** Schematic illustrating how secondary tumor cells were dissected from orthotopic xenografts and cultured *in vitro*. **(I.)** DNA qPCR was done for ecEGFR regions (EGFRi13, red, and SEPT14i7, blue) and a deleted region of ecEGFR (EGFRi6, pink) for GBM39 cells pre-engraftment and extraparenchymally-derived GBM39 cells. ΔΔCt analysis was performed, and values were normalized within samples to DNA from regions on chromosome 7. Values were then normalized between samples to the diploid cell line iNPC12. **(J.)** RT-qPCR was performed to compare EGFR expression levels between GBM39 cells pre-engraftment (yellow) and extraparenchymally-derived GBM39 cells (purple). ΔΔCt analysis was performed by normalizing to GAPDH expression within samples and normalizing values to iNPC12. **(K.)** FACS histogram showing the distributions of EGFR expression across GBM39 cells pre-engraftment (yellow) and extraparenchymally-derived GBM39 cells (purple). GBM39 cells without EGFRvIII antibody (grey) were used as a negative control, and median fluorescence intensity values are indicated below for each population. **(L.)** EGFR FISH was performed on extraparenchymally-derived GBM39 cells in interphase. EGFR - green and DAPI - blue.

**Extended Data Figure 12.**
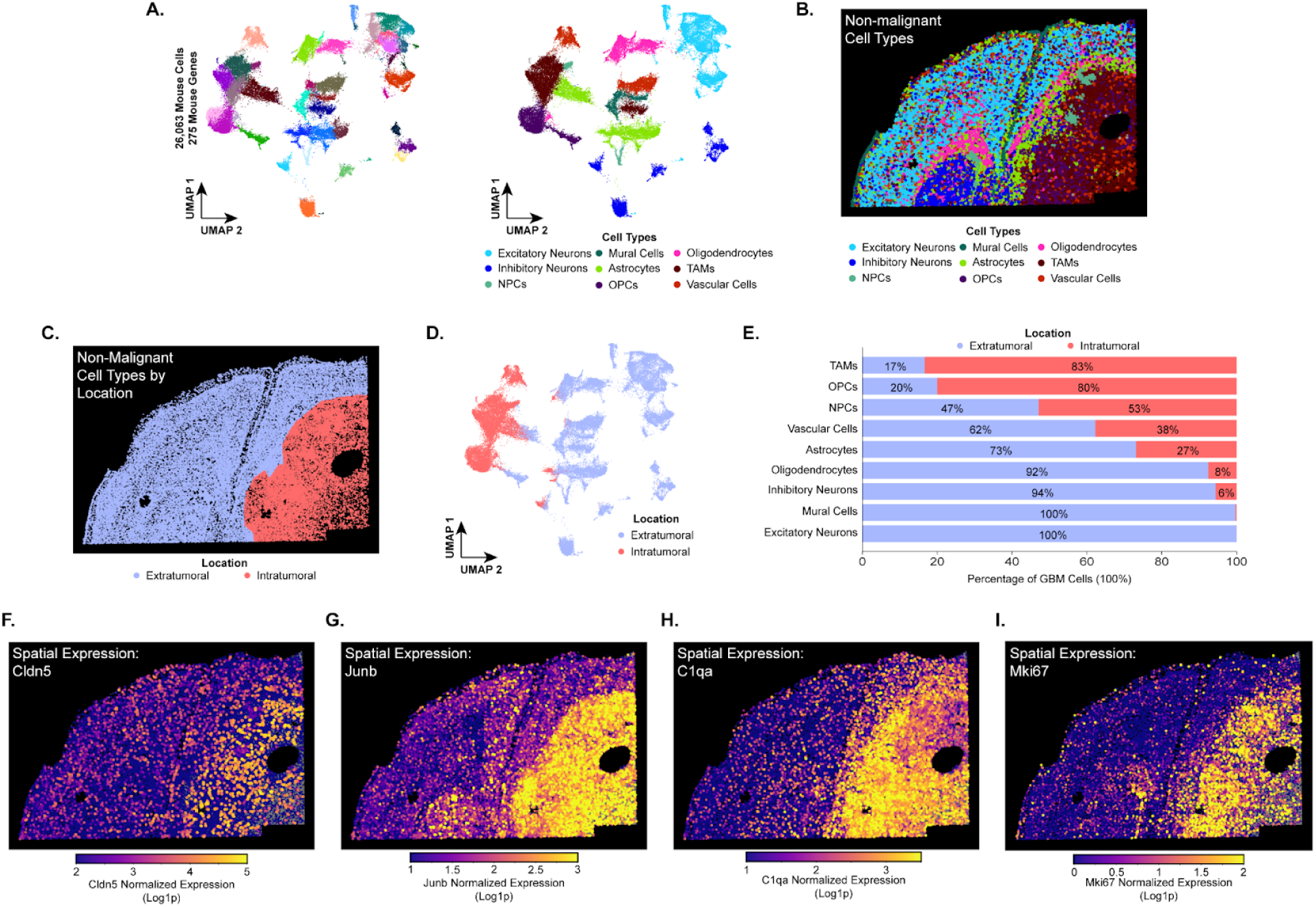
Defining the spatial organization of the non-malignant microenvironment in a xenograft model of GBM. **(A.)** UMAP of non-malignant cells annotated by Leiden cluster (left) and broad cell type (right) defined by the expression of 250 mouse genes. **(B.)** Spatial distribution of non-malignant cell types. **(C.)** Spatial distribution of non-malignant cell types residing within (intratumoral, salmon) or beyond (extratumoral, periwinkle) the tumor. **(D.)** UMAP of non-malignant cells annotated by their spatial relationship with the tumor. **(E.)** Stacked bar plots illustrating the proportions of non-malignant cell types within or beyond the tumor. **(F.-I.)** Spatial distribution of single-cell expression for genes enriched in intratumoral non-malignant cells: *Cldn5* (F.), *Junb* (G.), *C1qa* (H.), and *Mki67* (I.).

**Extended Data Figure 13.**
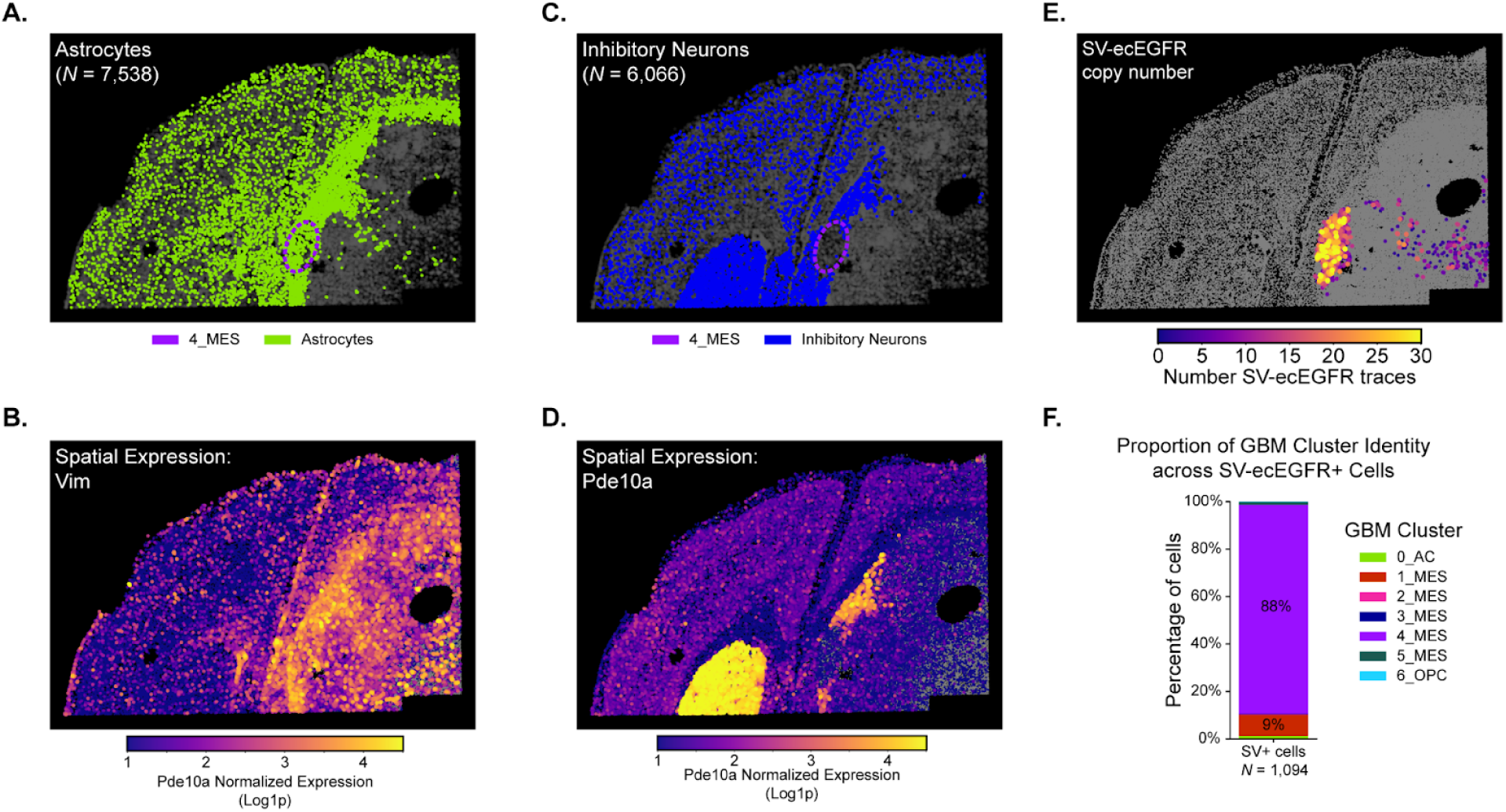
Selection of an ecEGFR structural variant in a unique microenvironment. **(A.)** Spatial distribution of astrocytes (green). Other cell types are plotted in grey. The 4_MES cluster is outlined with a dashed purple line. **(B.)** Single-cell gene expression of the astrocyte marker gene *Vim*. **(C.)** Spatial distribution of inhibitory neurons (blue). Other cell types are plotted in grey. The 4_MES cluster is outlined with a dashed purple line. **(D.)** Single-cell gene expression of the striatal medium spiny neuron marker gene *Pde10a*. **(E.)** Spatial distribution of SV-ecEGFR+ copy number. SV-ecEGFR was determined by loss of detection efficiency in the deleted region (chr7:55.000-55.977-Mb). Cells without SV-ecEGFR are annotated in grey. **(F.)** Stacked bar plot showing the GBM cluster proportions across SV-ecEGFR+ cells.

**Extended Data Figure 14.**
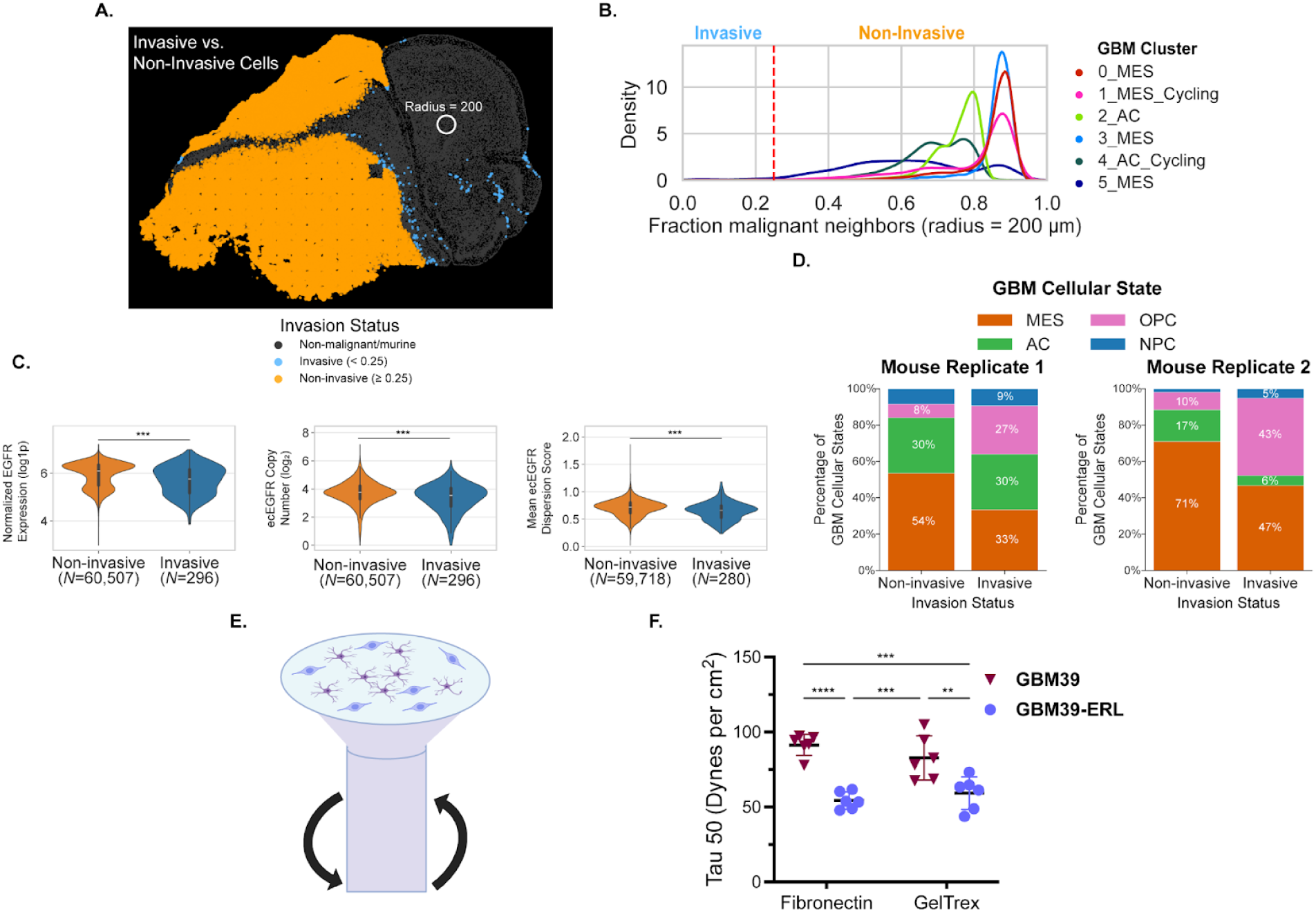
Invasion and cancer cell-ECM adhesion are associated with NPC-and OPC-like states and low EGFR expression. **(A.)** Spatial distribution of invasive (blue) and non-invasive (orange) malignant cells. Non-malignant cells - grey. The fraction of malignant cells within a 200 µm radius (white circle) of each GBM cell was used to define invasive (< 0.25 malignant fraction) and non-invasive (>0.25 malignant fraction) cells.**(B.)** Density plot showing the distribution of malignant neighbors across each GBM cluster. A dashed red line at .25 represents the threshold at which invasive status was classified. **(C.)** Violin plots showing the normalized EGFR exons (left), ecEGFR copy number (middle), and ecEGFR Dispersion Score (right) across non-invasive (orange) and invasive (blue) cells. A Welch t-test with FDR multiple hypothesis testing correction was used to test for statistical significance. *** = p-value < 0.0001. **(D.)** Stacked bar plots illustrating the proportional differences of GBM states across non-invasive and invasive cells for mouse replicate 1 (left) and mouse replicate 2 (right) experiments. **(E.)** Schematic of spinning disc assay that subjects cells to a radially increasing shear stress to assess cell adhesion. **(F.)** Relative Tau, which measures the force required to displace a given percentage of the cells (e.g., 50%) for GBM39 (maroon) and GBM39-ERL (purple), while on two separate extracellular matrices: fibronectin (left) and GelTrex (right). * = p-value < 0.05, ** = p-value < 0.01, *** = p-value < 0.001, **** = p-value < 0.0001.

**Extended Data Figure 15.**
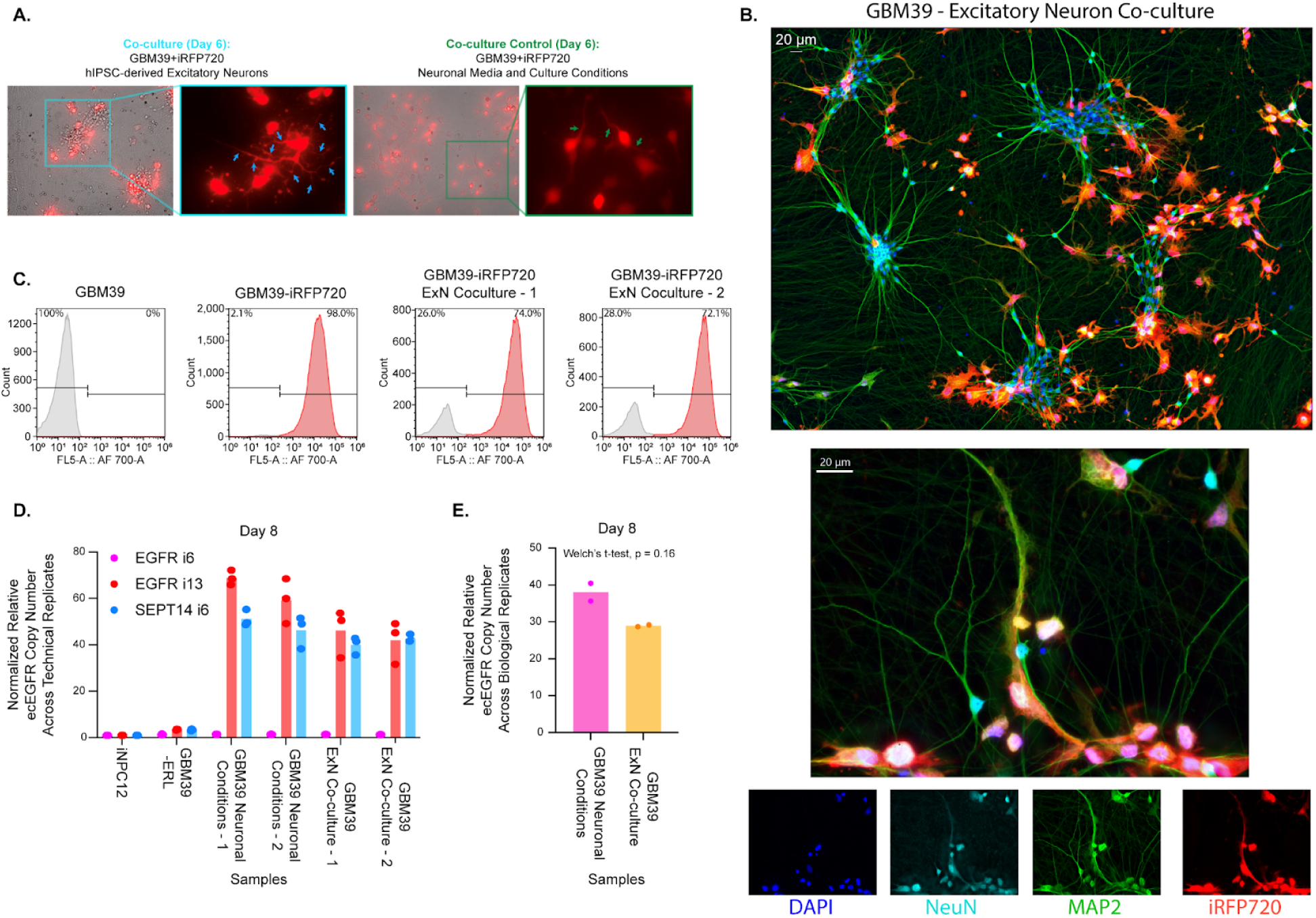
Interactions with excitatory neurons alter GBM cell morphology and decrease ecEGFR copy number. **(A.)** Fluorescence microscope images (20X objective) of GBM39+iRFP720 co-cultured with hIPSC-derived excitatory neurons (left) and GBM39+iRFP720 cells independently grown with the same conditions as the co-culture (right). Cells were imaged six days after co-culturing. Images were merged from brightfield and red channels. The right insets show magnified GBM39+iRFP720 cells solely in the red channel, highlighting differences in cell morphology. Arrows mark GBM projections. **(B.)** Immunofluorescence imaging of GBM39+iRFP720 co-cultured with hIPSC-derived excitatory neurons. DAPI is shown in blue; neuronal markers *NeuN* and *MAP2* are represented in green and magenta, respectively; and iRFP720 signal is shown in red. **(C.)** Flow cytometry-derived histogram showing the distributions of iRFP720 signal across two GBM39+iRFP720 and hIPSC-derived excitatory neuron co-culture experiments (last two panels). GBM39 cells without iRFP720 were used to establish positive gates (1st panel). GBM39+iRFP720 mono-culture (2nd panel) was used as a positive control. **(D.)** DNA qPCR was performed for ecEGFR regions (EGFRi13, red, and SEPT14i7, blue) and a DNA region not present on ecEGFR (EGFRi6, pink) for GBM39+iRFP720 and hIPSC-derived excitatory neuron co-culture experiments and GBM39 cells grown in identical culture conditions sans excitatory neurons. ΔΔCt analysis was performed, and values were normalized within samples to DNA from regions on the endogenous chromosome 7. Values were then normalized between samples to the diploid cell line iNPC12. **(E.)** Data from D. was re-analyzed to determine copy number differences between co-cultures and control cells. Mean values from technical qPCR replicates were taken for each measured locus, and the mean values from the two ecEGFR regions (EGFRi13 and SEPT14i6) were calculated and plotted for each biological replicate. Welch’s t-test was used to determine statistical significance.

**Extended Data Figure 16.**
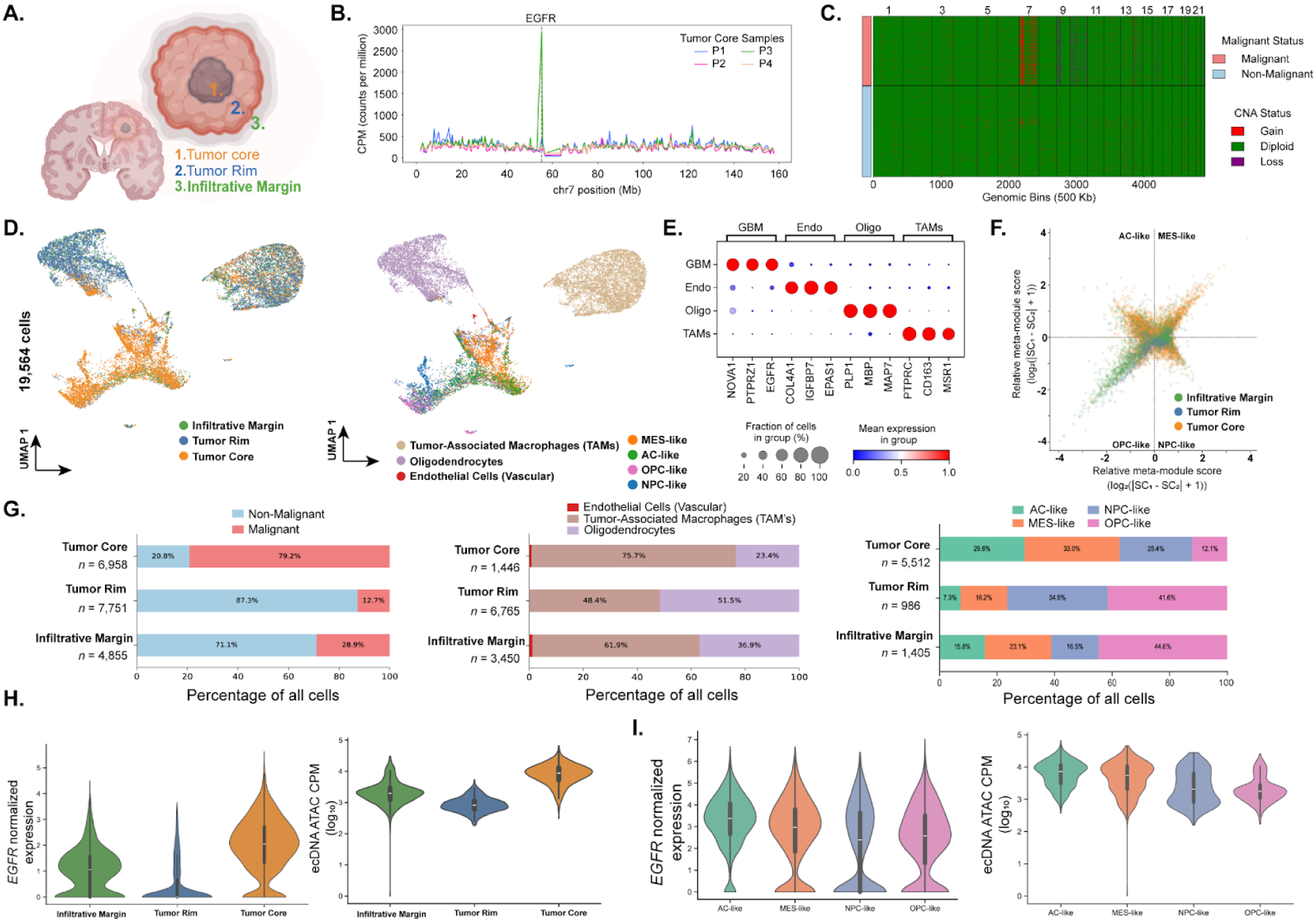
10x-Multiome characterization of ecEGFR and transcriptional states in primary GBM samples. **(A.)** Schematic of experimental design for multi-region ecDNA and cellular states characterization. **(B.)** ATAC fragments plotted across 1-Mb bins from chromosome 7 for each core primary GBM sample. **(C.)** Heatmap showing copy number gains (red), deletions (purple), or diploid (green) status computed by the EpiAneufinder^65^ algorithm. **(D.)** UMAP of primary patient tumor samples annotated by tumor region (left) and broad cell type (right) **(E.)** Dot plot illustrating scaled expression of cell type and cancer marker genes across Leiden clusters **(F.)** Four-quadrant plot illustrating the distribution of GBM cellular states annotated by GBM sample/region. **(G.)** Stacked bar plots showing the proportions of different layers of cell identity across GBM regions: non-malignant vs. malignant (left), normal cell types (middle), and GBM cellular states (right). **(H.)** The distribution of EGFR expression (left) and EGFR ATAC fragments coming from the ecEGFR locus (right) across GBM regions. The ecEGFR locus was inferred from Droplet-HiC.^19^ Log_10_ CPM of ATAC fragments was calculated after per-cell library-size normalization and GC-bias correction. **(I.)** As calculated in E., the distribution of normalized EGFR expression (left) and EGFR ATAC fragments (right) across GBM cellular states.

**Extended Data Figure 17:**
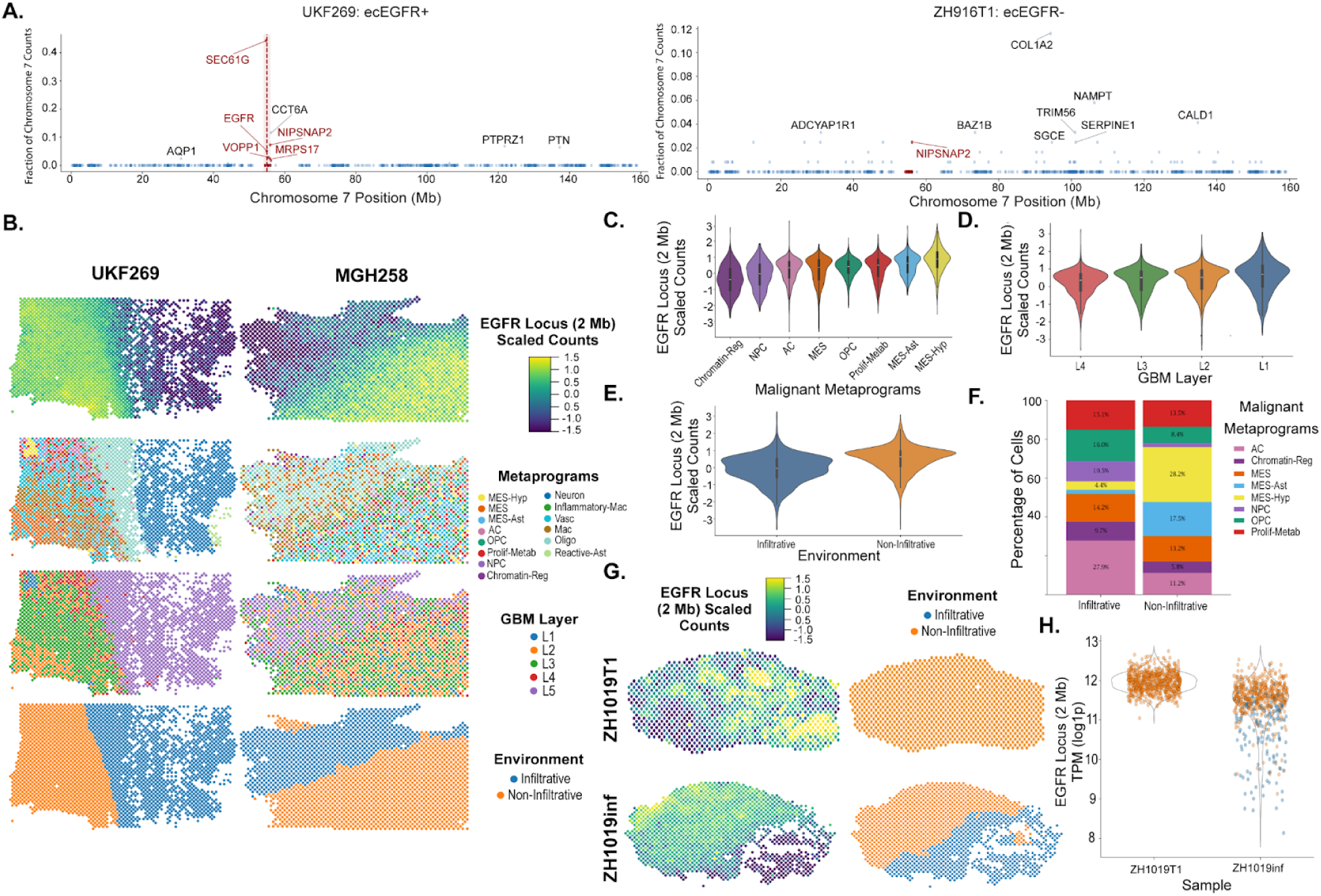
Transcripts from the EGFR locus are spatially organized across the layered architecture of GBM in a broader cohort of patient tumors. **(A.)** Representative Manhattan plots displaying the fraction of gene counts over the total chr 7 counts (Y-axis) against the gene’s position on chr 7 for a sample classified as ecEGFR+ (left) and ecEGFR-(right). The red dotted line and window demarcate 1-Mb up- and downstream of *EGFR* locus (2-Mb total) in the ecEGFR+ sample (left). All genes outside of the window are colored blue, while *EGFR* locus genes are colored red. Y-axes were scaled to samples. **(B.)** Two ecDNA+ samples plotted spatially with each spot annotated by the scaled counts of all genes within the 2-Mb EGFR locus (1^st^ row), their metaprogram (2^nd^ row), their GBM spatial layer (3^rd^ row), and their infiltrative environment status (4^th^ row). **(C.)** Violin plot showing the scaled counts for all genes in the 2-Mb EGFR locus by GBM metaprogram. Only malignant metaprogram spots were analyzed, and groups were ordered by rank. **(D.)** Violin plot showing the scaled counts for all genes in the 2-Mb EGFR locus by GBM layer. Only malignant metaprogram spots were analyzed, and groups were ordered by rank. **(E.)** Violin plot showing the scaled counts for all genes in the 2-Mb EGFR locus by GBM infiltrative environment status. Spots with >30% of neighbors that are neurons, oligodendrocytes, or astrocytes were labeled infiltrative. Only malignant metaprogram spots were analyzed, and groups were ordered by rank. **(F.)** Stacked bar plot of the proportion of GBM metaprograms across infiltrative environment status. **(G.)** Spatial plots for two samples that derive from the same primary GBM patient tumor: one from a T1-contrast enhancing region (top) and one from an infiltrating region (bottom). The plots were annotated by the scaled counts of all genes within the 2-Mb EGFR locus (1^st^ column) and their infiltrative environment status (2^nd^ column). **(H.)** A violin plot of the TPM (log1p) for all counts in the 2-Mb EGFR locus by the samples from (G.) annotated by their infiltrative environment status.

**Extended Data Figure 18.**
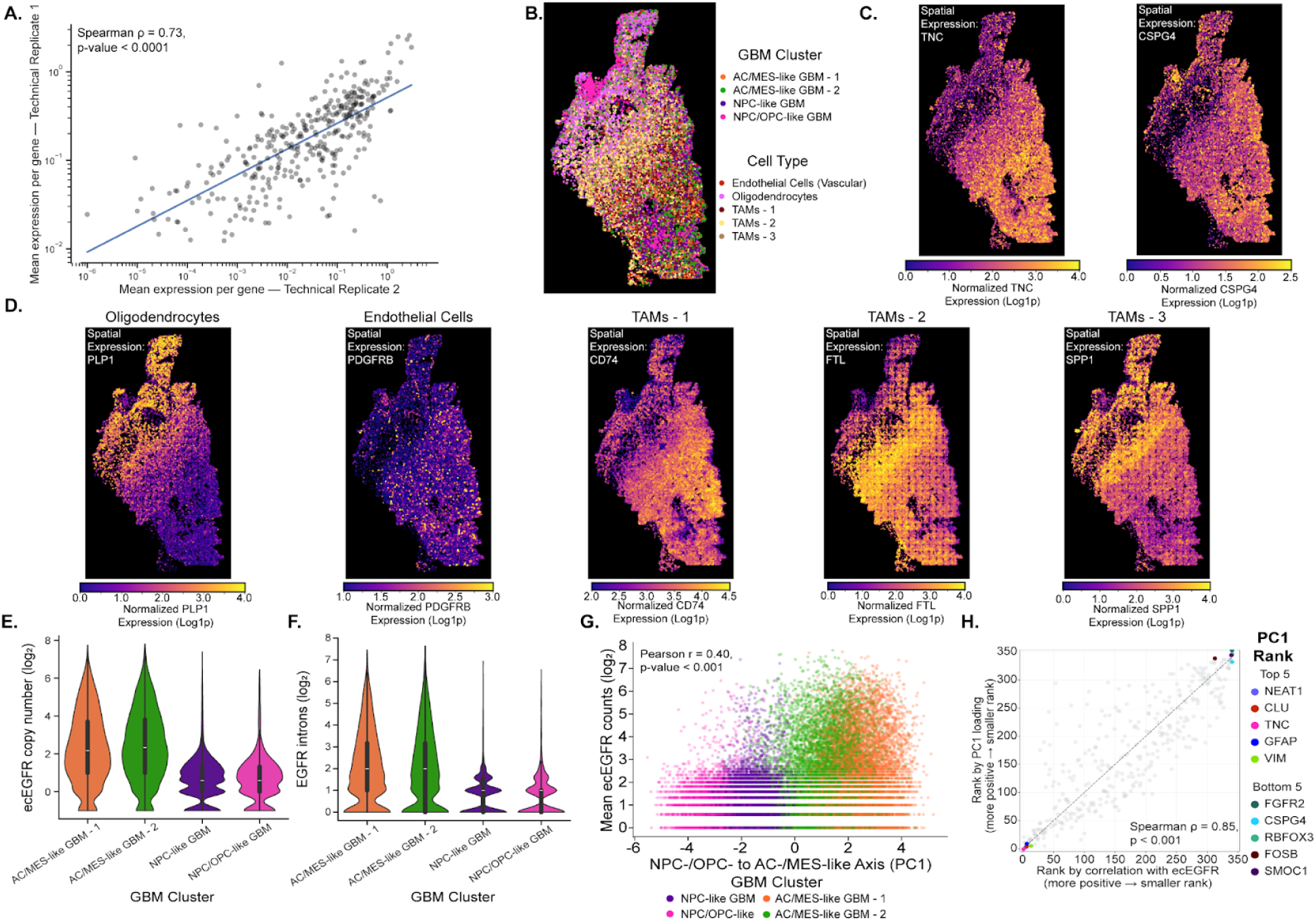
ecEGFR copy number varies across the layered organization of GBM cellular states. **(A.)** Scatter plot showing the log-log gene expression values for primary GBM replicate 2 (X-axis) and primary GBM replicate 1 (Y-axis). The blue line shows the linear regression with confidence intervals (light red bands), Spearman correlation coefficient, and p-value indicated. **(B.)** Spatial plot of primary GBM sample annotated by GBM cluster and cell type identity. **(C.-D.)** Spatial plot as in B. annotating all human cells by their normalized expression of different genes representing different spatial and cellular state patterns: (C.) *TNC* and *CSPG4* (AC-/MES- and NPC-/OPC-like marker genes, respectively), and (D.) *PLP1* (oligodendrocytes), *PDGFRB* (endothelial cells), *CD74* (TAMs - 1), *FTL* (TAMs - 2), and *SPP1* (TAMS - 3). Expression values were clipped at a per gene basis. **(E.-F.)** Violin plots displaying ecEGFR copy number (E.) and EGFR introns (F.) across GBM clusters. **(G.)** Each cell’s PC1 score (X-axis) and EGFR copy number (Y axis) were plotted for all primary GBM cells and were annotated by their GBM cluster identity. **(H.)** Similar to Fig.7G (see methods), rank-rank plot showing each gene’s ranked correlation with ecEGFR copy number and ranked PC1 loading score. The top five genes for the positive (“Top 5”) and negative (“Bottom 5”) PC1 scores for primary GBM were annotated.

**Extended Data Figure 19.**
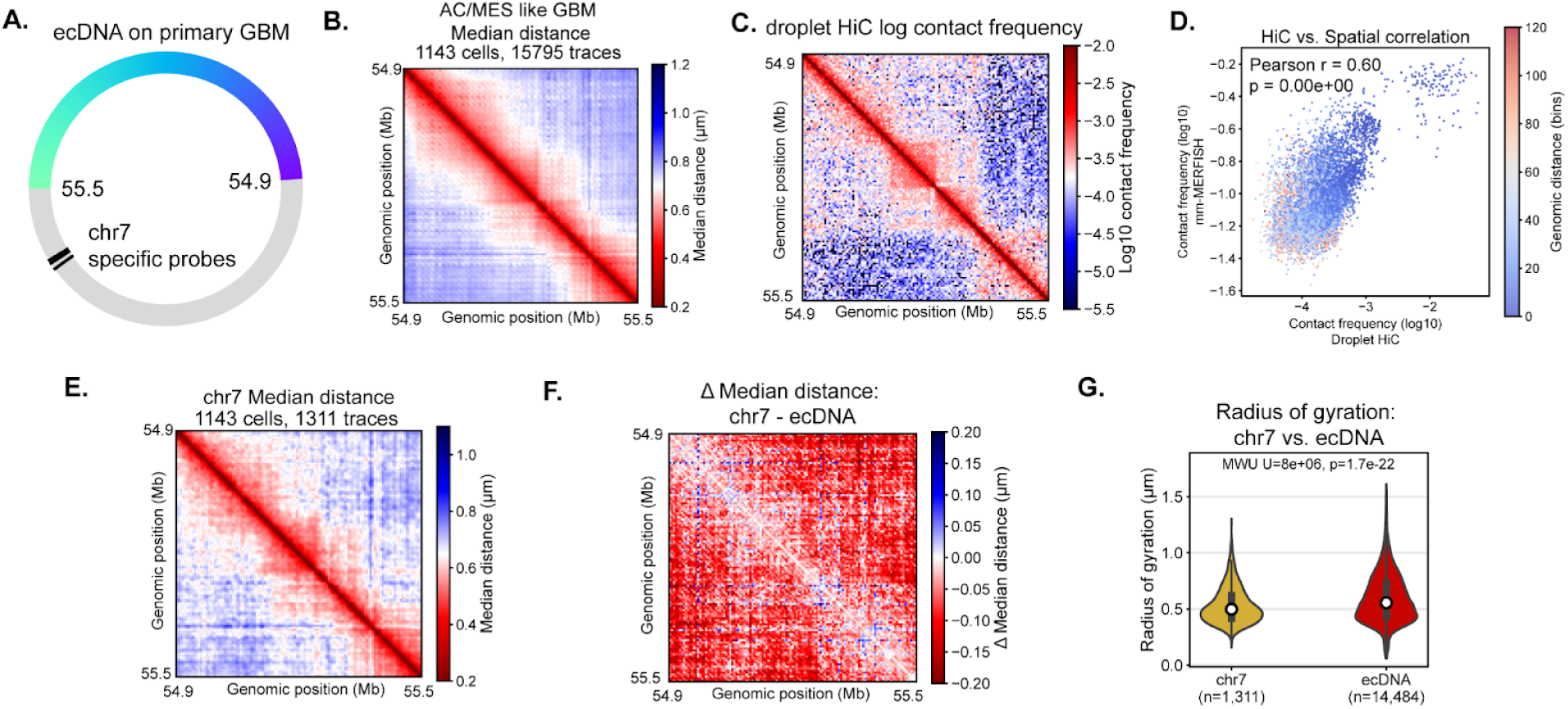
ecDNA is more expanded compared to its chromosomal counterpart in patient-derived GBM tissue. **(A.)** Diagram of the genomic regions of ecDNA in the primary GBM that overlaps with the ecEGFR in GBM39. Black lines denote the segments that are used to distinguish the chr7 EGFR locus from the ecDNA. **(B.)** Median distance between all 5-Kb segments across all traces in the AC/MES like GBM cells. **(C.)** Droplet HiC log10 contact frequency on the corresponding EGFR locus from the same patient sample. **(D.)** Correlation between HiC contact frequency and mm-MERFISH derived contact frequency. Contact is considered if the pairwise distance between loci is smaller than 250 nm. **(E.)** Median distance matrix of the chr7 EGFR locus in AC-/MES-like GBM cells. **(F.)** Difference in median physical distance between chr7 EGFR locus and ecDNA. **(G.)** Violin plots of the radius of gyration of chr7 EGFR locus and ecDNA. A two-sided Mann Whitney U test was used to test for statistical significance.

**Extended Data Figure 20:**
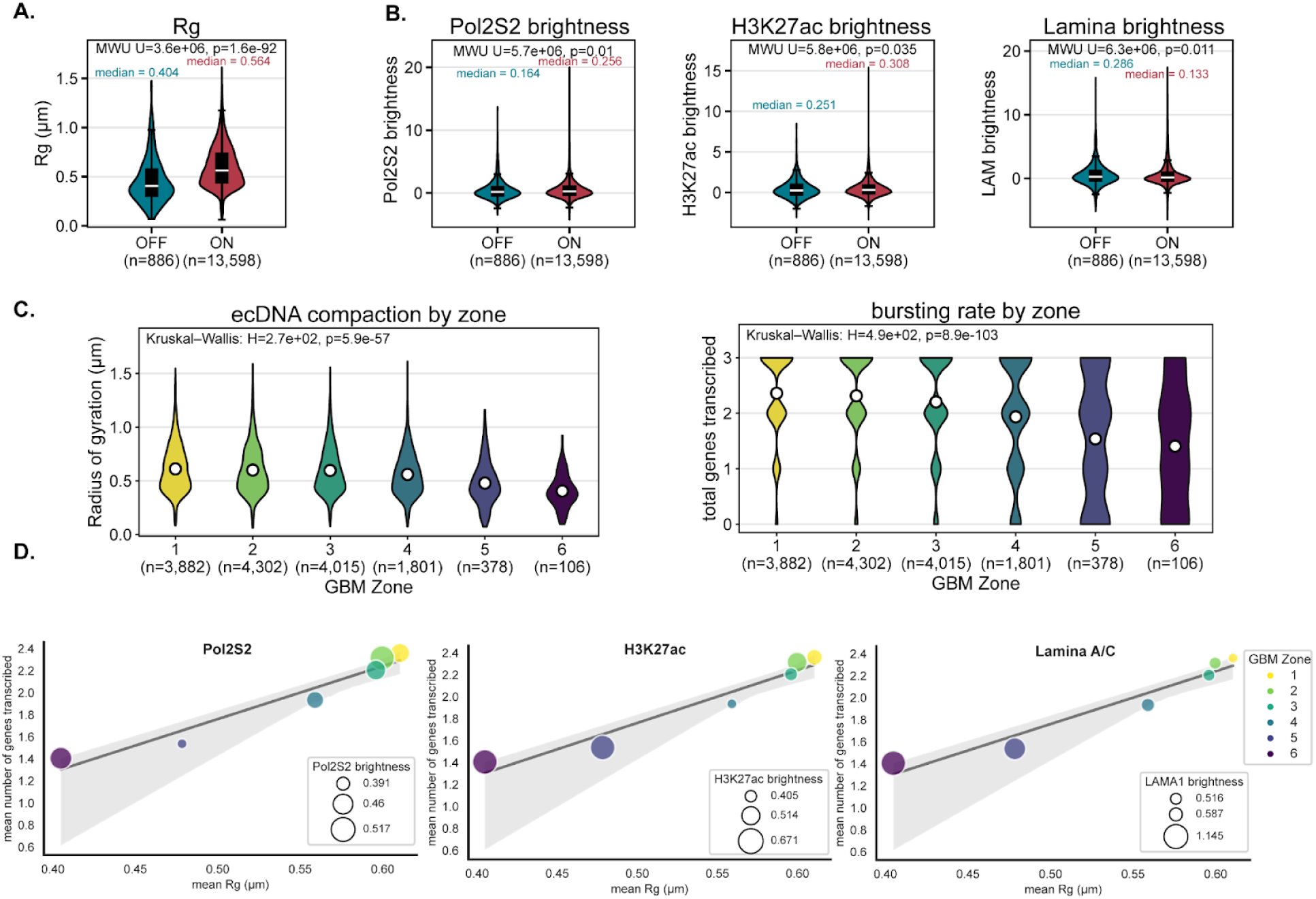
mm-MERFISH resolves the single-molecule chromatin structure and transcriptional activity across the architecture of primary GBM. **(A.-B.)** Violin plot of radius of gyration (A.) and regulatory machinery association (B.) across ecDNA molecules that are not actively transcribing (OFF, teal) and actively transcribing (red). A two-sided Mann Whitney U test was used to test for statistical significance. **(C.)** Violin plots of radius of gyration (left) and total number of genes transcribed (right) of all ecDNA in AC/MES GBM like cells across GBM zones. White dots mark the mean in each group. **(D.)** Scatter plot of the mean Rg (X-axis) and mean genes transcribed (Y-axis) of all EGFR locus traces from AC-/MES-like cells across each GBM zone. The “Mean Genes Transcribed” metric was derived from three introns captured within these traces: *EGFR*, *EGFR-AS1*, and *LANCL2*. Each point is annotated by color based on their GBM zone and by size based on their mean normalized Pol2S2 brightness (left), H3K27ac brightness (middle) and Lamina brightness (right).

## Supplementary Tables

**Supplementary Table 1**: Sequences of mm-MERFISH probes

**Supplementary Table 2**: Sequences of mm-MERFISH associated adaptors, primers, and read out

**Supplementary Table 3**: Statistical output for classifying the presence of ecDNA from 10X-Visium data

